# The longitudinal behavioral effects of acute exposure to galactic cosmic radiation in female C57BL/6J mice: implications for deep space missions, female crews, and potential antioxidant countermeasures

**DOI:** 10.1101/2024.04.12.588768

**Authors:** S Yun, FC Kiffer, GL Bancroft, CS Guzman, I Soler, HA Haas, R Shi, R Patel, J Lara-Jiménez, PL Kumar, FH Tran, KJ Ahn, Y Rong, K Luitel, JW Shay, AJ Eisch

## Abstract

Galactic cosmic radiation (GCR) is an unavoidable risk to astronauts that may affect mission success. Male rodents exposed to 33-beam-GCR (33-GCR) show short-term cognitive deficits but reports on female rodents and long-term assessment is lacking. Here we asked: What are the longitudinal behavioral effects of 33-GCR on female mice? Also, can an antioxidant/anti-inflammatory compound mitigate the impact of 33-GCR? Mature (6-month-old) C57BL/6J female mice received the antioxidant CDDO-EA (400 µg/g of food) or a control diet (vehicle, Veh) for 5 days and either Sham-irradiation (IRR) or whole-body 33-GCR (0.75Gy) on the 4th day. Three-months post-IRR, mice underwent two touchscreen-platform tests: 1) location discrimination reversal (which tests behavior pattern separation and cognitive flexibility, two abilities reliant on the dentate gyrus) and 2) stimulus-response learning/extinction. Mice then underwent arena-based behavior tests (e.g. open field, 3-chamber social interaction). At the experiment end (14.25-month post-IRR), neurogenesis was assessed (doublecortin-immunoreactive [DCX+] dentate gyrus neurons). Female mice exposed to Veh/Sham vs. Veh/33-GCR had similar pattern separation (% correct to 1st reversal). There were two effects of diet: CDDO-EA/Sham and CDDO-EA/33-GCR mice had better pattern separation vs. their respective control groups (Veh/Sham, Veh/33-GCR), and CDDO-EA/33-GCR mice had better cognitive flexibility (reversal number) vs. Veh/33-GCR mice. Notably, one radiation effect/CDDO-EA countereffect also emerged: Veh/33-GCR mice had worse stimulus-response learning (days to completion) vs. all other groups, including CDDO-EA/33-GCR mice. In general, all mice show normal anxiety-like behavior, exploration, and habituation to novel environments. There was also a change in neurogenesis: Veh/33-GCR mice had fewer DCX+ dentate gyrus immature neurons vs. Veh/Sham mice. Our study implies space radiation is a risk to a female crew’s longitudinal mission-relevant cognitive processes and CDDO-EA is a potential dietary countermeasure for space-radiation CNS risks.

## 1. INTRODUCTION

The interplanetary space between Earth and Mars consists of a complex field of dynamic particle radiation, derived primarily from galactic cosmic rays (GCR). GCR can penetrate current spacecraft shielding, and thus the risk of GCR to humans on interplanetary missions merits thorough evaluation (Nelson 2016). GCR risk estimates rely on animal models exposed to accelerated particles at energies encountered in space (Schimmerling 2016). However, particle accelerators have been limited in the number of particle types and energies that can be generated in a reasonable time-frame (Simonsen *et al*. 2020). Therefore, most space radiation risk estimates have relied on studies exposing animals to monoenergetic single particle types whose energies, on average, are higher than the true multi-energetic, multi-particle spectrum GCR encountered in space. Fortunately, GCR production capabilities at Brookhaven National Laboratories are now sufficiently advanced that scientists can now assess if exposure to whole-body multi-particle irradiation (IRR) — including 33-ion GCR (33-GCR) — alters cognition when compared to sham-IRR rodents. Taken together, studies show single-ion, multi-ion, and even 33-GCR disturb a range of rodent central nervous system (CNS) functions, including memory, pattern separation, anxiety, vigilance, social novelty, and motor control (Kiffer *et al*. 2019b; Cekanaviciute *et al*. 2018; Clément *et al*. 2020). However, narrowing these studies to just those that use multi-particle GCR (multi-ion, up to 33-GCR) highlights some contradictory results (Raber *et al*. 2016; Kiffer *et al*. 2018; Krukowski *et al*. 2018a; Raber *et al*. 2019; Kiffer *et al*. 2020; Raber *et al*. 2020; Klein *et al*. 2021; Boutros *et al*. 2021; Krukowski *et al*. 2021; Raber *et al*. 2021; Schaeffer *et al*. 2022; Britten *et al*. 2022; Puukila *et al*. 2023; Alaghband *et al*. 2023; Kiffer *et al*. 2021). In addition, while most work has been performed in just male rodents, some of these studies show sex-specific outcomes of multi-ion whole-body exposure. Thus, the impact of multi-ion GCR, particularly the Mars-relevant 33-GCR, on higher cognitive function in female mice warrants more research. In addition, while a few papers have now looked at the behavioral impact of 33-GCR up to 6-months (mon) post-IRR (Alaghband *et al*. 2023; Desai *et al*. 2023), more work is needed to assess the even longer-term impact of 33-GCR, particularly given concerns about astronaut morbidity.

In addition to studying the sex-specific and long-term impact of GCR exposure, there is a need to either prevent or ameliorate the CNS effects of GCR. Prevention strategies — such as improved or additional spacecraft shielding — are challenging. In fact, additional shielding may paradoxically lead to absorption of higher GCR dose due to particle fragmentation, back-scatter, and linear energy transfer modulation (Warden and Bayazitoglu 2021; Naito *et al*. 2021). Because of this, amelioration strategies — such as the use of pharmacological countermeasures — have received significant attention (Singh and Seed 2022; Dynan *et al*. 2022; DiCarlo *et al*. 2022). Medical countermeasures for space radiation exposure are currently evaluated by lifetime carcinogenic mortality (Werneth *et al*. 2022); no guidelines or modeling yet exist for the mitigation of acute, in-flight CNS or operational performance risks for flight crews (Nelson *et al*. 2016). However, the ideal pharmacological countermeasure for in-flight use would have the following characteristics: 1) high distribution to target organs; 2) high biological half-life; 3) amenable to non-invasive administration; 4) be radioprotective and mitigative, acting via multiple points of radiogenic damage, including DNA damage, oxidative stress, and inflammation, and promote tissue repair (Pariset *et al*. 2021). With these criteria in mind, 2-cyano-3, 12-dioxooleana-1, 9-dien-28-oic acid-ethylamide (CDDO-EA) warrants careful examination as a potential countermeasure. CDDO-EA is a successful pharmacotherapeutic in animal models for diseases that target the CNS and other organs, acting by promoting endogenous antioxidant pathways, protecting and promoting DNA repair, reducing inflammation, and promoting cellular repair (Wible *et al*. 2018; Mathis and Cui 2016; Wang *et al*. 2014; Neymotin *et al*. 2011; Luitel *et al*. 2020; Stack *et al*. 2010; Lei *et al*. 2020; Crowley *et al*. 2017). The human version of CDDO-EA known as omaveloxolone (Skyclarys, CDDO-ME) was recently approved by the FDA for the treatment of Friedreich’s ataxia, a neuromuscular disorder (Biogen.com). Recently, we reported that male mice exposed 6-mon prior to 33-GCR and concomitantly given CDDO-EA had worse social novelty than sham-irradiated mice, and that CDDO-EA was not an effective countermeasure (Kiffer *et al*. 2021). However, the impact of 33-GCR and the countermeasure potential of CDDO-EA on female mice and their CNS function are unknown.

There were three goals to this study. First, we wanted to define the impact of 33-GCR on female cognitive tests with relevance to mission success. We employed the translationally-relevant touchscreen platform to probe multiple domains of cognitive performance in female mice previously exposed to 33-GCR, the potential countermeasure CDDO-EA, and a combination of IRR and diet. Our choice of touchscreen task was guided by past work showing IRR changed function linked to many brain regions. For example, hippocampal dysfunction is seen in a sex– and task-specific manner up to 8-mon post-^56^Fe IRR (Whoolery *et al*. 2020; Soler *et al*. 2021) and 2-3.5 mon post-33-GCR (Alaghband *et al*. 2023). Notably absent from this literature is the impact of 33-GCR on behavioral pattern separation in female mice, an ability that is assessed by the touchscreen test location discrimination reversal (LDR) (Oomen *et al*. 2013; McTighe *et al*. 2009). Dysfunction in the prefrontal cortex (PFC) is also evident in the literature, including IRR-induced dysfunction in cognitive flexibility in female and male rodents (Whoolery *et al*. 2020; Soler *et al*. 2021; Shukitt-Hale *et al*. 2000; Villasana *et al*. 2013; Britten *et al*. 2014; Bellone *et al*. 2015; Hadley *et al*. 2015; Rudobeck *et al*. 2017; Britten *et al*. 2022; Burket *et al*. 2021; Britten *et al*. 2023; Desai *et al*. 2023). However, there is no work examining the impact of 33-GCR on the performance in two PFC-involved touchscreen tasks: LDR (Swan *et al*. 2014; Soler *et al*. 2021; Yun *et al*. 2023), specifically the reversal part which is reliant on PFC, and extinction of simple stimulus-response learning (Mar *et al*. 2013). Notably, the acquisition phase of the simple stimulus-response learning task relies on two other brain regions: the striatum and amygdala (Fernando *et al*. 2013). Performance on the simple stimulus-response learning task has not yet been assessed after exposure to 33-GCR. It would be important to do this since older work shows single-particle exposure disrupts amygdala and striatal functional integrity (Kiffer *et al*. 2019b). There is an additional benefit to reaching this first goal via touchscreen testing: Traditional studies in radiation have, with a few exceptions, relied on arena-based tests that involve animals freely-behaving in open areas. These arena-based tests provide excellent insight into behavior, but the open spaces also are anxiogenic in prey species, such as rodents. Arena-based also typically involve only a few trials, limiting temporal resolution of behavioral data captured. The touchscreen-based platform involves sound-isolated, confined spaces with large, versatile visual stimuli and the ability to have appetitive rewards for correct stimulus interaction, prompting repeated, high-resolution behavioral evaluations that are highly adaptive and can probe multiple brain systems. Thus, here we probe multiple brain circuit integrity by testing both on the touchscreen platform (via LDR and simple stimulus-response learning/extinction) and on a battery of arena-based tests. In addition to probing the function of multiple brain regions after 33-GCR exposure, our second goal was to assess the ability of CDDO-EA to prevent 33-GCR-induced changes in long-term behavior or cognition. Prior work has shown CDDO-EA given before, during, and after 33-GCR or sham IRR blunted GCR-induced changes in social behavior when assessed 6-mon later (Kiffer *et al*. 2021). However, the radioprotective impact of CDDO-EA in female mice has not yet been assessed. Our third and final goal was to evaluate behavior over many months and conclude with a meaningful assessment of a cellular index. To this end, once the 14-mon of behavior and cognitive testing was complete, we assessed hippocampal dentate gyrus neurogenesis in these female mice. Dentate gyrus neurogenesis is critical to a number of behavioral domains, ranging from memory to spatial learning to the social domain (Rivera *et al*. 2013; DeCarolis *et al*. 2014; Whoolery *et al*. 2017; Zanni *et al*. 2018; Whoolery *et al*. 2020). Decreased neurogenesis is seen after whole-body exposure to single particle types at Mars-mission-relative doses, and some, but certainly not all, work suggests that the number of dentate gyrus immature neurons correlates with hippocampal function, such as improved pattern separation.

With these four goals in mind, we set out to understand the potential risks of 33-GCR to the CNS. We exposed mature (6-mon-old) female C57BL/6J mice to the whole-body Mars mission-relevant dose of 75cGy. Over the subsequent 14.25 mon, we assessed spaceflight-relevant behavioral and cognitive domains using both operant touchscreen and arena-based testing. We also examined the cellular outcome of hippocampal neurogenesis in Sham vs 33-GCR mice. Subgroups of Sham and 33-GCR mice were given CDDO-EA or Veh chow during IRR, thus allowing us also to assess CDDO-EA as a candidate countermeasure in astronaut-aged female mice.

## 2. METHODS

### 2.1 Animals and housing

Six mon-old female C57BL/6J mice (Jackson Laboratory stock #000664, weanling mates, Bar Harbor, ME) were ordered and shipped directly to Brookhaven National Laboratories (BNL), specifically the Brookhaven Laboratory Animal Facility (BLAF). The day before irradiation (IRR), mice were transported in their same cages from the BLAF to the NSRL (NASA Space Radiation Laboratory) and were returned to BLAF the day after irradiation. At BLAF and NSRL, mice were housed 4/cage in a HEPA-filtered, closed airflow vivarium system under a 12:12 h (6am to 6pm light) light/dark cycle at 22°C, 30-70% humidity and given a standard rodent chow (LabDiet 5015 #0001328) and water *ad libitum*. Two days after irradiation, mice were transported to CHOP by ground shipment. At CHOP, mice were held in quarantine for 1.5 mon (6 weeks) with a 4% fenbendazole chow (vermicide). Behavioral testing began at 2.5 mon (10 weeks) post-IRR when mice were released from quarantine and were back on regular chow (LabDiet 5015). At CHOP, mice were housed in a HEPA-filtered, closed airflow vivarium system (Lab Products Inc., Enviro-Gard™ III, Seaford, DE) under a 12:12 h (6am to 6pm light) light/dark cycle at 20-23°C and 30-40% humidity) and given a standard rodent chow (LabDiet 5015 #0001328) and water *ad libitum*. At cage change, each cage was given a nestlet square; otherwise, no enrichment was provided. All care and procedures were approved by the Institutional Animal Care and Use Committees (IACUC) at BNL and CHOP and held in accordance with the Association for Assessment and Accreditation of Laboratory Animal Care (AAALAC) and National Institute of Health (NIH) guidelines for the care and use of laboratory animals. Our scientific reporting adheres to the ARRIVE 2.0 guidelines.

### 2.2 Diet

After 3 days of acclimation to the BNL BLAF, mice were split into 2 diet groups to receive either a vehicle chow (Veh: Purina Rodent Diet 5002, 12.5 g EtOH, 37.5 g Neobee Oil) or a CDDO-EA chow (400 µg RTA 405 per 1g of Veh; Reata Pharmaceuticals, Irvine, TX) *ad libitum* for 5 days. Both the Veh and CDDO-EA formulations were prepared by Purina Mills, LLC. The irradiation (next section) took place on Day 4 of the 5-day diet.

### 2.3 Irradiation

On Day 4 of the Veh or CDDO-EA diet, mice in both groups were pseudorandomly subdivided into Sham or 33-GCR groups (n=13-16 per Diet/Radiation group; **Fig. 1**). The irradiation with 33-GCR took place during the BNL 19A campaign, as previously described (Kiffer *et al*. 2021). In brief, mice were placed in well-ventilated 10×10×4.5cm holders in pairs with cagemates. They were then exposed to 75cGy of NASA’s whole-body 33-beam 33-GCR in a 60×60cm beam spread. The 75cGy was delivered over ∼1.5 hours, and beam uniformity and dosimetry were evaluated and monitored by the staff at NSRL. The remaining mice that received Sham exposure were also paired with cagemates in the holders for 1.5 hours but were not placed on the beam line. The four experimental groups that resulted were Veh/Sham, Veh/33-GCR, CDDO-EA/Sham, and CDDO-EA/33-GCR.

**Figure 1.**
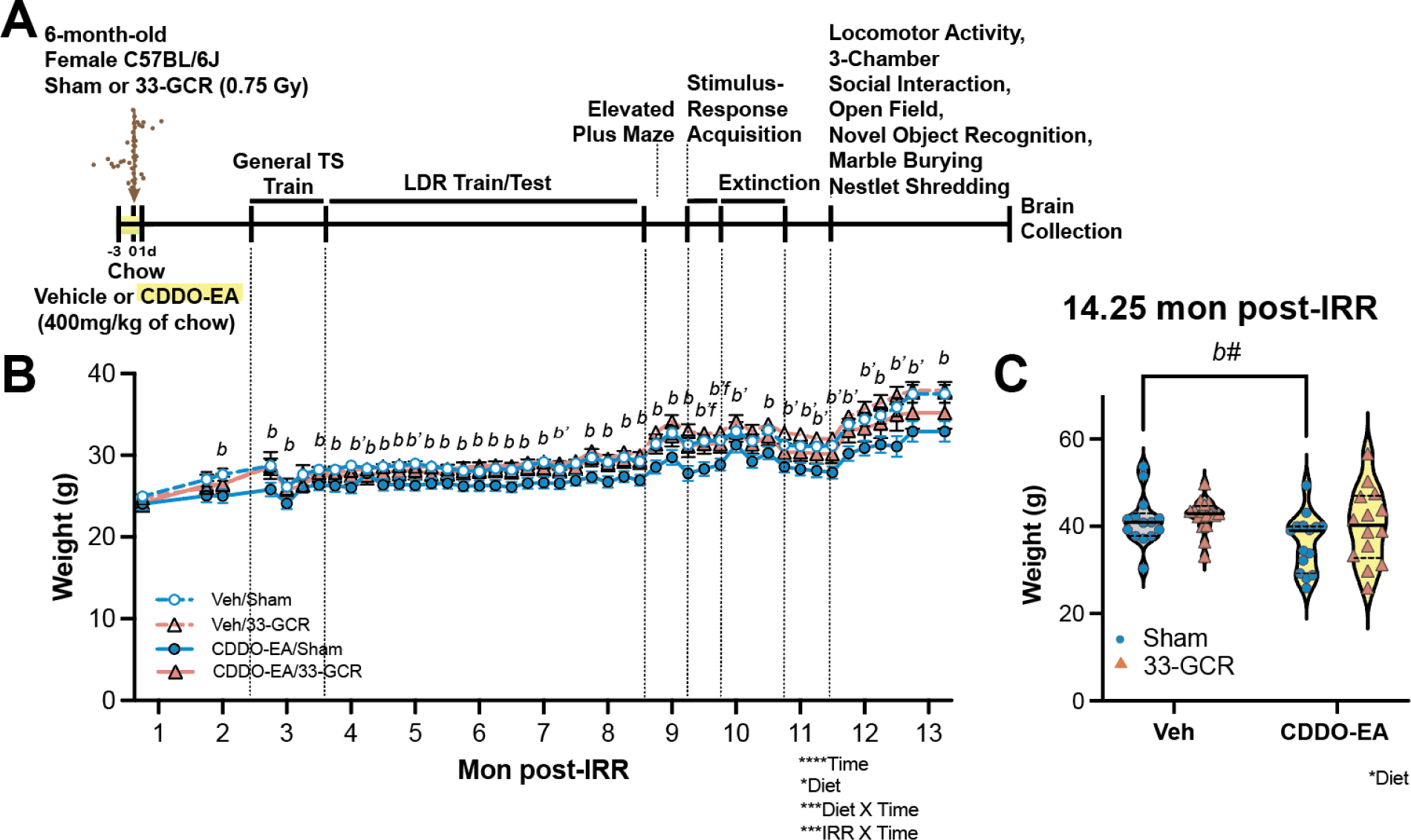
Female C57BL/6J mice given the antioxidant/anti-inflammatory agent CDDO-EA chow and exposed to Sham IRR weigh slightly less than Veh/Sham mice throughout the experiment. This lower weight is not seen in CDDO-EA/33-GCR mice. (**A**) <u>Timeline of experiment.</u> Female mice from Jackson Laboratory (#000664) were delivered to Brookhaven National Laboratory (BNL) at 6-months (mon) of age. After acclimation to the BNL animal housing facilities, mice were given CDDO-EA (400 mg/kg of food) in chow or vehicle chow (Veh) for five days. On Day 3 of CDDO-EA or Veh diet, mice were exposed to one of two IRR conditions: whole-body 33-GCR (0.75 Gy) or Sham IRR. Mice were then shipped to Children’s Hospital of Philadelphia Research Institute where they underwent general training on a touchscreen (TS) platform followed by a task to assess behavioral pattern separation, Location Discrimination Reversal (LDR) and a simple stimulus-response learning and extinction learning. Mice then went through a series of classical arena-based behavioral tests prior to brain collection at 14.25 mon. (**B**) <u>Body weight of mice over the course of experiment.</u> CDDO-EA/Sham mice showed lower weight relative to Veh/Sham mice throughout the experimental period (both before and after food restriction began), and at two time points CDDO-EA/33-GCR weighed 8-9.8% less than CDDO-EA/Sham. (**C**) <u>Weight at 14.25 mon.</u> When the experiment ended, body weight measures showed that CDDO-EA/Sham mice weighed 9.9% less vs Veh/Sham mice. Statistical analysis in B: Mixed effected three-way RM ANOVA (IRR X Diet X Time); in C: Two-way ANOVA (IRR X Diet). Main effect and/or interaction are denoted by *P<0.05, **P<0.01, ***P<0.001, ****P<0.0001; post-hoc analysis significance denotations between groups: Veh/Sham vs CDDO-EA/Sham, *b* P<0.05, *b’* P<0.01, *b#* = 0.065; CDDO-EA/Sham vs CDDO-EA/33-GCR, *f* P<0.05. 14-16 mice per group. Dotted vertical lines indicate the point on the behavioral timeline where weight was recorded in B and violin plots depict median (solid line) and 25% and 75% quartiles (dotted line) in C. Complete subject numbers and detailed data analysis are provided in **Supplemental Table 1** (**Supp. Table 1**).

### 2.4 General physiological measures

Mouse body weight data were regularly collected from their arrival at BNL at 6-mon of age, through behavioral testing, and until brain collection 14.25mo (57 weeks) post-IRR. “Survival” of mice was recorded throughout the experiment, with the survival graph reflecting mice that were lost due to possible diet and/or radiation effects; mice that were lost due to husbandry issues were not considered in the survival graph (see **Subject Number** section below). Alopecia (coat state) scores were recorded twice (10.5 mon or 42 weeks post-IRR and 13.5 mon or 54 weeks post-IRR) as general indication of stress or maladaptation to treatment (Bechard *et al*. 2011).

### 2.5 Behavioral testing

#### Overview of Behavioral Testing

Two and one-half months (10 weeks) post-IRR, mice began behavioral testing **(Fig. 1A)**. Mice first underwent <u>touchscreen training and testing</u> interspersed with and followed by <u>arena-based tests</u> to assess early to late effects of GCR on multiple behavioral domains, as described in separate sections below. Mice were weighed weekly to ensure that weights did not fall below 80% of their starting weight. Mice that exceeded this weight loss limit would be excluded from the experiment, however no mouse met this exclusion criterion.

Mice underwent <u>touchscreen training and testing</u> on the location discrimination reversal (LDR; program LD1 choice reversal v3; ABET II software, Cat #89546-6) from 2.5 to 8.6 mon post-IRR (general touchscreen training, location discrimination reversal training [LDR Train], and location discrimination reversal testing [LDR Test]) and then the Extinction training [Acq] and testing 9.25 to 10.8 mon post-IRR (Ext; ABET II Software, Cat #89547). During these time periods, mice were food restricted Monday through Friday, with food only provided ∼4 hours (hr) each day after touchscreen testing. Food was provided *ad libitum* from Friday afternoon through Sunday evening. Data were automatically recorded from the operant touchscreen platform.

Mice underwent <u>arena-based testing</u> from 8.6 to 8.9 mon post-IRR (elevated plus maze [EPM]) and from 10.8 to 12 mon post-IRR (locomotor activity, 3-chamber social interaction [3-CSI], open field [OF], novel object recognition [NOR], marble burying, nestlet shredding). During these time periods, mice were given unrestricted food access. For arena-based testing (with the exception of marble burying and nestlet shredding), data and video recordings were captured by a ceiling-mounted camera (Ace acA640–90gc, Basler) and tracking was performed by Ethovision XT (Noldus Information Technology). Nose point, center point, and tailbase were captured. Nose points were used for exploratory measures, whereas center points were used for gross locomotion. Marble burying was scored by the handler performing the experiment, and nestlet shredding was assessed by weighing pre– and post-test intact materials.

In all tests, the handler placed the test mouse in the testing apparatus and left the room for the duration of the trial. Between mice, behavioral equipment was disinfected and deodorized with 10% ethanol or 10% TB-10 (Birex) and allowed to dry. The four experimental groups were run through tests in a counterbalanced order to minimize the effect of testing order.

##### Touchscreen Training and Testing

Touchscreen testing was performed between 8am and 2pm daily (Mon-Fri). The rewarding appetitive stimulus (“reward”) used throughout training and testing was Strawberry Ensure^®^ Nutrition Shake (Abbott Laboratories).

#### General Touchscreen Training and Food Restriction Overview

Prior to behavioral testing in the Bussey-Saksida operant touchscreen platform, mice moved through several general training stages, as previously described (Soler *et al*. 2021; Whoolery *et al*. 2020). Specifically, general touchscreen training consists of six phases, two of which are set for one-day duration (<u>Habituation [Hab] 1</u> and <u>Hab 2</u>) and four of which are criterion-based (and thus have varied durations): <u>Initial Touch</u>, <u>Must Touch (MT)</u>, <u>Must Initiate</u>, and <u>Punish Incorrect (PI)</u>. <u>Hab 1</u> (1 day) began 2.5 mon post-IRR and consisted of 10 mins of free movement in the touchscreen chamber. A 2×6 window grid is provided, but no reward, tone, or house light is provided. This is the first time the mice are in the touchscreen chamber. Beginning 14h prior to Hab 2 and continuing for the duration of touchscreen experiments, mice were restricted to their standard chow beginning at 6pm each evening until 2pm the following day during touchscreen training or testing sessions. From Fri (2pm) to Sun (5pm), unrestricted food is provided. Mice were weighed weekly to ensure that weights did not fall below 80% of their starting weight. Mice that exceeded this weight loss limit would be excluded from the experiment, however no mouse met this exclusion criterion.

For <u>Hab 2</u> (1 day), mice are placed in the same touchscreen chambers for 20 min. The reward tray light is on, and an initial priming reward of 150uL is provided simultaneously with a tone (70dB, 500 Hz, 1s). After the mouse inserts and withdraws their head from the reward tray, the tray light turns off, there is a 10s delay, followed by reactivation of the tray light and tone, and 7uL of Ensure is dispensed into the reward tray.

For the subsequent <u>Initial Touch</u> phase (1-2 days, criteria 25 trials/30 min), mice begin with one of the bottom six grids pseudorandomly lit. The stimulus was not displayed in the same position more than 3 times in a row. Mice have 30s to touch the lit square. If the correct square is touched, 7uL of the rewarding stimulus is dispensed, the reward tray lights up and a tone is played. If, however, no touches or incorrect touches are made, the reward tray dispenses only 7uL of the reward stimulus, but still illuminates and plays a tone. Regardless of response, there is a 10s intertrial interval (ITI) where the reward tray light turns off. To complete this phase mice must complete 25 trials (regardless of touch or omission) in 30 mins. Mice who complete 25 trials are promptly removed from the arena and returned to their home cage.

Next, mice learn <u>MT</u> (criteria 25 trials/30 min), where they must touch the screen stimuli to receive the appetitive reward. This phase is identical to the Initial Touch phase but requires mice to touch the correct lit square on the screen to receive the reward. Correct touches result in only 7uL of the reward in conjunction with the reward tray light and tone. Incorrect touches (touches to unlit squares) yield no reward, light or sound cues.

Once MT criteria are reached, mice then advance to the <u>Must Initiate</u> phase (criteria 25 trials/30 min), where trials do not begin until a mouse inserts her head into the reward tray. Reward rules and completion criteria are the same as in the MT, but once rewards are collected, mice must reinsert their head into the reward tray to initiate the following trial.

Once mice reach Must Initiate criteria, they move on to the final training phase, <u>PI</u> (criteria: total trial counter 30, consecutive 76% correct for the first day and 80% correct for the 2nd day in a row). This phase “punishes” incorrect touches (touching an unlit grid square) by turning on the chamber light for 5s. Then a correction trial occurs, where the same rewarding stimulus square as the previous trial is once again lit. A correction trial is repeated until the mouse touches the rewarding stimulus. Correction trials do not count in the total trial counter in regard to completion criteria.

#### Location Discrimination Reversal training (LDR Train) and testing (LDR Test)

After general touchscreen training, mice then proceed to <u>LDR training</u> (LDR Train [50 Max trials/30min] and testing (LDR Test [81 Max trials, 8 Max reversal], program LD1 choice reversal v3; ABET II software, Cat #89546-6). In the LDR Train phase, mice learn that touching one of two squares separated by an “intermediate” distance results in an appetitive reward. Faced with the same 2×6 grid used for general touchscreen testing, mice are presented with one of two lit squares, either the 2nd or 5th square in the bottom row. Only one of the squares is the rewarded stimulus (S+) at a given time during the daily 30 min session. Touching the correct side leads to the same rewarding conditions as during training (reward tray light, tone, and 7uL of the reward), whereas incorrect touches (touches to S-) yield no reward and a time out period w/ house light on. After reward collection or a time-out end, ITI (10s) will be followed before a new trial starts. The following trial will start when animals poke their noses in the reward tray. The following day, the other lit square becomes S+ based on the previous day’s final trial, and continues alternating until each mouse reaches 50 trials (regardless of accuracy) within 30 min. Reversals occur when the S+ is correctly chosen in 7 of 8 consecutive trials, and the opposite side of the screen becomes S+ within the same 30 min testing session. Location discrimination reversal training completion occurs when mice reach at least 2 reversals in 2 out of 3 consecutive days. Mice then move onto the <u>LDR Test</u> phase where there are two difficulty settings: ‘Large Separation’, where the S+ and S-stimuli are the most lateral squares (7th and 12th windows on the bottom row of a 2 x 6 grid) and ‘Small Separation’ where the S+ and S-stimuli are the center-most squares (9th and 10th windows of the bottom row on the same grid window). The difficulty settings at the start are assigned pseudorandomly to each mouse and are counterbalanced within-group. Similar to the training phase, reversals occur once 7/8 correct consecutive S+ response are made, the S+ side is switched to the opposite side of the screen. Separation difficulties alternate every 2 days, and sessions end when mice complete 81 trials (regardless of accuracy) or 30 mins passed. This test phase ends after 20 sessions (10 in each Large and Small separation) are completed. These LDR Train data were analyzed: % subject to reach criteria, days to completion, the Last Day performance measurement (completed trial #, session length, percent correct). These LDR Test discrimination learning (Large separation) and pattern separation (Small separation) data were analyzed: percent correct to 1st reversal over all 5 blocks (Last Day of each block), and percent correct, trial number, and session length to 1st reversal in the first block (Block 1) and last block (Block 5). These LDR Test reversal learning/cognitive flexibility data were analyzed: number of reversals achieved in Large and Small separations in the Last Day of Block 5 as well as in the Last Day of Blocks 1-5.

#### Stimulus-Response Acquisition (Acq) and Extinction (Ext) Learning

After the completion of LDR Test at 8.6 weeks post-IRR, all mice received normal chow *ad libitum* for two weeks. At the end of one week, mice first underwent elevated plus maze (see Arena-based Testing, below). At 9.25 weeks post-IRR, mice were placed back into the touchscreen operant chamber with a 1×3 grid for a reintroduction period (10 min, 1 day) which used the Hab 1 program. That evening and 14h prior to the next touchscreen session (simple stimulus-response acquisition, or Acq), mice were again placed under food restriction.

Testing for <u>simple stimulus-response acquisition (Acq) and extinction (Ext) learning</u> in an operant touchscreen paradigm began the next day and used the programs Extinction 1 and 2, respectively, in the ABET II Software (Cat #89547). For both Acq and Ext, mice are faced with a 1×3 grid with the middle-lit rectangle as the S+. <u>Acq:</u> Acq sessions occur daily. As the mouse is placed into the chamber, the reward tray is illuminated and a reward is provided. Once the mouse nose pokes the tray, S+ appears in the middle of the touchscreen. Once touched, a reinforcing appetitive reward is delivered and a tone is played. Touches to either of the two lateral blank rectangles resulted in no reward and no tone. The mouse collects the reward and an intertrial interval (ITI) begins. Then the reward tray is illuminated again to begin the next trial. Mice were tested daily and sessions ended once mice completed 30 trials by touching the conditioned stimulus 30 times or after 30 mins has elapsed. Criteria for Acq of a simple stimulus-response include completion of 30 trials in <15 min in four consecutive days. Although each Acq session is 30-min in length, only the first 15-min are considered for completion criteria in order to ensure that subsequent Ext learning will occur (Mar *et al*. 2013; Soler *et al*. 2021; Cotter *et al*. 2023; Bussey *et al*. 2012). Once completed, each mouse moves directly to the Ext learning sessions. <u>Ext</u>: Ext sessions occur daily and begin with an initial 5-sec interval where the screen is blank. The stimulus then appears for ≤10 sec. If touched, the stimulus disappears, no reward is provided, but the food hopper is illuminated and the tone still plays. Touches to the blank parts of the screen result in no response from the apparatus and the 10-sec timer is unchanged. Regardless of omission (defined as no touch as all to screen), blank touch, or stimulus touch, a 10-sec interval of no stimulus presentation occurs between trials, but the hopper remains illuminated, and entries to the hopper turn the light off until the stimulus is touched in the following trial. Daily sessions consist of 30 trials. Mice complete the Ext once ≥24 trials are successfully omitted across 3/4 consecutive days. Measures collected for Hab 1 include beam breaks (total from reward tray, two side bars, and screen beam), for Acq include days to completion, session length, correct touch latency, reward collection latency, and for Ext include days to completion, session length, and response number. Acq performance over time was analyzed on Day 1, Day 4, and Last Day; Day 4 was selected since the Acq criteria requires completion of 30 trials in <15 min on four consecutive days. Ext performance over time was analyzed on Day 1, Day 8, and Last Day; Day 8 was selected since the average group Ext completion was 8 days.

##### Arena-based testing

For at least two weeks before all arena-based tests, mice had *ad libitum* access to food and water.

#### Elevated-Plus Maze (EPM)

Behavior in the <u>EPM</u> (Harvard Apparatus, #760075) was assessed 8.75 mon post-IRR **(Fig. 1A)**. Mice were placed in the center of the arena pseudorandomly facing one of two open arms (43×33×6cm, height × arm length × arm width) and freely allowed to explore the arena for a single 5-min trial. EPM was run under ∼25 lux (red light) and from 12:55 to 18:33. In Ethovision, arena areas were defined as open arms, closed arms, and the center zone between the four arms, and the center point of each animal was used for tracking. Measures collected for EPM include duration in open and closed arms and distance traveled in open and closed arms. From these measures, an exploration index was calculated:

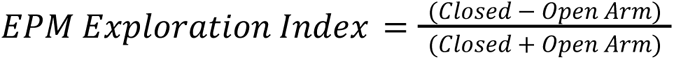

#### Gross Locomotor Activity in a novel environment

Gross <u>locomotor activity</u> was examined 11.5 mon post-IRR **(Fig. 1A)** after Acq and Ext. This test differed from other assessments of locomotion in this experimental timeline in that the mice were placed in a sound-isolated chamber to allow exploration of this novel environment and thus decrease anxiety-like behavior in subsequent 3-chamber social interaction. The activity chambers used were from Med Associates Inc. (#ENV-510, 27×27×20cm) and mice underwent a single 30-min session in ∼32 lux (white light) from 09:00 to 16:00. Measures collected by an array of infrared sensors include total distance moved, mean velocity, ambulatory time, ambulatory episodes, rearing time, and number of rearings.

#### 3-Chamber Social Interaction Test (3-CSI)

The <u>3-CSI</u> test was run ∼11.75 mon post-IRR **(Fig. 1A)** as a one-day, three-trial test where mice are placed in a bespoke three-chamber arena (Kiffer *et al*. 2021) to assess exploration during habituation, sociability, and social novelty. <u>Habituation trial</u>: Mice are placed in the empty, novel 3-CSI testing arena to assess inherent bias in the novel testing environment which might entice a mouse to spend more time in the left, middle, or right chamber. There are no objects/enclosures inside the 3-CSI chamber during habituation. <u>Sociability trial</u>: A novel conspecific mouse (“Stranger 1”) inside a small, perforated enclosure was pseudorandomly placed in the left or right chamber. In the opposite chamber, an empty perforated enclosure (“Empty”) is placed. <u>Social novelty trial:</u> The enclosure with Stranger 1 (a mouse which is now familiar) was placed back in the same left or right chamber and a second age– and sex-matched stranger conspecific (“Stranger 2”) was placed in a perforated enclosure in the opposite chamber. Each of the three trials (habituation, sociability, social novelty) were 10 min each and separated by 1 min. Experimenter gloves were switched frequently throughout 3-CSI and the arena and enclosures were cleaned with MB-10 and allowed to dry in-between test mice. Six age-matched C57BL/6J female stranger mice were used as the social stimuli pseudorandomly such that they were never used in two consecutive tests to minimize stress. All trials of 3-CSI were run under ∼25 lux (red light) and performed from 09:30 to 20:00. The cage tested first was chosen randomly each day, and cages alternated groups throughout the rest of that day. Ethovision arena dimensions and exploration areas are designated as previously described (Kiffer *et al*. 2021). For analysis of all three trials, arena boundaries were drawn for each chamber (left, middle, right). For analysis of social novelty, a “sniff zone” was designated in the left and right chamber and defined as a 1.5cm radius around the enclosure (which were either empty or held stranger mice). Measures collected for habituation include time spent in the left and right chamber. Measures collected for sociability include time spent in the social stimulus-containing chamber (Stranger 1) and the empty enclosure chamber (Empty). Measures collected for social novelty include time in the sniff zone around the familiar stimulus (Stranger 1) and around the novel social stimulus (Stranger 2).

#### Open Field (OF) in novel environment

<u>OF</u> was run between 12 and 13 mon post-IRR **(Fig. 1A)**. Two days of OF were performed (Day 1, Day 2) on subsequent days in order to capture exploration and locomotion in a novel environment (Day 1) and in a now-familiar environment (Day 2). Mice were placed in the center of a novel OF arena and allowed free exploration for one 10-min session for Day 1 and Day 2. OF was run under ∼25 lux (red light). OF Day 1 was run from 11:00 to 17:30, and OF Day 2 was run from 14:00 to 18:30. Exploration zones defined in Ethovision include the four 5×5cm corners (Corner) and the 20×20cm arena center (Center). OF measures collected include distance traveled, time in Center, and Exploration Index:

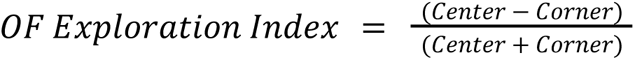

Day 1 and Day 2 metrics are shown and analyzed separately as they reflect different measures. As a measure of habituation learning, data are also presented comparing the distance traveled, time in Center, and Exploration Index of Day 1 vs Day 2.

#### Novel Object Recognition (NOR)

The <u>NOR</u> test was performed immediately after Day 2 of OF between 12 and 13 mon post-IRR **(Fig. 1A)**. NOR is a two-day test, but as it is run in the same arena as OF. Therefore, OF Day 1 and Day 2 served as two habituation trials for NOR, and the two NOR days can be considered Day 3 (familiarization with objects) and Day 4 (test day). On Day 3, two identical objects were placed in the center of the OF arena and mice freely explored for 10 min. The two identical objects were 50 mL blue-capped centrifuge tubes filled with blue nitrile gloves and water. On Day 4, one of the two objects was replaced with a novel object, a 200mL polycarbonate cell culture flask filled with light white aquarium pebbles, and the mouse explored for another 10 min. NOR was run under ∼25 lux (red light) and from 13:30 to 19:00. The arena, object dimensions, and exploration boundaries used for analysis are as previously described (Kiffer *et al*. 2021). Objects used were deliberately heavy to prevent mice from climbing on or pushing the objects and were made of common laboratory equipment to enable reproducibility. Measures collected for NOR include object exploration time on Day 4 (Familiar and Novel). From these data, object discrimination index was calculated as follows:

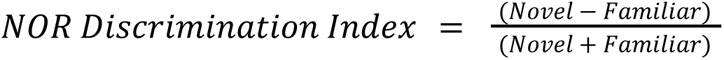

#### Marble burying in a home cage

<u>Marble burying</u> activity was measured between 12 and 13 mon post-IRR **(Fig. 1A)** as a measure of repetitive, compulsive-like behavior independent of anxiety or exploratory behaviors (Thomas *et al*. 2009; Luigjes *et al*. 2019). We used standard glass marbles (Ø = 15mm) placed upon 3cm of standard rodent hardwood bedding (Beta Chip, Northeastern Products Corp.) in a 20cm×35cm×15cm arena. Mice were allowed to freely interact with marbles in a single 20-min trial. At the end of the trial, mice were gently picked up and returned to their home cages. Marble burying was run in ∼25 lux (red light) and from 13:30 to 17:00. Measures collected for marble burying include the number of marbles buried which was evaluated by the experimenter. Marbles that were approximately ≤ 25% visible were considered buried.

#### Nestlet shredding in a home cage

<u>Nestlet shredding</u> was measured between 12 and 13 mon post-IRR **(Fig. 1A)** as a complementary method of measuring compulsive-like behaviors (Jeyabalan *et al*. 2022). At 20:00, mice were singly housed into identical cages with standard chow and water *ad libitum*. Mice were given pre-weighed nestlets (Animal Specialties and Provisions, ANNES) and no additional enrichment. Cages remained on the rack in the housing room overnight. Twelve hours later, mice were placed back into their homecage. Measures collected for nestlet shredding include the weight of the intact portions of the nestlets, with intact portions operationally defined by the experimenter.

### 2.6 Tissue Collection and Processing

Approximately 1 mon after the completion of behavioral tests (14.25 mon post-IRR), cages of mice were brought one at a time into the procedure room where mice were decapitated with 1min of being brought into the room. Brains were promptly dissected and immersion fixed in 4% paraformaldehyde. The following day, paraformaldehyde solution was replaced with fresh solution, and the day after brains were immersed in a 30% sucrose, 0.01% NaN_3_ solution. Brains were stored at 4°C until sectioning. Brains were sectioned (40um) in the coronal plane on a freezing microtome in preparation for stereological assessment of cells throughout the hippocampus, as previously reported (Lagace *et al*. 2010). Sections were stored in 1xPBS 0.01% NaN_3_ solution, and stored at 4°C. Prior to staining, whole hippocampus sections sampled at every 360μm (1:9 series) were mounted on superfrost-plus slides and dried at room temperature overnight.

### 2.7 Immunohistochemistry (IHC)

A random selection of mice that went through behavior were selected for DCX IHC (n=11/group). Slide-mounted IHC was performed as previously described (Yun *et al*. 2023). In brief, slide-mounted sections were placed into near-boiling citric acid (pH 6.0) for antigen unmasking. Endogenous peroxidases were inhibited via incubation in 0.3% hydrogen peroxide, and nuclear permeabilization was achieved by incubation in 0.1% Trypsin, 0.1M Tris, 0.1% calcium chloride solution. DNA denaturation was performed by incubation in 2N HCl in 1xPBS, and a blocking buffer of 3% normal donkey serum in 0.3% Triton X in PBS was used to block nonspecific binding. Sections were incubated in goat-α-doublecortin (DCX; 1:8,000, Santa Cruz Biotechnology) in 3% normal donkey serum and 0.3% Tween-20 overnight. After several 1xPBS washes, sections were then incubated in the secondary antibody (biotinylated-donkey-α-goat-IgG, 1:200, Jackson ImmunoResearch Laboratories Inc.) for 1h. After another series of 1xPBS washes, the signal was amplified and HRP-tagged via incubation in avidin-biotin complex (ABC Elite, Vector Laboratories), and then the HRP signal was colorimetrically visualized by diaminobenzidine (Pierce Scientific). Nuclear counterstaining was achieved by incubation in Fast Red (Vector Laboratories). Sections were dehydrated by incubation in a series of increasingly concentrated ethanol washes followed by delipidization by incubation in Citrasolv (Decon Laboratories). Coverslipping was done with DPX (Electron Microscopy Services) and 24×60 mm coverglass (VWR).

### 2.8 Stereological Cell Counts

DCX-immunoreactive (DCX+) cells in the subgranular zone (SGZ) of the dentate gyrus granule cell layer (GCL) were quantified in one hemisphere. An observer blind to experimental conditions counted all SGZ DCX+ cells by using an Olympus^®^ BX-51 brightfield microscope with 400x magnification and examining the entire focal (Z) plane. Stereological principles were followed (Whoolery *et al*. 2017; Lagace *et al*. 2010). Quantification was performed in the SGZ along the entire anterior-posterior hippocampal axis (–0.82 to –4.33mm from bregma). When counted, cells were categorized as DCX+ immature neurons (brown-stained soma located in SGZ, presence of neurite and dendrite outgrowth, some dendritic branching) or DCX+ progenitor cells (brown-stained soma located in the SGZ, no neurite outgrowth). For total cell number, the value collected for each mouse was multiplied by the section sampling fraction (here 1/18 since one hemisphere was counted). For cell numbers across bregma, the raw value collected at each bregma level is provided.

### 2.9 Computer Scripts

A custom Python (3.8.6) script developed by the Eisch Lab was used to extract touchscreen data, calculate outcome values, and compile the data into a database (Yun *et al*. 2023). These scripts and sample data files are available at https://github.com/EischLab/ts-data-analysis-app. Extracting the data into an output CSV file was managed with another custom script, and these outputs were verified manually. Following this verification, the data were analyzed using GraphPad Prism 10 according to the tests detailed in the Statistical Analysis section. Survival analyses were performed in R, and a full analysis notebook can be found at https://github.com/EischLab/19AFemalesSurvival.

### 2.10 Randomization, Blinding, Subject Number, and Data Removal

At the start of the experiment, cages of mice were weighed to ensure similarities. Afterwards, cages were randomly assigned to one of the four experimental groups: Veh/Sham (n=16), Veh/33-GCR (n=16), CDDO-EA/Sham (n=16), CDDO-EA/33-GCR (n=16). For neurogenesis studies, the subjects (n=11/group) were randomly chosen among animals that underwent behavior tests. Experimenters were not blinded to Diet or Radiation group at BNL but were blinded to these variables throughout the subsequent behavioral experiments. The subject number of each group in each experiment is provided in **Supplementary Table 1** (**Supp. Table 1**). The following rules applied for data removal/censorship. Mice were considered for removal from the entire study if they had husbandry issues (n=2, Veh/Sham mice, flooded cage on week 6) or veterinarian recommendation (n=3; n=1 CDDO-EA/33-GCR mouse, hindlimb wounds on week 5; n=1 CDDO-EA/Sham mouse, severe dermatitis and hunched posture on week 51; n=1 CDDO-EA/33-GCR mouse, underweight, abnormal gait, and hunched posture on week 55). Ultimately, data from the 3 mice who were considered for removal early in the study (n=2 Veh/Sham mice from week 6 and n=1 CDDO-EA/33-GCR mouse from week 5) were excluded from the entire study. Data from the remaining two mice were included in all graphs up until their removal from the study in weeks 51 and 55, respectively. For general touchscreen training and LDR studies, the subject numbers were: Veh/Sham (n=14), Veh/33-GCR (n=16), CDDO-EA/Sham (n=16), CDDO-EA/33-GCR (n=15). For EPM, the subject numbers were: Veh/Sham (n=14), Veh/33-GCR (n=16), CDDO-EA/Sham (n=16), CDDO-EA/33-GCR (n=14; one CDDO-EA/33-GCR mouse stepped off of the EPM). For Acq, the subject numbers were: Veh/Sham (n=12; two did not meet Acq criteria), Veh/33-GCR (n=13; three did not meet Acq criteria), CDDO-EA/Sham (n=14; two did not meet Acq criteria), CDDO-EA/33-GCR (n=14; two did not meet Acq criteria). For Ext, the subject numbers were: Veh/Sham (n=12; carried through from Acq), Veh/33-GCR (n=12; 13 carried through from Acq, and one additional did not meet Ext criteria), CDDO-EA/Sham (n=14; carried through from Acq), CDDO-EA/33-GCR (n=14; carried through from Acq). For measures collected in general activity chambers, there were 6 subjects whose data were censored: 5 due to equipment failure (4 Veh/33-GCR, 1 Veh/Sham) and 1 CDDO-EA/Sham mouse due to outlier status. For 3-CSI, in the social novelty phase, 1 Veh/Sham mouse was excluded due to complete lack of movement, and 2 Veh/33-GCR mice were excluded due to equipment failure. For 3-CSI, the data from one Veh/Sham mouse was not collected during habituation due to equipment failure, and therefore the mouse’s entire data (including sociability and social novelty) was removed from 3-CSI. During 3-CSI social novelty, the data from an additional Veh/Sham and two Veh/33-GCR mice were censored due to equipment failure. For OF, four Sham/CDDO-EA and four Veh/33-GCR mice were censored due to equipment failure on Days 1 and 2, respectively. Also, the data from one additional Sham/CDDO-EA mouse and one Veh/Sham subject were removed due to outlier testing (ROUT).

### 2.11 Statistical Analysis

Robust outlier testing (ROUT, Q=1%) was performed for all measures. For normality testing, within-group distributions were assessed by the D’Agostino-Pearson test and/or Shapiro-Wilk test and visualized via Quantile-Quantile plots. Most data were normally distributed and thus parametric statistics were used for main effects and interactions. If data were not normally distributed, non-parametric testing was done via Kruskal-Wallis tests for main effects of Treatment (combined variable of Diet/IRR) followed by Dunn’s Correction for post-hoc analysis. If samples were missing or data was censored, a mixed model ANOVA was used. Additional non-parametric testing was done for survival and touchscreen completion measures, which were evaluated by the Log-Rank (Mantel-Cox) test. Analyses were performed on the following variables (levels): Diet (Veh, CDDO-EA), IRR (Sham, 33-GCR), Bregma (12 distances from bregma). Test-specific statistical variables with comparisons included: for 3-CSI, Exploration time (Left vs Right Chamber), Sniff Zone (Stranger 1 vs Empty, Stranger 1 vs Stranger 2); for NOR, Object (Familiar vs Novel). LDR Test analysis used a combined variable of Treatment (Diet/IRR) and repeated measures (Block), with subsequent Block analysis performed using a two-way ANOVA of Diet x IRR. Other analyses that used a combined variable of Treatment (Diet/IRR) included stimulus-response Acq and Ext, bregma analysis for DCX+ cells and alopecia analysis. Full survival and weights statistical analysis: https://github.com/EischLab/NSRL19AFemales. For effect size determination after ANOVA, partial omega squared (ω^2^) or generalized eta squared (ges; η^2^) are reported with ≤0.01 small effect size, ≤0.06 medium effect size, and ≤0.13 large effect size (Olejnik and Algina 2003; Lakens 2013)(https://doomlab.shinyapps.io/mote/) or RStudio, and 95% confidence interval (CI) is reported up to two significant digits. For effect size determination after Kruskal-Wallis, partial eta squared is reported with ≤0.01 small effect size, ≤0.06 medium effect size, and ≤0.14 large effect size (Olejnik and Algina 2003; Lakens 2013). If the 95% CI full, non-abbreviated value includes zero, the effect size is considered similar to zero (**Supp. Table 1**), although post-hoc analyses will still be performed if the main effect or interaction of P value(s) is/are <0.05. For effect size determination of post-hoc analysis of multiple comparisons, Cohen’s d and Cliff’s delta are reported for ANOVA results and Kruskal-Wallis test, respectively (https://www.psychometrica.de/effect_size.html). In the Results section, values for F, P, partial ω^2^, partial η^2^, post-hoc P, Cohen’s d, and Cliff’s delta are reported only when ANOVA analysis (main effect and/or interaction) or Kruskal-Wallis results in P<0.05. However, all statistics for all data are provided in **Supp. Table 1**. Data in line graphs are expressed as mean+/-SEM, data in violin plots are expressed as median+/-interquartile range, and data in Kaplan-Meier curves are expressed as +/-95% CI for survival. Percent change of a given effect was calculated from means which are provided in **Supp. Table 1**. Critical α was set to 0.05 for all test statistics. Significant P values are bolded in **Supp. Table 1** and reported unrounded to the third decimal. For main effects and interactions, the following applies: * P<0.05, ** P<0.01, *** P<0.001, and **** P<0.0001. For post-hoc and multiple comparisons, a lower case, italicized letter indicates specific group differences: *a*, Veh/Sham vs Veh/33-GCR; *b,* Veh/Sham vs. CDDO-EA/Sham; *c,* Veh/Sham vs. CDDO-EA/33-GCR; *d,* Veh/33-GCR vs. CDDO-EA/Sham; *e,* Veh/33-GCR vs CDDO/33-GCR; *f*, CDDO-EA/Sham vs. CDDO-EA/33-GCR. The P value for post-hoc analyses is indicated by the number of apostrophes after the lower case italicized letter: *x* P<0.05, *x’* P<0.01, *x’’* P<0.001, and *x’’’* P<0.0001. Post-hoc values of 0.05<P<0.08 are only considered notable (and indicated on graph by #) when two additional conditions are met: if effect size is medium or larger and if post-hoc 95% CI does not include zero (**Supp. Table 1**). Otherwise, the 0.05<P<0.08 is listed in **Supp. Table 1** but not considered relevant for data interpretation. Statistics were computed using Prism 10 (Graphpad) and R studio. A complete report of distributions, test statistics for main effects, P values, intergroup comparisons, corrections, sample sizes and effect sizes per measure are provided in **Supp. Table 1**.

### 2.12 Graphs, Schematics and Photomicrographs

Graphs were generated in Prism 10 and RStudio. Brightfield photomicrographs were collected with an Olympus digital camera and cellSens Standard software. The calibrated scale bar was photographed at the same objective used for the photomicrographs. Original photomicrographs were cropped in Adobe Photoshop, and the crop was pasted into a new Photoshop window with a 300-dpi resolution and saved as a .tif. The .tif was then placed into the figure layout in Adobe Illustrator and scaled down to final size. Final figures and schematics were generated in Adobe Illustrator (v2023).

## 3. RESULTS

The entire experimental period from mouse arrival at BNL at 6-mon-of-age to tissue collection at 20.25-mon of age (14.25 mon post-IRR) was 8.25 mon (**Fig. 1A, Supp. Table 1**). At no time was the weight of Veh/33-GCR mice different from the weight of Veh/Sham mice (**Fig. 1B, 1C, Supp. Table 1**). Beginning two-month post-IRR, all mice received food during a restricted period of time in preparation for operant touchscreen training and testing. At one time point before and most time points after food restriction, CDDO-EA/Sham mice showed lower weight relative to Veh/Sham mice, and at two time points CDDO-EA/33-GCR mice weighed slightly less than CDDO-EA/Sham mice (**Fig. 1B, Supp. Table 1;** interactions: IRR x Time F(4.35, 243.64)=2.692, P=0.028 and Diet x Time F(4.35,243.64)=3.710, P<0.001; post-hoc: Veh/Sham vs CDDO-EA/Sham: *b* P<0.05, *b’* P<0.01 and CDDO-EA/Sham vs CDDO-EA/33-GCR: *f* P<0.05). For the weight collected at the end of the experiment (14.25 mon post-IRR), CDDO-EA/Sham mice weighed on average 12.5% less than Veh/Sham mice (**Fig. 1C, Supp. Table 1;** main effect: Diet F(1,55)=5.057, P=0.028, partial ω^2^=0.06 (Medium), 95% CI [0.00,0.23]; post-hoc: Veh/Sham vs CDDO-EA/Sham: *b#* P=0.0652, Cohen’s d=0.885 (Large), 95% CI [-1.43,-0.111]). These body weight data show three things: 1) 33-GCR on its own does not change body weight, 2) CDDO-EA given for 5 days at 6-mon-of-age slightly decreases body weight for much of the experiment between two-mon and 14.5-mon post-IRR, and 3) exposure to 33-GCR eliminates this CDDO-EA-induced lower weight.

The four groups did not differ in overall survival (**Supp. Fig. 1A, Supp. Table 1**). The groups did differ in coat score collected 10.5 mon (42 weeks) and 13.5 mon (54 weeks) post-IRR. At 42 weeks, CDDO/33-GCR mice had worse alopecia than Veh/Sham mice and at 54 weeks, Veh/33-GCR and CDDO-EA/Sham mice had worse alopecia than Veh/Sham mice (**Supp. Fig. 1B, Supp. Table 1**; Treatment H=40.55, P<0.0001, partial ω^2^=0.23 (Large); post-hoc 42 weeks: Veh/Sham vs CDDO-EA/33-GCR, *c* P=0.010, Cliff’s delta=0.8 (Large), 95% CI [0.467,0.933], 54 weeks: Veh/Sham vs Veh/33-GCR: *a’* P=0.007, Cliff’s delta=0.728 (Large), 95% CI [0.357,0.911], 54 weeks: Veh/Sham vs CDDO-EA/Sham: *b* P=0.0234, Cliff’s delta=0.714 (Large), 95% CI [0.324,0.895]). These data suggest the exposure to 33-GCR or CDDO-EA diet at 6 mon-old may contribute worse coat condition near the end of the experiment (13.5 months-old age).

Locomotion in the touchscreen chambers during Hab 1 was the same among all four groups **(Fig. 2A)**, indicating no pre-existing differences in movement in the novel touchscreen chambers. There were six general touchscreen phases, and CDDO-EA influenced the number of days it took to reach the criterion in two phases: MT and PI **(Fig. 2B)**. In MT, CDDO-EA/Sham mice completed MT on average 2.8 days faster than Veh/Sham mice (**Fig. 2B, Supp. Table 1;** main effect: Diet F(1,57)=13.15, P=0.0006, partial ω^2^=0.17 (Large), 95% CI [0.03,0.36]; post-hoc: Veh/Sham vs CDDO-EA/Sham: *b* P=0.015, Cohen’s d=0.851 (Large), 95% CI [-1.43,-0.119]). Also, CDDO/33-GCR mice completed MT on average 2.4 days faster than Veh/33-GCR mice (**Fig. 2B, Supp. Table 1**; post-hoc: Veh/33-GCR vs CDDO/33-GCR: *e* P=0.041, Cohen’s d=1.09 (Large), 95% CI [-1.72,-0.372]). In PI, there was a main effect of the CDDO-EA diet, but no post-hoc differences (**Fig. 2B, Supp. Table 1;** main effect: Diet F(1,57)=6.838, P=0.011, partial ω^2^=0.09 (Medium), 95% CI [0.00,0.26]). PI is distinct from other general touchscreen training phases in that accuracy — in addition to days to completion — can be measured. Notably, during PI CDDO-EA influenced measures relevant to accuracy. Specifically, CDDO-EA/33-GCR mice took on average 44.4% fewer trials to reach criteria relative to Veh/33-GCR (**Fig. 2C, Supp. Table 1**; main effect: Diet F(1,57)=10.79, P=0.001, partial ω^2^=0.14 (Large), 95% CI [0.02,0.32]; post-hoc: Veh/33-GCR vs CDDO-EA/33-GCR: *e* P=0.011, Cohen’s d=0.845 (Large), 95%CI [-1.27,-0.273]) and made on average 50.9% fewer blank touches relative to Veh/33-GCR (**Fig. 2D, Supp. Table 1**; main effect: Diet F(1,57)=10.98, P=0.001, partial ω^2^=0.14 (Large), 95% CI [0.02,0.33]; post-hoc: Veh/33-GCR vs CDDO-EA/33-GCR: *e’* P=0.008, Cohen’s d=0.885 (Large), 95% CI [-1.32,-0.244]). Also in PI, CDDO-EA/Sham mice made correct touches on average 22.7% faster (had a shorter latency) relative to Veh/Sham mice (**Fig. 2E, Supp. Table 1**; main effect: Diet F(1,57)=11.71, P=0.001, partial ω^2^=0.15 (Large), 95% CI [0.02,0.34]; post-hoc: Veh/Sham vs CDDO-EA/Sham: *b’* P=0.008, Cohen’s d=1.07 (Large), 95% CI [-1.88,-0.257]). Together these data suggest that by 2.5 mon post-CDDO-EA and post-IRR, CDDO-EA mice — regardless of whether in the Sham or 33-GCR group — learned the general touchscreen training better/faster than Veh mice.

**Figure 2.**
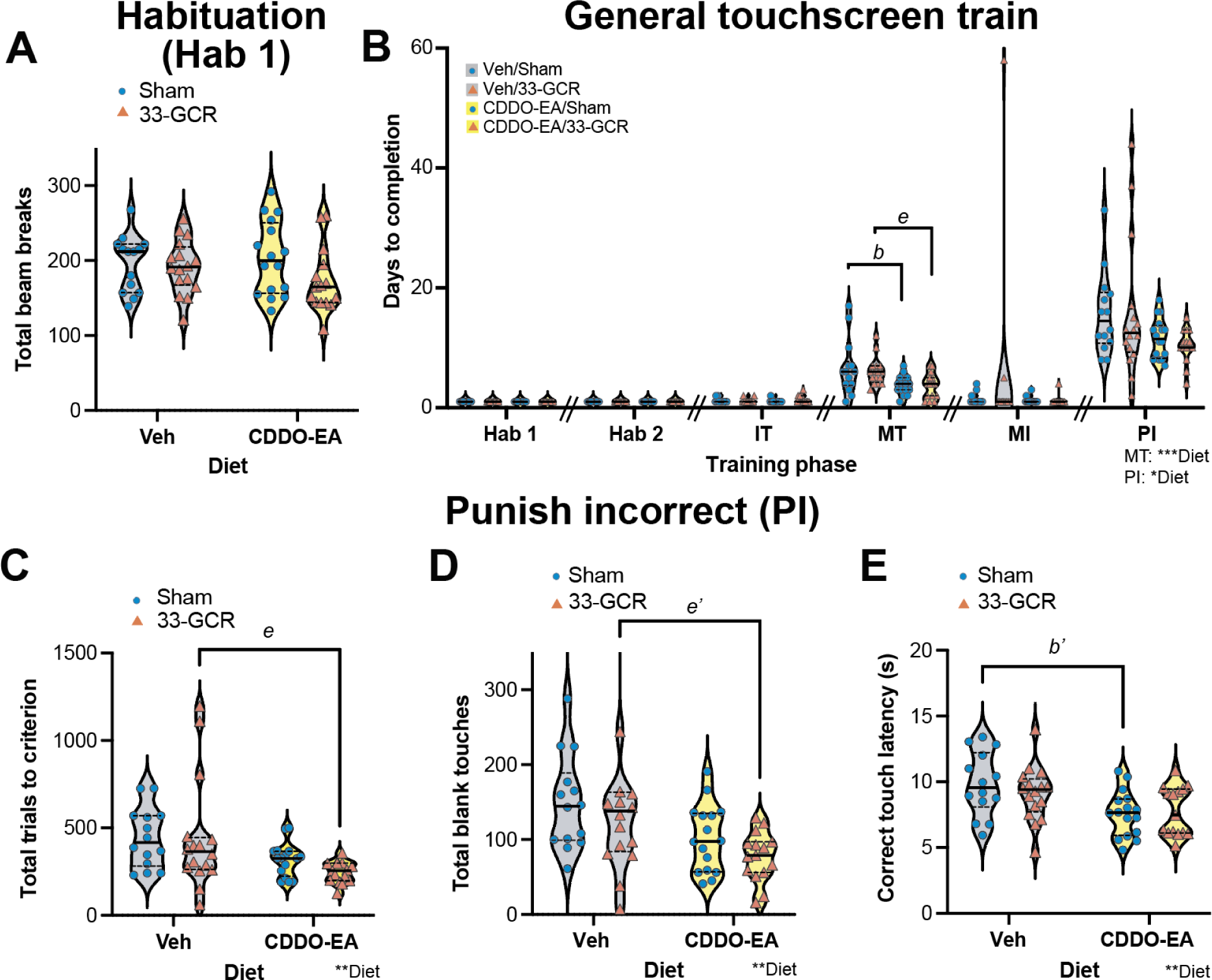
Two and one-half mon post-IRR, total beam breaks in a novel touchscreen operant chamber are similar among all groups, but some measures of general touchscreen performance were decreased by CDDO-EA, suggesting improved/faster learning. (**A**) <u>Beam breaks in novel environment.</u> The first time the mice were ever in the touchscreen chamber (the 10-min Habituation 1 [Hab1]), all four groups had a similar number of total beam breaks. **(B)** <u>Days to Completion for general touchscreen training phases.</u> In regard to Days to Completion for each phase of the general touchscreen training, there was a main effect of Diet in two phases: Must Touch (MT) and Punish Incorrect (PI). CDDO-EA/Sham mice completed MT 2.8 days earlier than Veh/Sham mice, CDDO/33-GCR mice completed MT 2.4 days faster than Veh/33-GCR mice. In PI, there was a main effect of the CDDO-EA diet, but no post-hoc differences. **(C-E)** <u>Measures relevant to accuracy from PI</u>. During PI, CDDO-EA/33-GCR mice took 44.4% fewer trials to reach criteria **(C)** and made 50.9% fewer blank touches **(D)** relative to Veh/33-GCR, suggesting faster learning. **(E)** Also in PI, CDDO-EA/Sham mice made correct touches 22.7% faster (shorter latency) relative to Veh/Sham mice. Statistical analysis in **A-E**: Two-way ANOVA (Diet x IRR). Main effect and interaction are denoted by *P<0.05, **P<0.01, ***P<0.001, post-hoc analysis significance denotations between groups: Veh/Sham vs CDDO-EA/Sham, *b* P<0.05 and *b’* P<0.01; Veh/33-GCR vs CDDO-EA/33-GCR, *e* P<0.05 and *e’* P<0.01. Initial Touch (IT) and Must Initiate (MI) in **B**: Kruskal-Wallis, P>0.05. 14-16 mice per group. IRR, irradiation. **A-E:** Veh, vehicle. Violin plots depict median (solid line) and 25% and 75% quartiles (dotted line). Complete subject numbers and detailed data analysis are provided in **Supp. Table 1.**

All mice completed LDR Train (earliest start: 3 mon post-IRR, **Fig. 3A**). All groups did so at a similar rate (**Fig. 3B, Supp. Table 1**) and in a similar number of days (**Fig. 3C, Supp. Table 1;** interaction: IRR x Diet F(1,56)=4.286, P=0.043, partial ω^2^=0.05 (Small), 95% CI [0.00,0.21]; post-hoc: all comparisons P>0.05). On the day each mouse completed LDR Train (Last Day), CDDO-EA/33-GCR mice took on average 6.8 (16.2%) more trials compared to Veh/33-GCR mice to reach the criteria and complete LDR Train (**Fig. 3D, Supp. Table 1;** main effect: Diet F(1,57)=11.90, P=0.001, partial ω^2^=0.15 (Large), 95% CI [0.02,0.34]; post-hoc: Veh/33-GCR vs CDDO-EA/33-GCR: *e’* P=0.003, Cohen’s d=1.14 (Large), 95% CI [0.478,1.88]). However, the remaining LDR Train Last Day metrics — session length (**Fig. 3E, Supp. Table 1**; main effect: Diet F(1,57)=7.293, P=0.009, partial ω^2^=0.09 (Medium), 95% CI [0.00,0.27]; post-hoc: all comparisons P>0.05) and percent correct, a measure of accuracy (**Fig. 3F, Supp. Table 1**) — were similar among the four groups. These data suggest that when LDR Train begins (3 mon post-CDDO-EA and post-IRR), all four groups of mice have roughly similar LDR Train performance metrics, including accuracy.

**Figure 3.**
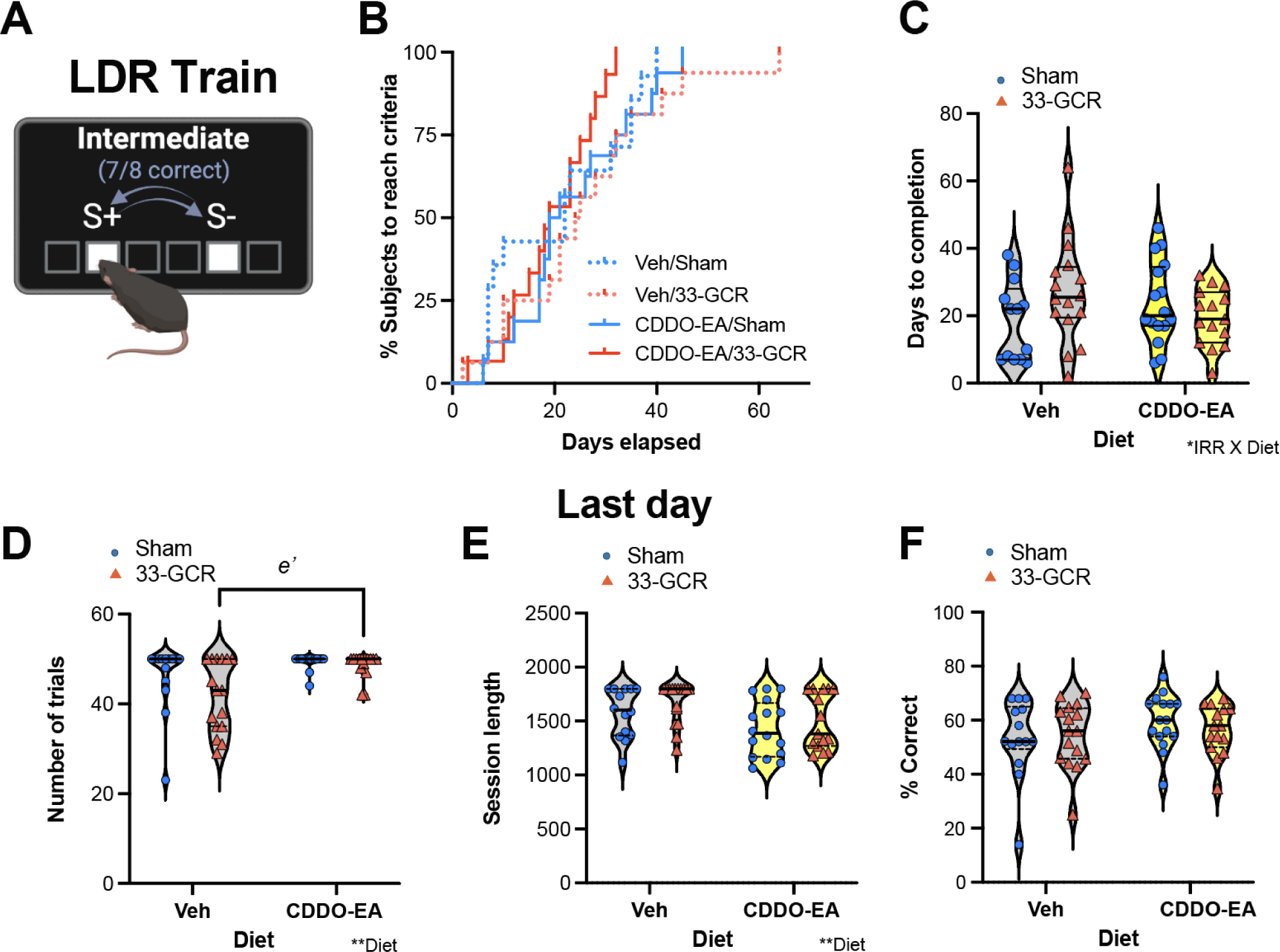
Three months post-IRR, performance in location discrimination reversal training (LDR Train) is generally similar among all groups. (**A**) <u>LDR Train touchscreen set-up.</u> Schematic of the lit squares and their “Intermediate” separation which were used for LDR Train. After a subject made 7 out of 8 correct touches, the windows indicated as S+ and S-switched. **(B)** <u>Percent of subjects reaching LDR Train criteria over time.</u> The percent of each group to reach criteria for LDR Train was similar among groups. **(C)** <u>Days to completion for LDR Train</u>. For the number of days it took each group to reach LDR Train completion, there was an IRR x Diet interaction but no post-hoc significance. **(D-F)** <u>LDR Train performance metrics on the day each mouse reached criteria (“Last Day”).</u> **(D)** On the Last Day of each mouse’s LDR Train, there was a main effect of Diet on the number of trials completed. Specifically, CDDO-EA/33-GCR mice took on average 6.8 (14%) more trials to complete Last Day LDR Train vs Veh/33-GCR mice. **(E-F)** However, on the Last Day of LDR Train, all groups had similar session length **(E)** and % correct **(F).** Violin plots depict median (solid line) and 25% and 75% quartiles (dotted line). Statistical analysis in **B**: Long-rank (Mantel-Cox) test, with a Kaplan-Meier completion graph and **C-F**: Two-way ANOVA (Diet x IRR). Main effect and/or interaction are denoted by *P<0.05, **P<0.01, post-hoc analysis significance denotations between groups: Veh/33-GCR vs CDDO-EA/33-GCR, *e’* P<0.01. Violin plots depict median (solid line) and 25% and 75% quartiles (dotted line). 14-16 mice per group. IRR, irradiation. Non-rewarding stimulus, S-, a touch to this lit square did not result in a reward. Rewarding stimulus, S+, a touch to this lit square resulted in a reward. Veh, vehicle. Complete subject numbers and detailed data analysis are provided in **Supp. Table 1.**

A major benefit of the LDR Test is that it can have varying “load”; placing the lit squares far apart (Large separation, **Fig. 4A-B**) is low load and considered discrimination while placing the lit squares close together (Small separation, **Fig. 4C-D**) is high load and considered pattern separation (Clelland *et al*. 2009; Oomen *et al*. 2013; Soler *et al*. 2021; Vivar and van Praag 2013; Zhuo *et al*. 2016). This assessment of discrimination (low load, Large separation) and pattern separation (high load, Small separation) can be measured by the percentage of trials which are correct <u>prior</u> to the first reversal. A second benefit of the five-month-long LDR Test is performance can be assessed over time in “blocks”; each of the five blocks consists of four days of LDR Test (two days Large separation + two days Small separation).

**Figure 4.**
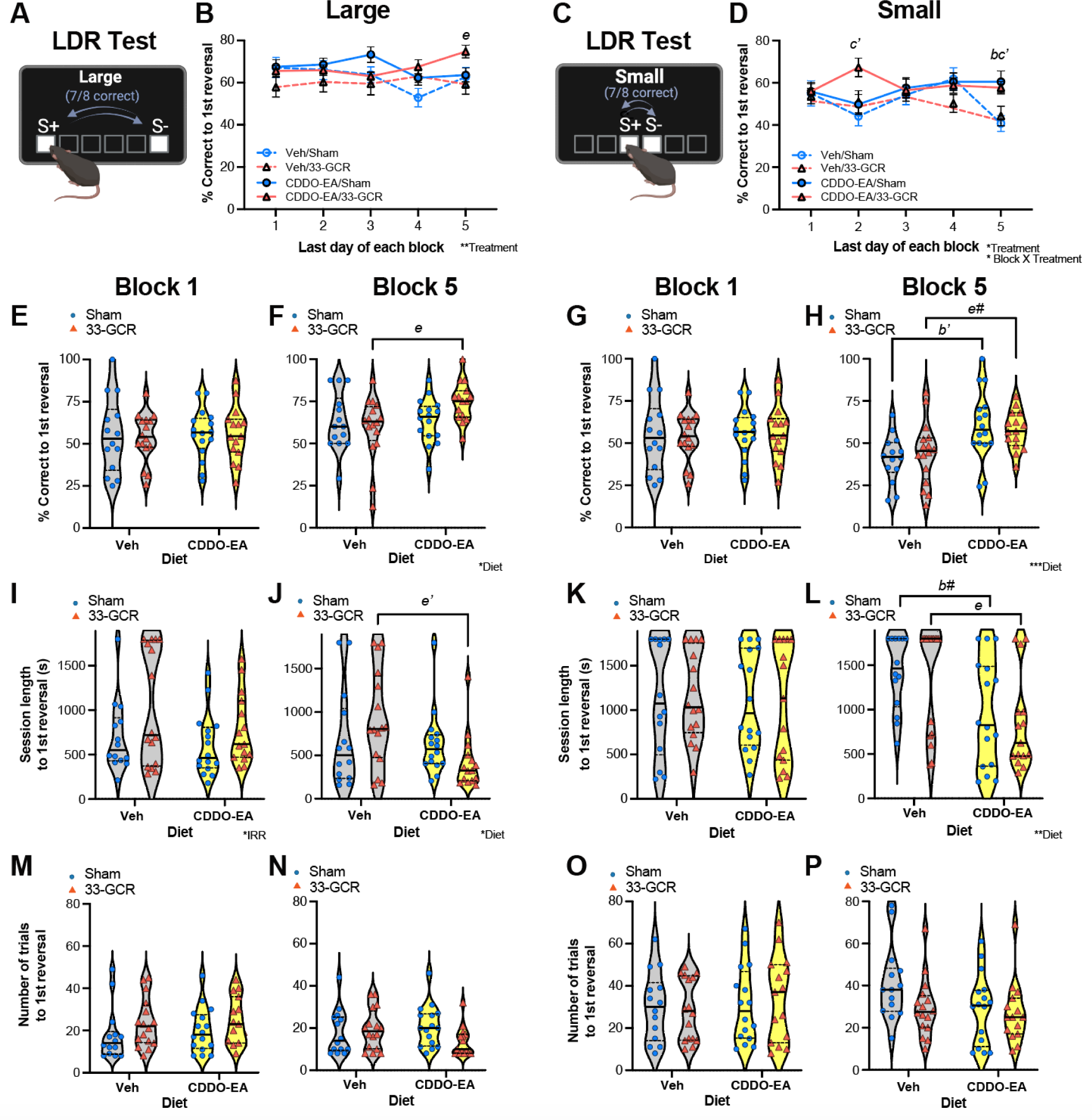
By the end of the 24-day LDR Test, CDDO-EA/33-GCR mice show better discrimination vs Veh/33-GCR mice, and CDDO-EA mice (regardless of IRR) show improved pattern separation vs Veh mice. **(A-B)** <u>Schematic of the lit squares in LDR Test with Large separation</u> **(A)** and <u>accuracy</u> to 1st reversal over 5 blocks in Large separation **(B)**. Large separation prior to the 1st reversal in LDR Test is considered a test of Discrimination due to the “load low” from the high resolution of the two, maximally separated squares. In Large separation in Block 5, CDDO-EA/33-GCR mice were 20.7% more accurate than Veh/33-GCR mice. **(C-D)** <u>Schematic of the lit squares in LDR Test with Small separation</u> **(C)** and <u>accuracy</u> to 1st reversal over 5 blocks in Small separation **(D).** Small separation prior to the 1st reversal in LDR Test is considered a test of Pattern Separation due to the “high load” from the low resolution of the two, minimally separated squares. In Small separation Block 5, CDDO-EA/Sham mice performed 32.7% better relative to Veh/Sham mice. In Block 2 and 5, CDDO-EA/33-GCR mice were 34.3% (Block 2) and 29.4% (Block 5) more accurate than Veh/Sham mice. **(E-P)** <u>Metrics of Discrimination (Large separation, **E, F, I, J, M, N**) and Pattern Separation (Small separation, **G, H, K, L, O, P**) in LDR test</u> were analyzed in Block 1 **(E, I, M, G, K, O)** and separately in Block 5 **(F, J, N, H, L, P)** based on accuracy **(E-H)**, session length **(I-L)**, and trial number to the 1st reversal **(M-P)**. In Large separation, all groups had similar accuracy **(E)** and speed **(I)** in Block 1, but in Block 5 CDDO-EA/33-GCR mice were 20.7% more accurate **(F)** and reached the 1st reversal 54% faster **(J)** vs Veh/33-GCR mice. In Small separation, all groups had similar accuracy **(G)** and session length to 1st reversal **(K)** in Block 1. In Block 5 of Small separation, CDDO-EA/Sham mice were 32.7% more accurate **(H)** and reached 1st reversal 32.3% faster **(L)** vs Veh/Sham mice, and CDDO-EA/33-GCR mice were 23.3% more accurate **(H)** and reached 1st reversal 38.1% faster **(L)** vs Veh/33-GCR mice. Statistical analysis in **B, D**: Two-way RM measures ANOVA (Block X Treatment), main effect and an interaction followed by Tukey’s post-hoc and **E-P**: Two-way ANOVA (Diet x IRR), main effect and interaction followed by Bonferroni’s post-hoc test. Main effects and/or interaction are denoted by *P<0.05, **P<0.01, ***P<0.001. Post-hoc analysis significance denotations between groups: Veh/Sham vs. CDDO-EA/Sham, *b#* 0.05<P<0.08, *b* P<0.05, *b’* P<0.01; Veh/Sham vs. CDDO-EA/33-GCR, *c’* P<0.01; Veh/33-GCR vs. CDDO-EA/33-GCR, *e#* 0.05<P<0.08, *e* P<0.05, *e’* P<0.01. 14-16 mice per group. Non-rewarding stimulus, S-, a touch to this lit square did not result in a reward. Rewarding stimulus, S+, a touch to this lit square resulted in a reward. IRR, irradiation. Veh, vehicle. Violin plots depict median (solid line) and 25% and 75% quartiles (dotted line). Complete subject numbers and detailed data analysis are provided in **Supp. Table 1.**

On the Last Day of each LDR Test block, mouse accuracy was gauged by percent correct up to first reversal (**Fig. 4B, D**). In Large separation of LDR Test in Block 5, CDDO-EA/33-GCR mice were 26.1% more accurate than Veh/33-GCR mice (**Fig. 4B, Supp. Table 1**; Treatment F(3,57)=3.079, P=0.034, partial ω^2^=0.17 (Large), 95% CI [0.08,0.25]; post-hoc: Veh/33-GCR vs CDDO-EA/33-GCR in Block 5: *e* P=0.048, Cohen’s d=0.979 (Large), 95% CI [0.285,1.51]). In Small separation of LDR Test, CDDO-EA/Sham mice performed 48.5% better relative to Veh/Sham mice in Block 5 (**Fig. 4D, Supp. Table 1**; interaction: Block x Treatment F(12,228)=1.837, P=0.043, partial ω^2^=0.11 (Medium), 95% CI [0.01, 0.15]; post-hoc: Veh/Sham vs CDDO-EA/Sham in Block 5: *b* P=0.021, Cohen’s d=1.11 (Large), 95% CI [0.326,1.83]). In the same Small separation LDR Test, CDDO-EA/33-GCR mice were more accurate than Veh/Sham mice in Block 2 (52.3%) and 5 (41.5%; **Fig. 4D, Supp. Table 1**; post-hoc: Veh/Sham vs CDDO-EA/33-GCR in Block 2: *c’* P=0.005, Cohen’s d=1.36 (Large), 95% CI [0.59,2.1]; Veh/Sham vs CDDO-EA/33-GCR in Block 5: *c’* P=0.008, Cohen’s d=1.32 (Large), 95% CI [0.474,2.06]). Separate examination of accuracy among groups on Block 1 revealed this improved accuracy was not evident in Block 1 of either Large or Small separation (**Fig. 4E, G, Supp. Table 1**). However, on Block 5, in Large separation CDDO-EA/33-GCR mice were on average 26.1% more accurate than Veh/33-GCR mice, although the effect size was small (**Fig. 4F, Supp. Table 1**; main effect; Diet F(1,57)=4.283, P=0.043, partial ω^2^=0.05 (Small), 95% CI [0.00, 0.20]; post-hoc: Veh/33-GCR vs CDDO-EA/33-GCR: *e* P=0.043, Cohen’s d=0.979 (Large), 95% CI [0.285,1.51]). On Block 5 in Small separation, CDDO-EA/Sham mice were on average 48.5% more accurate than Veh/Sham mice (**Fig. 4H, Supp. Table 1**; Diet F(1,57)=15.08, P=0.0003, partial ω^2^=0.19 (Large), 95% CI [0.04, 0.38]; post-hoc: Veh/Sham vs CDDO-EA/Sham: *b’* P=0.004, Cohen’s d=1.11 (Large), 95% CI [0.326,1.83]) and CDDO-EA/33-GCR were on average 30.3% more accurate than Veh/33-GCR mice (**Fig. 4H, Supp. Table 1**; post-hoc: Veh/33-GCR vs CDDO-EA/33-GCR: *e#* P=0.057, 0.866 (Large), Cohen’s d=0.866 (Large), 95%C CI [0.0538,1.64]). Examination of a complementary variable — session length to first reversal — showed all groups performed similarly in Block 1 for both Large separation (**Fig. 4I, Supp Table 1**; IRR F(1,57)=4.589, P=0.036, partial ω^2^=0.06 (Medium), 95% CI [0.00, 0.021], suggesting effect size similar to zero) and Small separation (**Fig. 4K, Supp. Table 1**). On the Last Day of Block 5 in Large separation, CDDO-EA/33-GCR mice reached the first reversal on average 54% faster than Veh/33-GCR mice (**Fig. 4J, Supp. Table 1**; main effect: Diet F(1,57)=5.201, P=0.026, partial ω^2^=0.06 (Medium), 95% CI [0.00, 0.23]; post-hoc: Veh/33-GCR vs CDDO-EA/33-GCR: *e’* P=0.008, Cohen’s d=1.08 (Large), 95% CI [-1.81,-0.242]). However, on the Last Day of Block 5 in Small separation, CDDO-EA/Sham mice reached their first reversal on average 32.3% faster than Veh/Sham mice (**Fig. 4L, Supp. Table 1**; main effect: Diet F(1,57)=12.00, P=0.001, partial ω^2^=0.15 (Large), 95% CI [0.02,0.34]; post-hoc: Veh/Sham vs CDDO-EA/Sham: *b#* P=0.052, Cohen’s d=0.873 (Large), 95% CI [-1.64,-0.105]) and CDDO-EA/33-GCR mice reached their first reversal on average 38.1% faster than Veh-33-GCR mice (**Fig. L, Supp. Table 1**; post-hoc: Veh/33-GCR vs CDDO-EA/33-GCR: *e* P=0.022, Cohen’s d=0.906 (Large), 95% CI [-1.86,-0.0809]. Notably, the number of trials to first reversal did not differ among groups on the Last Day of Block 1 or on the Last Day of Block 5 in either Large separation (**Fig. 4M, 4N, Supp. Table 1**) or Small separation (**Fig. 4O, 4P, Supp. Table 1**). These data show that while mice perform LDR Test similarly in Block 1, by Block 5 CDDO-EA/33-GCR mice show better discrimination vs Veh/33-GCR mice and CDDO-EA mice (regardless of IRR) show better pattern separation vs Veh mice.

A third benefit of the LDR Test is that the number of “reversals” can be measured, and that reversal number can reflect cognitive flexibility (Swan *et al*. 2014). The number of reversals are typically analyzed in Blocks at the end of a LDR Test (Swan *et al*. 2014; Soler *et al*. 2021; Yun *et al*. 2023), as mice should be the most familiar with the LDR Test at this point. Thus, our main focus was the number of reversals on the Last Day of Block 5. In Block 5, CDDO-EA/33-GCR mice made on average 80.8% and 180% more reversals than Veh/33-GCR mice during Large separation (**Fig. 5A**) and Small separation (**Fig. 5B**), respectively (Large, **Fig. 5A, Supp. Table 1**; main effect: Diet F(1,57)=7.725, P=0.007, partial ω^2^=0.10 (Medium), 95% CI [0.00,0.28]; post-hoc: Veh/33-GCR vs CDDO-EA/33-GCR: *e* P=0.021, Cohen’s d=1.15 (Large), 95% CI [0.28,2.11]; Small, **Fig. 5B, Supp. Table 1**; main effect: Diet F(1,57)=8.877, P=0.004, partial ω^2^=0.11 (Medium), 95% CI [0.01,0.30]; post-hoc: Veh/33-GCR vs CDDO-EA/33-GCR: *e* P=0.018, Cohen’s d=0.998 (Large), 95% CI [0.177, 1.79]). These data suggest that CDDO-EA improves cognitive flexibility in IRR mice.

**Figure 5.**
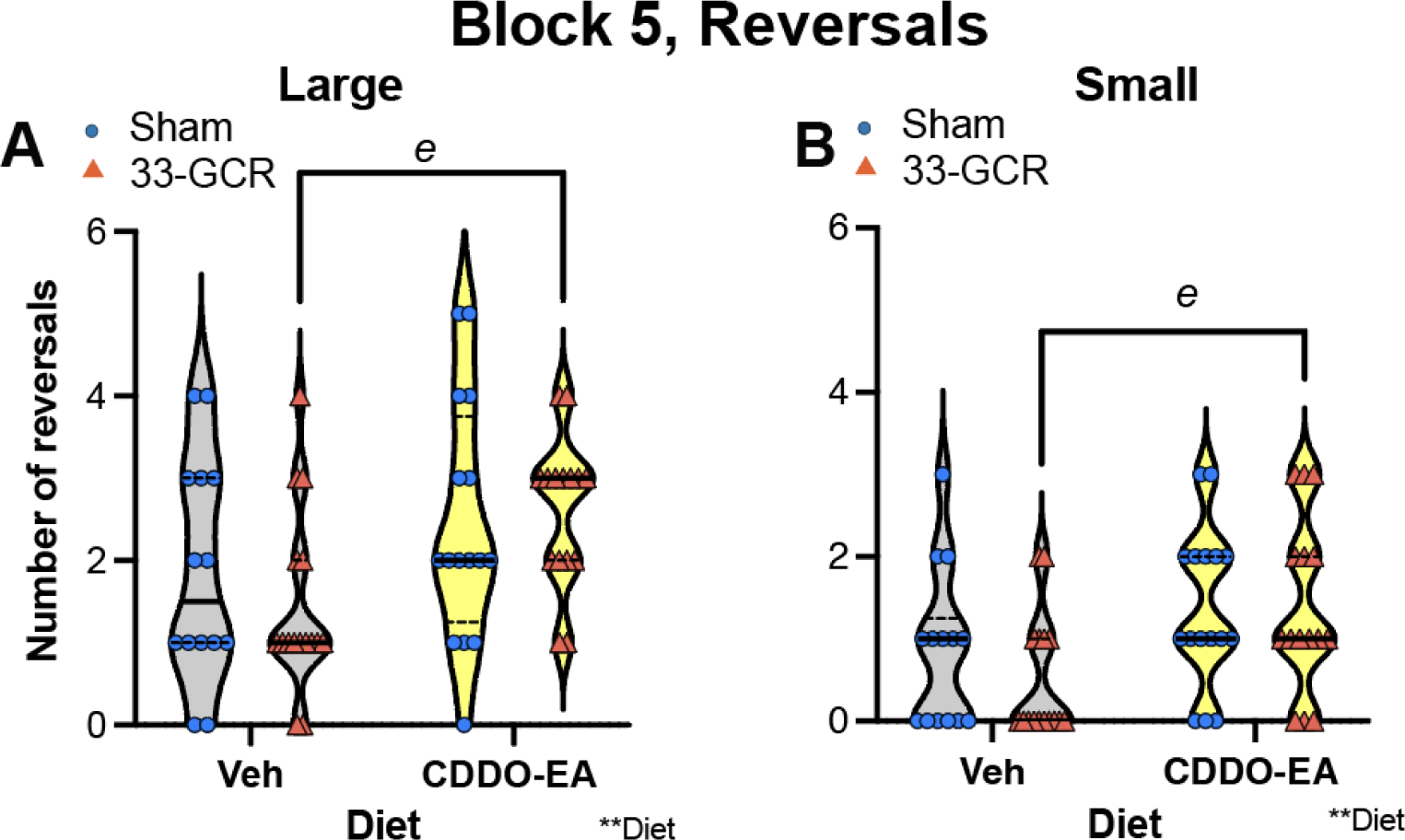
By the end of LDR Test (and 3 to 8.5 mon post-IRR), cognitive flexibility is improved by CDDO-EA during IRR exposure. <u>Number of reversals on the Last Day of the last block (Block 5) of LDR Test under conditions of Large **(A)** and Small **(B)** separation.</u> **(A)** <u>Large separation.</u> In Large separation Block 5, CDDO-EA/33-GCR mice made 44.7% more reversals than Veh/33-GCR mice during Large separation. **(B)** <u>Small separation.</u> In Small separation Block 5, CDDO-EA/33-GCR mice made 64.3% more reversals than Veh/33-GCR mice. Two-way ANOVA (IRR X Diet), main effect and/or interaction followed by Bonferroni’s post-hoc test. Main effects are denoted by **P<0.01. Post-hoc analysis significance denotations between groups: Veh/33-GCR vs CDDO-EA/33-GCR, *e* P<0.05. 14-16 mice per group. IRR, irradiation. Veh, vehicle. Violin plots depict median (solid line) and 25% and 75% quartiles (dotted line). Complete subject numbers and detailed data analysis are provided in **Supp. Table 1.**

On the EPM, all groups spent similar time in the open arms **(Fig. 6A, Supp. Table 1)** and made a similar number of visits to the open arms **(Fig. 6B, Supp. Table 1)**. All groups had similar EPM exploration indices (time spent in the open vs closed arms, **Fig. 6C, Supp. Table 1**) and moved a similar distance during the 5-min EPM test **(Fig. 6D, Supp. Table 1)**. These data suggest that at this time point (8.8 mon post-IRR), neither CDDO-EA nor 33-GCR influence this measure of anxiety-like behavior.

**Figure 6.**
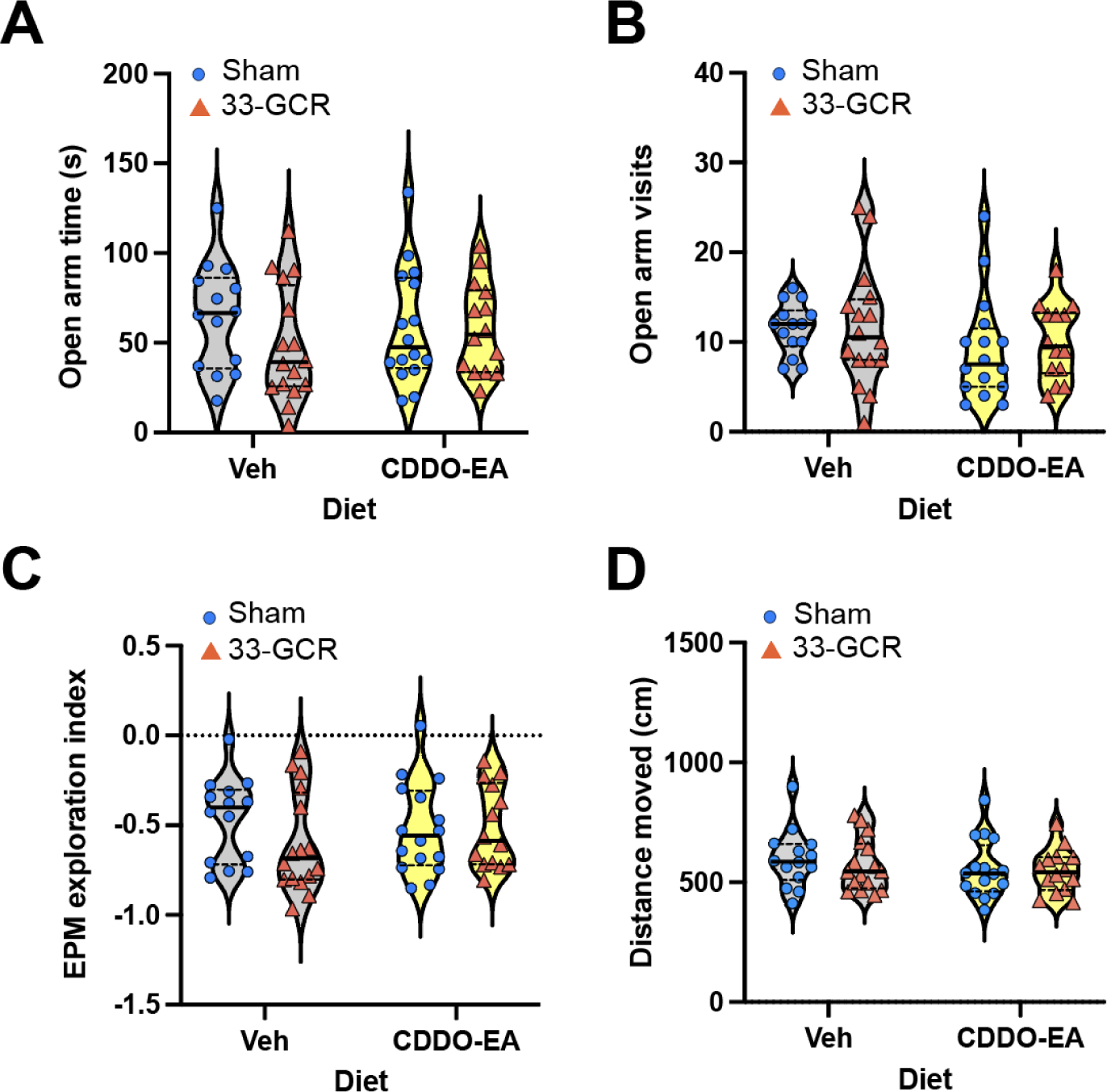
Eight months post-IRR, anxiety-like behavior on the elevated plus maze (EPM) is similar among all groups. (**A-D**) EPM metrics of **(A)** time in open arms, **(B)** visits to open arms, **(C)** exploration index (time spent in open vs closed arms), and **(D)** distance moved. There was no difference among the four groups in time spent in the open arms **(A)**, visits to the open arms **(B),** exploration indices **(C),** and distance moved during the 5min EPM test **(D)**. Two-way ANOVA (IRR X Diet), main effects and interaction are all P>0.05 in **A-D**. 14-16 mice per group. IRR, irradiation. Veh, vehicle. Violin plots depict median (solid line) and 25% and 75% quartiles (dotted line). Complete data analysis details are provided in **Supp. Table 1.**

Mice were reintroduced to the now-familiar touchscreen chambers for a second 10-min habituation (Hab 1) session as part of the training for the subsequent Acq and Ext testing. Veh/33-GCR mice moved on average 30.4% less in these familiar chambers 9.25 mon post-IRR vs Veh/Sham mice (**Fig. 7A, Supp. Table 1**; main effect: IRR F(1,49)=10.01, P=0.002, partial ω^2^=0.11 (Medium), 95% CI [0.01,0.30], post-hoc: Veh/Sham vs Veh/33-GCR: *a#* P=0.063, Cohen’s d=1.21 (Large), 95% CI [-2.22,-0.216]). These data suggest that mice that were exposed to Veh/33-GCR moved less in these familiar chambers 9.25 mon post-IRR compared to Veh/Sham mice, and that this difference is not seen in CDDO-EA/33-GCR vs CDDO-EA/Sham mice.

**Figure 7.**
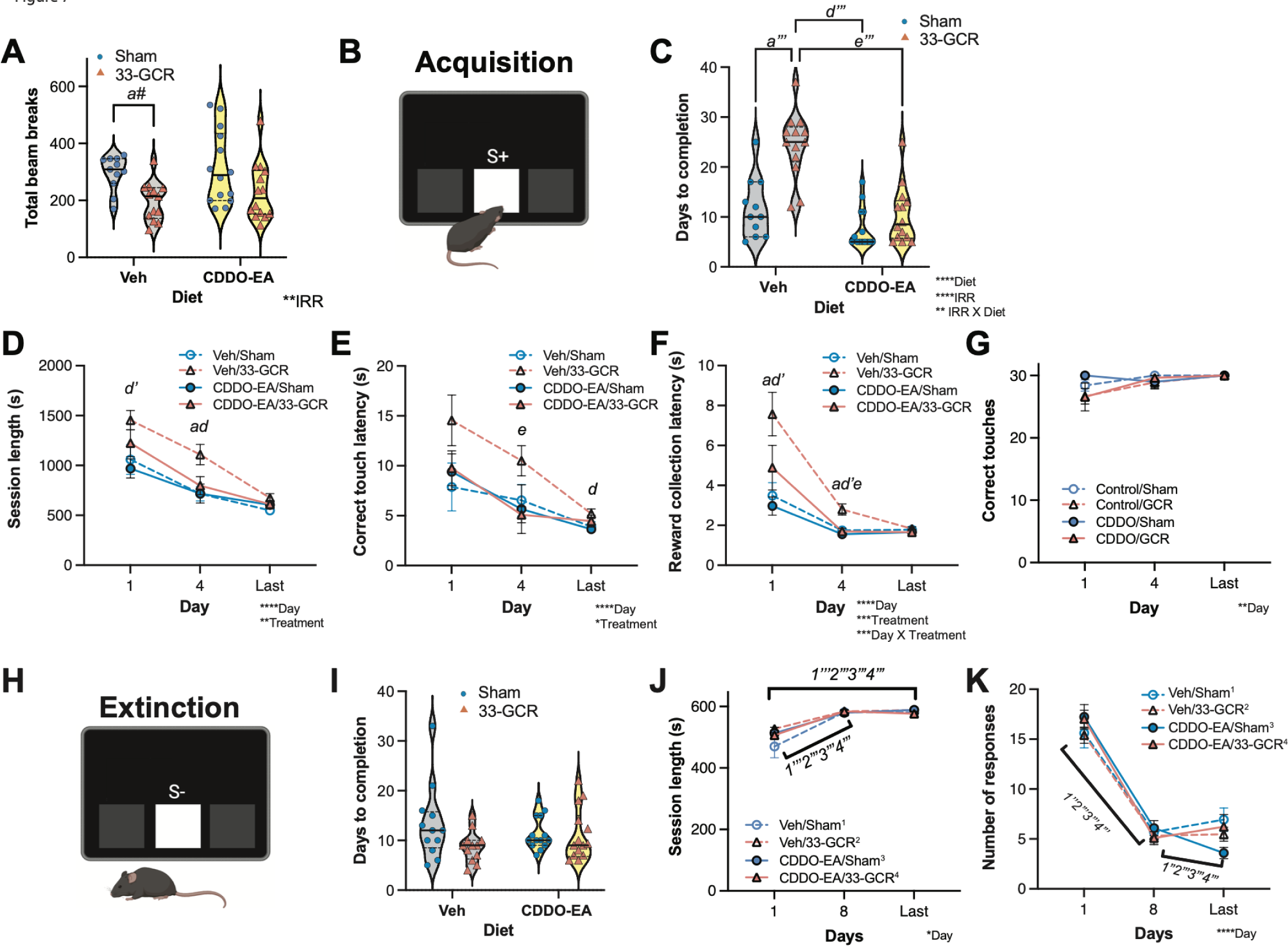
9.25 months post-IRR, 33-GCR decreases locomotion in a familiar environment and acquisition of a simple stimulus-response learning task, and CDDO-EA appears to counter the 33-GCR effects. No effect is seen on subsequent extinction learning. (**A**) <u>Beam breaks in familiar operant environment.</u> Mice were reintroduced to the touchscreen chamber (10-min Habituation 1 [Hab1]) 3 weeks post-LDR to measure locomotion activity via beam breaks. Mice exposed to Veh/33-GCR moved 30.4% less in these familiar chambers 9.25 mon post-IRR compared to Veh/Sham mice; this difference is not seen in CDDO-EA/33-GCR vs CDDO-EA/Sham mice. **(B)** <u>Schematic of simple stimulus-response learning, “acquisition (Acq)”</u>. The middle-lit square (S+) was the conditioned stimulus. Touches to S+ led to reward delivery, while touches to blank areas on either side of S+ did not. **(C)** <u>Days to complete (acquire) simple stimulus-response learning.</u> Mice are run for 30 min, but only the first 15-min are considered for these data. Veh/33-GCR mice took 47.4% more days vs Veh/Sham, CDDO-EA/Sham mice took 69% fewer days vs Veh/33-GCR, and CDDO-EA/33-GCR mice took 57.8% fewer days vs Veh/33-GCR mice. **(D-F)** <u>Acq metrics</u> were analyzed in the 1st, 4th, and Last Day based on session length **(D),** correct touch latency **(E),** and reward collection latency **(F)** Note that **D-G** are from the first 30-min of each session. **(D)** <u>Acq session length:</u> On Day 1, Veh/33-GCR mice had session length that was 33.3% longer than CDDO-EA/Sham mice. On Day 4, Veh/33-GCR mice had session length that was on average 35.5% longer than Veh/Sham mice, and Veh/33-GCR mice had on average a session length that was 35.3% longer than CDDO-EA/Sham mice. **(E)** <u>Acq correct touch latency:</u> On Day 4 Veh/33-GCR mice took 47% longer to make a correct touch compared to CDDO-EA/33-GCR mice. On the Last Day Veh/33-GCR mice took 30% longer to make a correct touch compared to CDDO-EA/Sham mice. **(F)** <u>Acq reward collection latency:</u> On Day 1, Veh/33-GCR mice took 54% longer to collect the reward compared to Veh/Sham mice and 60.8% longer to collect the reward compared to CDDO-EA/Sham mice. On Day 4, Veh/33-GCR mice took 37.5% longer to collect the reward compared to Veh/Sham mice, 44.3% longer to collect the reward compared to CDDO-EA/Sham mice, and 40.2% longer to collect the reward compared to CDDO-EA/33-GCR mice. **(G)** <u>Acq correct touches:</u> On all days, all mice had a similar number of correct touches. **(H)** Schematic of a middle-lit square (S-) in extinction learning. **(I-K)** <u>Measures of Ext</u> after a simple-response learning based on days to completion **(I)**, session length **(J)**, and response number (#; **K**). **(I)** <u>Ext days to completion</u>: All four groups took a similar number of days to complete extinction learning. **(J)** <u>Ext session length:</u> Within each group, Day 1 was a shorter session vs the Day 8 session and the Last Day session. **(K)** <u>Ext number of responses:</u> Within each group, the fewest number of responses were on Day 8 and the Last Day vs Day 1. Statistical analysis: **A, C, I:** Two-way ANOVA (Diet x IRR); **D, J, K**: Two-way RM (Day x Treatment); **E, F, G:** Mixed Effects 2-way RM ANOVA (Day x Treatment). Main effects and interaction are denoted by *P<0.05, **P<0.01, ***P<0.001, or ****P<0.0001. Bonferroni **(A)** or Tukey **(C-G)** post-hoc analysis significance denotations between groups: Veh/Sham vs. Veh/33-GCR, *a#* 0.08< P<0.05, *a* P<0.05, *a’’’* P<0.001; Veh/33-GCR vs. CDDO-EA/Sham, *d* P<0.05, *d’* P<0.01, *d’’’* P<0.001; Veh/33-GCR vs. CDDO-EA/33-GCR, *e* P<0.05, *e’’’* P<0.001; CDDO-EA/Sham vs. CDDO-EA/33-GCR, *f#* 0.08< P<0.05. Tukey **(J, K)** post-hoc analysis significance denotations between days within a given group: *1* = within Veh/Sham, *2* = within Veh/33-GCR; *3* = within CDDO-EA/Sham; *4* = within CDDO-EA/33-GCR with *’’* = P<0.001 and *’’*’ = P<0.0001. IRR, irradiation. Veh, vehicle. Violin plots (**A, C, I**) depict median (solid line) and 25% and 75% quartiles (dotted line). Complete subject numbers and detailed data analysis are provided in **Supp. Table 1.**

After Hab 1, mice began the daily 30-min Acq sessions (**Fig. 7B**). All groups of mice learned to touch the conditioned stimulus within the 30-min sessions (data not shown). However, Acq session data are typically analyzed in only the first 15-min of the session (Mar *et al*. 2013; Soler *et al*. 2021; Cotter *et al*. 2023; Bussey *et al*. 2012), as the faster learning sets the stage for learning Ext in subsequent sessions. Analysis of the first 15-min of the Acq sessions revealed a visible difference among groups with CDDO-EA/Sham mice learning fastest (mean ∼7.5 days) and Veh/33-GCR mice taking the longest time to learn (mean ∼24.4 days, **Fig. 7C**). Further analysis revealed Veh/33-GCR mice took on average 90% more days to complete Acq vs Veh/Sham (**Fig. 7C, Supp. Table 1**; interaction: IRR x Diet F(1,49)=7.457, P=0.0088, partial ω^2^=0.11 (Medium), 95% CI [0.00,0.30], post-hoc: Veh/Sham vs Veh/33-GCR: *a’’’* P<0.0001, Cohen’s d=1.67 (Large), 95% CI [0.632, 2.8]). In contrast, two CDDO-EA groups of mice learned Acq faster than Veh/33-GCR mice: CDDO-EA/Sham mice took on average 69% fewer days vs Veh/33-GCR mice (**Fig. 7C, Supp. Table 1**; post-hoc: Veh/33-GCR vs CDDO-EA/Sham: *d’’’* P<0.0001, Cohen’s d=3.09 (Large), 95% CI [-4.73,-1.77]) and CDDO-EA/33-GCR mice took on average 57.8% fewer days vs Veh/33-GCR mice (**Fig. 7C, Supp. Table 1**; post-hoc: Veh/33-GCR vs. CDDO-EA/33-GCR: *e’’’* P<0.0001, Cohen’s d=2.28 (Large), 95% CI [-3.63,-1.12]). Thus, when only the first 15-min were analyzed, as is standard in the field, Veh/33-GCR mice took more days to acquire this simple stimulus-response learning.

To gain greater insight into Acq learning, other metrics of performance from the full 30-min sessions were analyzed on Day 1, Day 4, and Last Day; Day 4 was selected since the Acq criteria requires meeting trial number of four consecutive days. Analysis of the 1st, 4th, and Last Day of each mouse’s Acq performance showed three effects on session length (**Fig. 7D**). On Day 1, the session length of Veh/33-GCR mice was on average 33.3% longer than CDDO-EA/Sham mice (**Fig. 7D, Supp. Table 1**; main effect of Treatment: F(3,49)=4.892, P=0.004, partial ω^2^=0.12 (Medium), 95% CI [0.01,0.24]; main effect of Day: F(1.819,89.15)=56.32, P<0.0001, partial ω^2^=0.34 (Large), 95% CI [0.19,0.48]; post-hoc: Veh/33-GCR vs CDDO-EA/Sham on Day 1: *d’* P=0.007, Cohen’s d=1.37 (Large), 95% CI [-2.32,-0.45]). On Day 4, Veh/33-GCR mice had session length that was on average 55% longer than Veh/Sham mice (**Fig. 7D, Supp. Table 1**; post-hoc: Veh/Sham vs Veh/33-GCR on Day 4: *a* P=0.022, Cohen’s d=1.25 (Large), 95% CI [0.456,1.98]). Also on Day 4, Veh/33-GCR mice had on average a session length that was 54.5% longer than CDDO-EA/Sham mice (**Fig. 7D, Supp. Table 1**; Veh/33-GCR vs CDDO-EA/Sham on Day 4: *d* P=0.046, Cohen’s d=1.08 (Large), 95% CI [-1.97,0.0495]). In regard to Acq correct touch latency, on Day 4 Veh/33-GCR mice took on average 88.6% longer to make a correct touch compared to CDDO-EA/33-GCR mice (**Fig. 7E, Supp. Table 1**; main effect of Day: F(1.868,90.62)=19.64, P<0.0001; main effect of Treatment: F(3,49)=2.844, P=0.047; Veh/33-GCR vs CDDO-EA/33-GCR Day 4: *e* P=0.048, Cohen’s d=1.11 (Large), 95% [-1.91,-0.335]), and on the Last Day Veh/33-GCR mice took on average 42.8% longer to make a correct touch compared to CDDO-EA/Sham mice (**Fig. 7E, Supp. Table 1**; post hoc Veh/33-GCR vs CDDO-EA/Sham Last Day: *d* P=0.047, Cohen’s d=1.11 (Large), 95% CI [-1.88,-0.299]. In regard to latency to collect the reward during Acq, On Day 1, Veh/33-GCR mice took on average 117% longer to collect the reward compared to Veh/Sham mice (**Fig. 7F, Supp. Table 1**; interaction: Day x Treatment: F(6,97)=4.430, P=0.0005; post hoc: Veh/Sham vs Veh/33-GCR on Day 1: *a* P=0.020, Cohen’s d=1.27 (Large), 95% CI [0.284, 2.02]). Also on Day 1, Veh/33-GCR mice took on average 155% longer to collect the reward compared to CDDO-EA/Sham mice (**Fig. 7F, Supp. Table 1**; post hoc: Veh/33-GCR vs CDDO-EA/Sham Day 1: *d’* P=0.006, Cohen’s d=1.55 (Large), 95% CI [-2.24,-0.743]). On Day 4, Veh/33-GCR mice took on average 59.9% longer to collect the reward compared to Veh/Sham mice (**Fig. 7F, Supp. Table 1**; post hoc: Veh/Sham vs Veh/33-GCR Day 4: *a* P=0.020, Cohen’s d=1.27 (Large), 95% CI [0.493,1.9]), Veh/33-GCR mice took on average 79.7% longer to collect the reward compared to CDDO-EA/Sham mice (**Fig. 7F, Supp. Table 1**; post hoc: Veh/33-GCR vs CDDO-EA/Sham Day 4: *d’* P=0.004, Cohen’s d=1.69 (Large), 95% CI [-2.34,-0.981]), and Veh/33-GCR mice took on average 67.3% longer to collect the reward compared to CDDO-EA/33-GCR mice (**Fig. 7F, Supp. Table 1**; post hoc: Veh/33-GCR vs CDDO-EA/33-GCR Day 4: *e* P=0.010, Cohen’s d=1.41 (Large), 95% CI [-2.1,-0.577]). Notably, mice of all groups showed similar correct touches on Days 1, 4 and the Last Day (**Fig. 7G, Supp. Table 1**). These data show three things: 1) 33-GCR exposure slows acquisition learning of conditioned stimulus learning 9-mon post-IRR; 2) Concomitant administration of CDDO-EA at the time of IRR normalizes key aspects of this learning; 3) The negative impact of 33-GCR is on speed of completing the task, and not on accuracy.

Following Acq, mice are presented with similar opportunities to respond to a visual stimulus, but no reward and no cues that are associated with the reward. The time it takes and the extent to which the mice suppress their response allows insight into their ability to extinguish prior learning. Ext performance over time was analyzed on Day 1, Day 8, and Last Day; Day 8 was selected since the average group Ext completion was ∼8 days. For Ext of the conditioned stimulus learning (**Fig. 7H**), all groups of mice learned to extinguish their simple stimulus-response association and did so over a similar number of days (**Fig. 7I, Supp. Table 1**). Analysis of the 1st, 8th, and Last Day of each mouse’s Ext performance showed within each group of mice, the length of the sessions on Day 8 and Last Day were longer than the Day 1 session (**Fig. 7J, Supp. Table 1**; main effect of Day: F(1.880,90.23)=196.9, P<0.0001, partial ω^2^=0.72 (Large), 95% CI [0.60,0.80]; post hoc information in **Supp. Table 1**) and the number of responses in the sessions on Day 8 and Last Day were less than the Day 1 session (**Fig. 7K, Supp. Table 1**; main effect of Day: F(1.880,90.23)=196.9, P<0.0001, partial ω^2^=0.72 (Large), 95% CI [0.60,0.80]; post hoc information in **Supp. Table 1**). Together these data — longer sessions with fewer responses — suggest all mice underwent Ext learning. As there was no difference in Ext performance across the groups, these data also show no overt effect of 33-GCR or CDDO-EA on Ext.

11.5 mon post-IRR and one day prior to 3-CSI, the movement of mice was collected while they spent 30 min in a novel activity chamber (**Fig. 8**). The four groups of mice moved a similar distance (**Fig. 8A; Supp. Table 1**; main effect of IRR: F(1,50)=4.737, P=0.0343, partial ω^2^=0.06 (Medium), 95% CI [0.00,0.24]; post hoc: all comparisons P>0.05). All groups were also similar in average velocity (**Fig. 8B; Supp. Table 1**). CDDO-EA/33-GCR mice spent on average 16.8% less time ambulating and had 16.8% fewer ambulatory events than CDDO-EA/Sham mice (ambulation time, **Fig. 8C, Supp. Table 1**; main effect of IRR: F(1,50)=7.612, P=0.008, partial ω^2^=0.11 (Medium), 95% CI [0.00,0.30]; post hoc CDDO-EA/Sham vs CDDO-EA/33-GCR: *f* P=0.037, Cohen’s d=0.839 (Large), 95% CI [-1.55,-0.0436]; **Fig. 8D, Supp. Table 1**; IRR: F(1,50)=6.606, P=0.0132, partial ω^2^=0.09 (Medium), 95% CI [0.00, 0.028]; post hoc CDDO-EA/Sham vs CDDO-EA/33-GCR: *f#* P=0.062, Cohen’s d=0.774 (Medium), 95% CI [-1.46,-0.00907]). All four groups spent a similar amount of time rearing **(Fig. 8E, Supp. Table 1)** and reared a similar number of times **(Fig. 8F, Supp. Table 1)**. These data suggest that movement in a novel environment was not grossly impacted by 33-GCR or CDDO-EA, although CDDO-EA in the context of 33-GCR did decrease ambulation metrics vs CDDO-EA in the context of Sham IRR.

**Figure 8.**
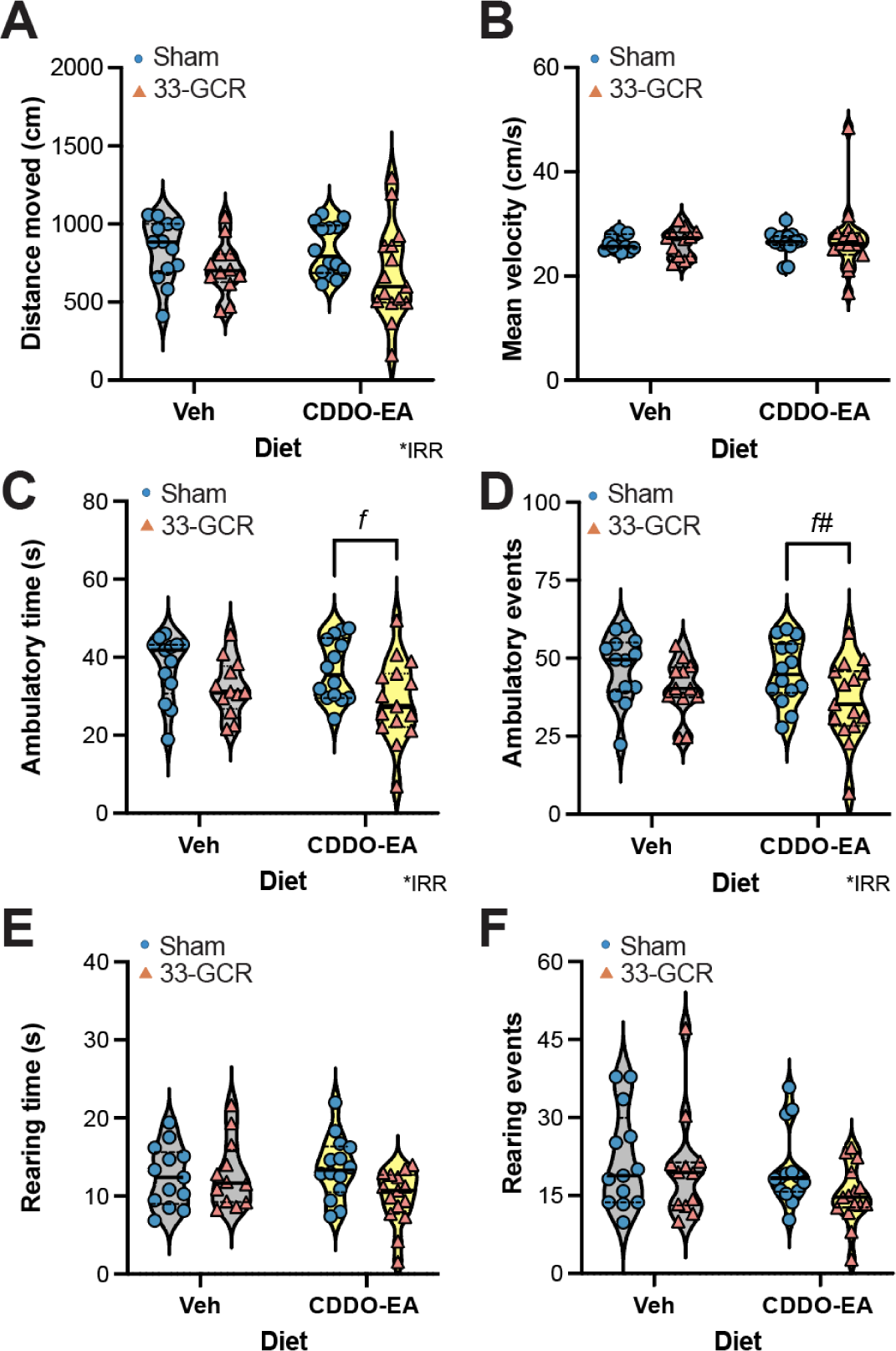
Eleven and one-half mon post-IRR, locomotor activity in a novel environment was grossly similar among groups. (**A-F**) <u>Locomotion metrics</u> for a 30-min session in a novel activity chamber. **(A)** <u>Total distance moved.</u> There was no difference among the IRR groups. **(B)** <u>Mean velocity.</u> All groups had similar average velocity. **(C)** <u>Ambulatory time (seconds, s)</u>. CDDO-EA/33-GCR mice spent 16.8% less time moving than CDDO-EA/Sham mice. **(D)** <u>Ambulatory events.</u> CDDO-EA/33-GCR mice had 16.8% fewer ambulatory events than CDDO-EA/Sham mice. **(E)** <u>Rearing time (s).</u> All groups spent similar time rearing. **(F)** <u>Rearing events.</u> All groups had a similar number of rearing events. **(A-F)** Two-way ANOVA (Diet x IRR). Main effect is denoted by *P<0.05. Bonferroni post-hoc analysis significance denotations between groups: CDDO-EA/Sham vs. CDDO-EA/33-GCR, *f* P<0.05, *f#* 0.08<P<0.05. IRR, irradiation. Veh, vehicle. Violin plots depict median (solid line) and 25% and 75% quartiles (dotted line). Complete data analysis details are provided in **Supp. Table 1.**

During habituation to the novel 3-CSI chamber (**Fig. 9A**), mice in each group showed no preference for either the left or right chamber (**Fig. 9A; Supp. Table 1**; main effect of Chamber: F(1,55)=13.60, P=0.0005, partial ω^2^=0.17 (Large), 95% CI [0.03,0.37]; post hoc: all comparisons P>0.05). Although these mice were all >17-mon-old at the time of the 3-CSI sociability trial (**Fig. 9B**), mice in all groups still spent more time in the chamber with the novel mouse (Stranger 1) vs the chamber with the empty enclosure (**Fig. 9B; Supp. Table 1**; main effect of Chamber: F(1,55)=86.05, P<0.0001, partial ω^2^=0.58 (Large), 95% CI [0.39,0.72]; post hoc: all within group comparisons P≤0.005, all effect sizes Large, all 95%CI exclude 0). This shows all groups of mice, regardless of Diet or IRR, prefer a novel social stimulus more than a novel in animate object. In the final 3-CSI trial, which tests social novelty (**Fig. 9C**), mice in all groups spent similar time near the now-familiar social stimulus (Stranger 1) and a novel social stimulus (Stranger 2, **Fig. 9C, Supp. Table 1**; main effect of Diet: F(1,52)=4.157, P=0.046, partial ω^2^=0.01 (Small), 95% CI [0.00,1.00]; post hoc: all comparisons P>0.05); there is a similar outcome if data are presented as time spent in each chambers rather than time in sniff zone (data not shown). These data show that while social novelty/social memory is deficient in >17-mon-old mice, neither 33-GCR nor CDDO-EA nor the combination change any 3-CSI measures.

**Figure 9.**
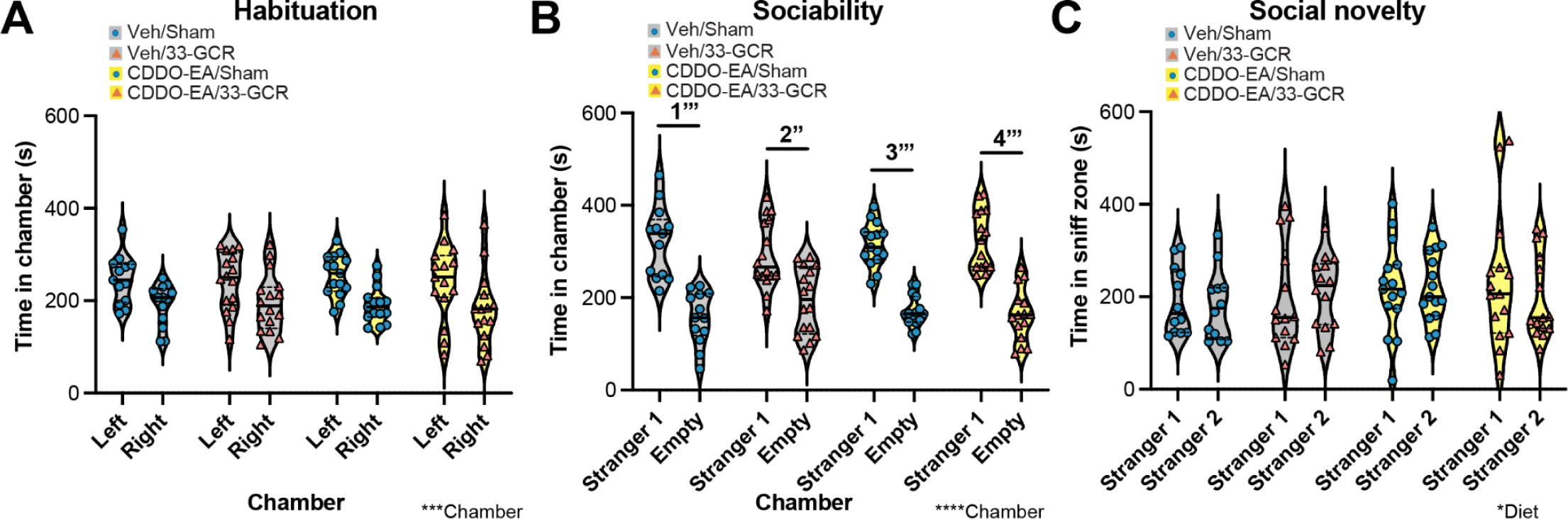
11.75 mon post-IRR, measures in the three-chamber sociability interaction (3-CSI) task are similar among groups. (**A**) <u>3-CSI habituation trial.</u> All groups had similar average time spent in the left or right chamber. **(B)** <u>3-CSI sociability trial.</u> All groups had similar average time spent in the chamber containing a novel conspecific mouse in an enclosure (Stranger 1) and the chamber containing an empty enclosure (Empty). **(C)** <u>3-CSI social novelty trial.</u> All groups had similar average time spent in the sniff zone around a now-familiar conspecific mouse in an enclosure (Stranger 1) and a novel conspecific mouse in the opposite enclosure (Stranger 2). Three-way ANOVA (Chamber x Diet x IRR). Main effects are denoted by *P<0.05, ***P<0.001, or ****P<0.0001. IRR, irradiation. Veh, vehicle. Violin plots depict median (solid line) and 25% and 75% quartiles (dotted line). Complete data analysis details are provided in **Supp. Table 1.**

The first day in the OF (Day 1) when the OF arena is novel, Veh/Sham and Veh/33-GCR mice moved a similar distance, whereas CDDO-EA/33-GCR mice covered on average 14.9% less distance than CDDO-EA/Sham mice (**Fig. 10A, Supp. Table 1**; main effect IRR: F(1,47)=4.673, P=0.0358, partial ω^2^=0.07 (Medium), 95% CI [0.00,0.25]; post hoc: CDDO/Sham vs CDDO/33-GCR: *f* P=0.030, Cohen’s d=0.874 (Large), 95% CI [-1.64,-0.0333]). Mice in the 33-GCR groups also spent less time in the center of the OF (**Fig. 10B; Supp. Table 1**; main effect of IRR: F(1,47)=5.952, P=0.0185, partial ω^2^=0.09 (Medium), 95% CI [0.00,0.28]) and had a lower OF exploration index (**Fig. 10C, Supp. Table 1**; main effect of IRR: F(1,47)=5.831, P=0.0197, partial ω^2^=0.09 (Medium), 95% CI [0.00,0.27]) vs mice in the Sham groups, although there were no post-hoc differences.

**Figure 10.**
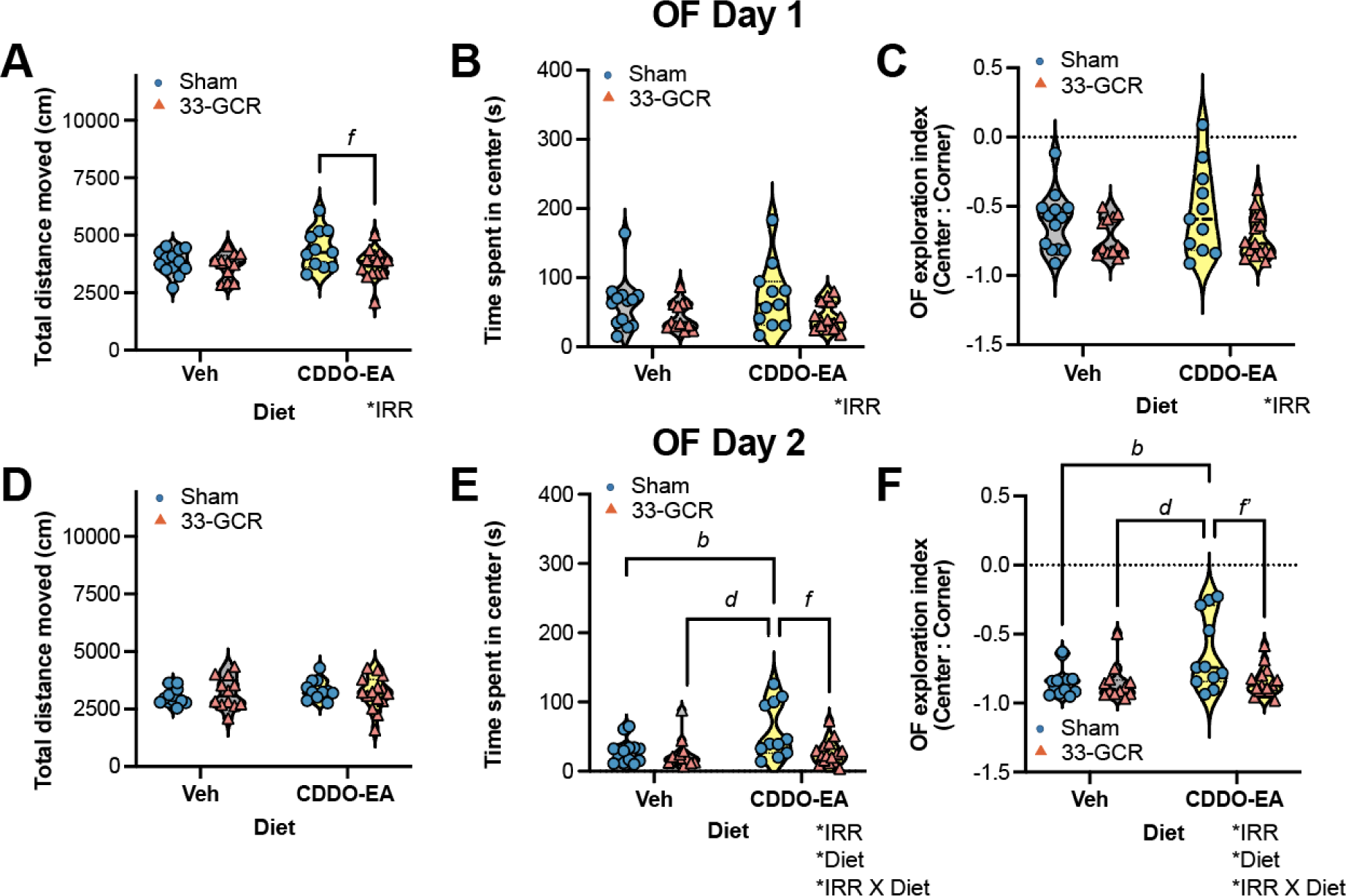
12 mon post-IRR, CDDO-EA/33-GCR mice showed less exploration in a novel, bright open field (OF) relative to CDDO-EA/Sham mice. In contrast, in a familiar OF CDDO-EA/Sham mice showed greater exploration relative to all other groups. (**A-F**) <u>OF metrics on Day 1 **(A-C)** and Day 2 **(D-F)**.</u> **(A)** <u>Day 1, distance moved</u>. CDDO-EA/33-GCR mice moved on average 14.9% less than CDDO-EA/Sham mice. **(B)** <u>Day 1, center time.</u> Mice in the 33-GCR groups spent less time in the center than mice in the Sham group. **(C)** <u>Day 1, OF exploration index</u>. Mice in the 33-GCR groups also had a lower exploration index than mice in the Sham groups. **(D)** <u>Day 2, distance moved</u>. All groups moved a similar average distance on Day 2. **(E)** <u>Day 2, center time.</u> CDDO-EA/Sham mice spent 51-56% more time in the center relative to Veh/Sham, Veh/33-GCR, and CDDO-EA/33-GCR mice. **(F)** <u>Day 2, OF exploration index</u>. CDDO-EA/Sham mice had a higher OF exploration index relative to Veh/Sham, Veh/33-GCR, and CDDO-EA/33-GCR mice. Two-way ANOVA (Diet x IRR). Main effects are denoted by *P<0.05. Bonferroni post-hoc analysis significance denotations between groups: Veh/Sham vs. CDDO-EA/Sham, *b* P<0.05. CDDO-EA/Sham vs. CDDO-EA/33-GCR, *d* P<0.05. CDDO-EA/Sham vs. CDDO-EA/33-GCR, *f#* 0.08<P<0.05, *f* P<0.05. IRR, irradiation. Veh, vehicle. Violin plots depict median (solid line) and 25% and 75% quartiles (dotted line). Complete data analysis details are provided in **Supp. Table 1.**

On the second day of OF (Day 2), when the arena is now familiar, the total distance moved was similar among all four groups (**Fig. 10D, Supp. Table 1**). However, two additional measures from OF Day 2 were notable. In regard to time in the center, CDDO-EA/Sham mice spent 104-129% more time in the center vs Veh/Sham, Veh/33-GCR mice, and CDDO-EA/33-GCR mice (**Fig. 10E, Supp. Table 1**; interaction of IRR x Diet: F(1,47)=4.130, P=0.0478, partial ω^2^=0.06 (Medium), 95% CI [0.00,0.23]; post-hoc: Veh/Sham vs CDDO-EA/Sham: *b* P=0.026, Cohen’s d=1 (Large), 95% CI [0.178,1.84]; Veh/33-GCR vs CDDO-EA/Sham: *d* P=0.013, Cohen’s d=1.04 (Large), 95% CI [0.117,1.82]; CDDO-EA/Sham vs CDDO-EA/33-GCR: *f* P=0.012, Cohen’s d=1.09 (Large), 95% CI [-1.96,-0.258]). OF exploration index followed the same pattern; CDDO-EA/Sham mice had a higher OF exploration index vs Veh/Sham, Veh/33-GCR mice, and CDDO-EA/33-GCR mice (**Fig. 10F, Supp. Table 1**; interaction of IRR x Diet: F(1,47)=4.958, P=0.0308, partial ω^2^=0.07 (Medium), 95% CI [0.00,0.26]; post-hoc: Veh/Sham vs CDDO-EA/Sham: *b* P=0.012, Cohen’s d=0.209 (Small), 95% CI [0.0609,0.392]; Veh/33-GCR vs CDDO-EA/Sham: *d* P=0.010, Cohen’s d=0.217 (Small), 95% CI [0.0627,0.394]; CDDO-EA/Sham vs CDDO-EA/33-GCR: *f’* P=0.009, Cohen’s d=0.209 (Small), 95% CI [0.0636, 0.39]). These data from OF Day 2 suggest that CDDO-EA induces an anti-anxiety-like effect which is negated by 33-GCR.

Habituation (adaptation to a novel environment) is important evolutionarily. We examined habituation and anxiety-relevant metrics 12 mon post-IRR by comparing OF on Day 1 and Day 2. For distance moved, two of the four groups moved less on Day 2 vs Day 1, suggesting normal habituation: Veh/Sham mice (75.6% less on Day 2) and CDDO-EA/Sham mice (69% less on Day 2; **Supp.** Fig. 2A**, Supp. Table 1**; main effect of Day: F(1,47)=45.71, P<0.0001, partial ω^2^=0.25 (Large), 95% CI [0.06,0.46]; post-hoc: Veh/Sham Day 1 vs Day 2: *1’’* P=0.007, Cohen’s d=1.65 (Large), 95% CI [-2.66,-0.591]; CDDO-EA/Sham Day 1 vs Day 2: *3’’’* P=0.0003, Cohen’s d=1.55 (Large), 95% CI [-2.35,-0.671]). For time in the center, only Veh/Sham mice spent 53.7% less time in the center on Day 2 vs Day 1 (**Supp.** Fig. 2B**, Supp. Table 1**; main effect of Day: F(1,47)=27.88, P<0.0001, partial ω^2^=0.12 (Medium), 95% CI [0.00,0.32]; post-hoc: Veh/Sham Day 1 vs Day 2: *1’* P=0.001, Cohen’s d=1.15 (Large), 95% CI [-1.83,-0.35]). For OF exploration index, only Veh/Sham mice had a 37.6% lower OF exploration index on Day 2 vs Day 1 (**Supp.** Fig. 2C**, Supp. Table 1**; main effect of Day: F(1,47)=44.90, P<0.0001, partial ω^2^=0.14 (Large), 95% CI [0.01, 0.35]; post-hoc: Veh/Sham Day 1 vs Day 2: *1’’’* P<0.0001, Cohen’s d=1.38 (Large), 95% CI [-2.17,-0.571]). These data show that Veh/Sham mice and CDDO-EA/Sham mice habituate normally to a new environment, while Veh/33-GCR and CDDO-EA/33-GCR mice do not. In contrast, only Veh/Sham mice are also less anxious Day 2 vs Day 1; CDDO-EA/Sham, Veh/33-GCR, and CDDO-EA/33-GCR mice show similar levels of anxiety-like behavior on both days.

Mice in all four groups faced with a novel and familiar object spent more time exploring the novel object (**Fig. 11A, Supp. Table 1**; interaction of Object x Diet: F(1,56)=4.330, P=0.0420, partial ω^2^=0.02 (Small), 95% CI [0.00,0.14]; post-hoc: Veh/Sham, Familiar vs Novel: *1’’’* P<0.0001, Cohen’s d=1.57 (Large), 95% CI [0.678,2.56]; Veh/33-GCR, Familiar vs Novel: *2’’* P<0.0001, Cohen’s d=2.33 (Large), 95% CI [1.17,3.47]; CDDO-EA/Sham, Familiar vs Novel: *3’’’* P<0.0001, Cohen’s d=2.21 (Large), 95% CI [1.32,3.15]; and CDDO-EA/33-GCR, Familiar vs Novel: *4’’’* P<0.0001, Cohen’s d=1.96 (Large), 95% CI [1.06,2.67]). All groups also showed a similar discrimination index, favoring the novel object (**Fig. 11B, Supp. Table 1**). Taken together, these data suggest that all four groups show preference for a novel vs familiar object.

**Figure 11.**
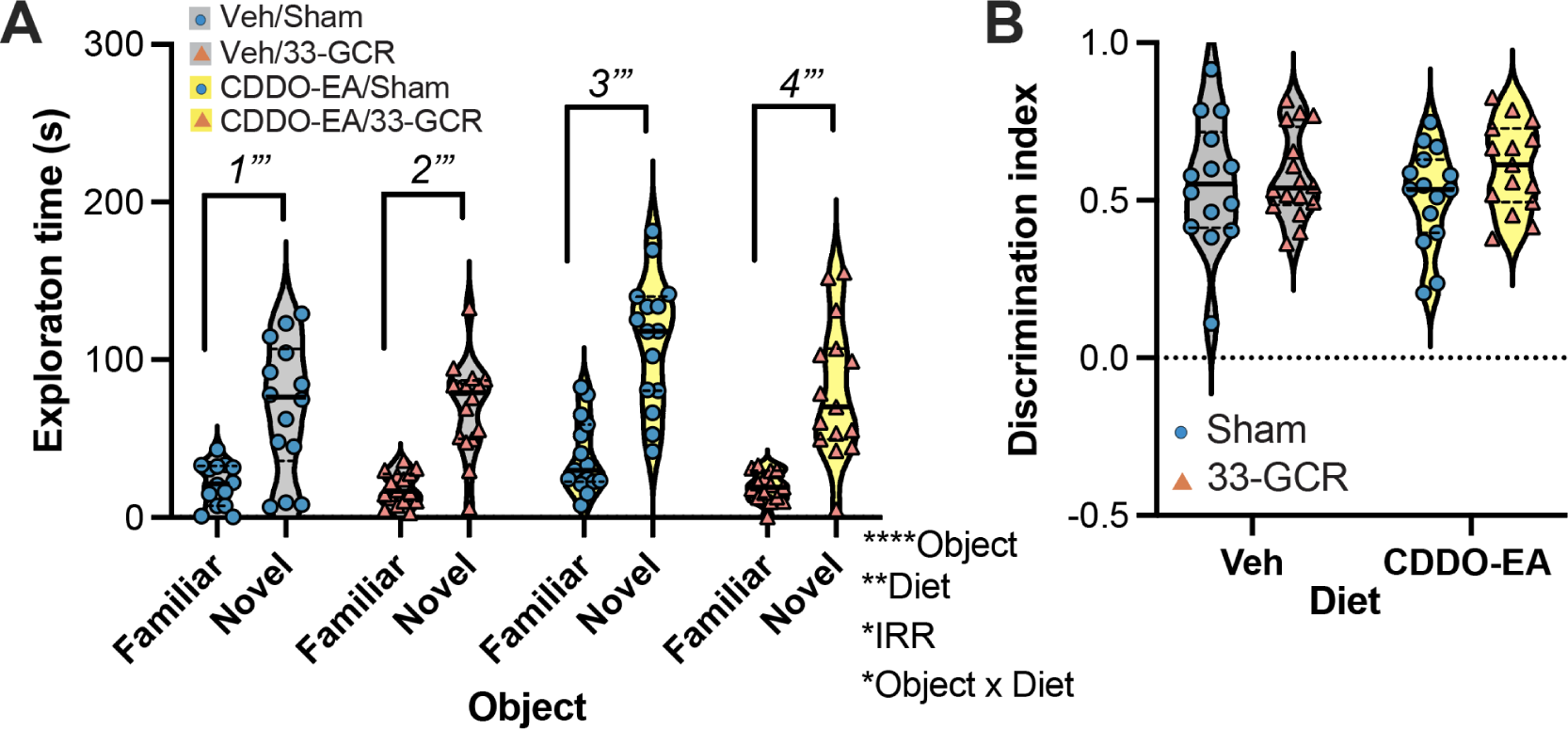
12.5 mon post-IRR, mice in all groups can recognize novel objects, and they discriminate between novel and familiar objects to a similar extent. (**A**) <u>Time spent exploring a novel vs familiar object</u>. All groups spent more time exploring the novel vs familiar object. **(B)** <u>Object discrimination index.</u> All groups have a discrimination index above zero, indicating preference for spending time with a novel vs familiar object. **(A)** Three-way ANOVA (Object x Diet x IRR). Main effects and interactions are denoted by *P<0.05, ****P<0.0001. Bonferroni post-hoc analysis significance denotations between objects for a given group: *1*=within Veh/Sham, *2*=within Veh/33-GCR; *3*=within CDDO-EA/Sham; *4*=within CDDO-EA/33-GCR with *’’*=P<0.001 and *’’*’=P<0.0001. **(B)** Two-way ANOVA (Diet x IRR). IRR, irradiation. s, seconds. Veh, vehicle. Violin plots depict median (solid line) and 25% and 75% quartiles (dotted line). Complete data analysis details are provided in **Supp. Table 1.**

In the marble burying test, all four groups of mice buried a similar number of marbles (**Fig. 12A; Supp. Table 1**). In the nestlet shredding test, 33-GCR groups had marginally lower weight of intact nestlet vs Sham groups, but there was no post-hoc difference (**Fig. 12B; Supp. Table 1**; main effect of IRR: F(1,56)=6.094, P=0.016, partial ω^2^=0.08 (Medium), 95% CI [0.00,0.25]). These data suggest equivalent performance in tests of compulsive behavior in all four groups.

**Figure 12.**
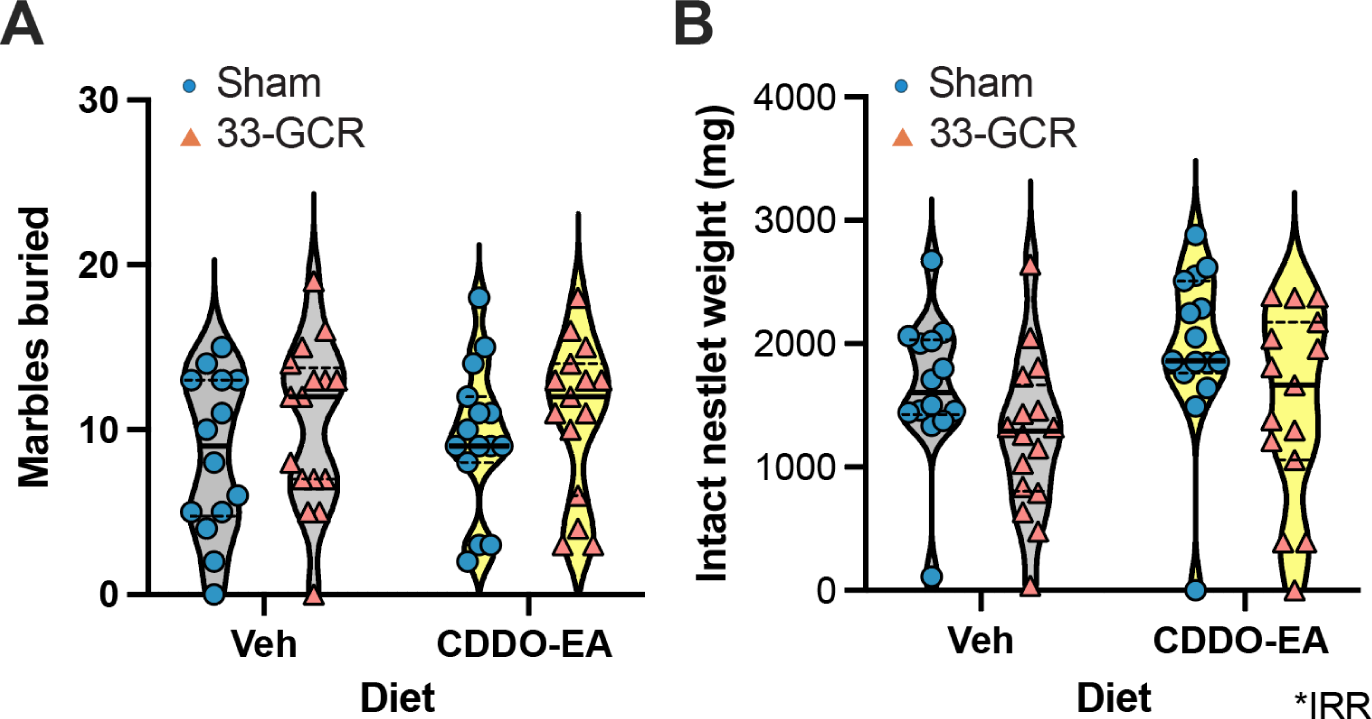
13 mon post-IRR, all groups of mice show similar measures in tests of compulsive behavior. (**A**) <u>Number of marbles buried.</u> Mice in all four groups buried a similar number of marbles. **(B)** <u>Weight of intact nestlet</u>. Mice in the 33-GCR groups had marginally greater intact nestlets vs mice in Sham groups. **(A-B)** Two-way ANOVA (Diet x IRR). IRR, irradiation. Veh, vehicle. Violin plots depict median (solid line) and 25% and 75% quartiles (dotted line). Complete data analysis details are provided in **Supp. Table 1.**

After brains were collected (14.25 mon post-IRR) and processed for IHC, all mice showed evidence of adult-generated neurons, showing DCX+ immature neurons in the subgranular zone of the dentate gyrus granule cell layer. DCX+ immature neurons had a characteristic pyramidal soma with processes of varying lengths emerging from the soma (**Fig. 13A**). When DCX+ immature neuron number was examined as a single value per mouse, Veh/33-GCR mice had 27.4% fewer DCX+ immature neurons vs Veh/Sham mice. However, there was no difference in DCX+ immature neurons in CDDO-EA/33-GCR and CDDO-EA/Sham mice (**Fig. 13B; Supp. Table 1**; main effect of IRR: F(1,40)=6.749, P=0.0131, partial ω^2^=0.12 (Medium), 95% CI [0.00,0.33]; post-hoc: Veh/Sham vs Veh/33-GCR, *a* P=0.0339, Cohen’s d=1.42 (Large), 95% CI [-2.35,-0.407]). When DCX+ immature neuron number was examined in across the anterior-posterior axis of the dentate gyrus, DCX+ immature neuron number was lower in Veh/33-GCR vs Veh/Sham mice at one anterior distance from bregma (bregma –2.02; **Fig. 13C, Supp. Table 1**; main effect of Treatment: F(3,480)=4.036, P=0.0075, partial ω^2^=0.06 (Medium), 95% CI [0.02,0.11]; post-hoc at Bregma –2.02: Veh/Sham vs Veh/33-GCR: *a* P=0.0372, Cohen’s d=1.21 (Large), 95% CI [-1.81,-0.488]) and lower in CDDO-EA/33-GCR vs CDDO-EA/Sham mice at a distinct, more posterior distance from bregma (bregma –3.22; **Fig. 13C, Supp. Table 1**; post-hoc at Bregma –3.22: CDDO/Sham vs CDDO/33-GCR: *f* P=0.0247, Cohen’s d=0.885 (Large), 95% CI [-1.43,-0.111]). In addition to DCX+ immature neurons, all mice showed DCX+ progenitor cells with their irregular soma and no processes emerging/seen in the Z plane of each cell (**Fig. 13D**). Unlike DCX+ immature neuron number, there was no difference among the four groups of mice in DCX+ progenitor cell number either when considered as a single value per mouse (**Fig. 13E, Supp. Table 1**) or examined across the anterior-posterior coronal axis (**Fig. 13F, Supp. Table 1**; main effect of Bregma: F(11,480)=71.85, P<0.0001, partial ω^2^=0.60 (Medium), 95% CI [0.53,0.64]). These data show that 33-GCR decreases DCX+ immature neuron number but not progenitor cell number relative to Sham mice.

**Figure 13.**
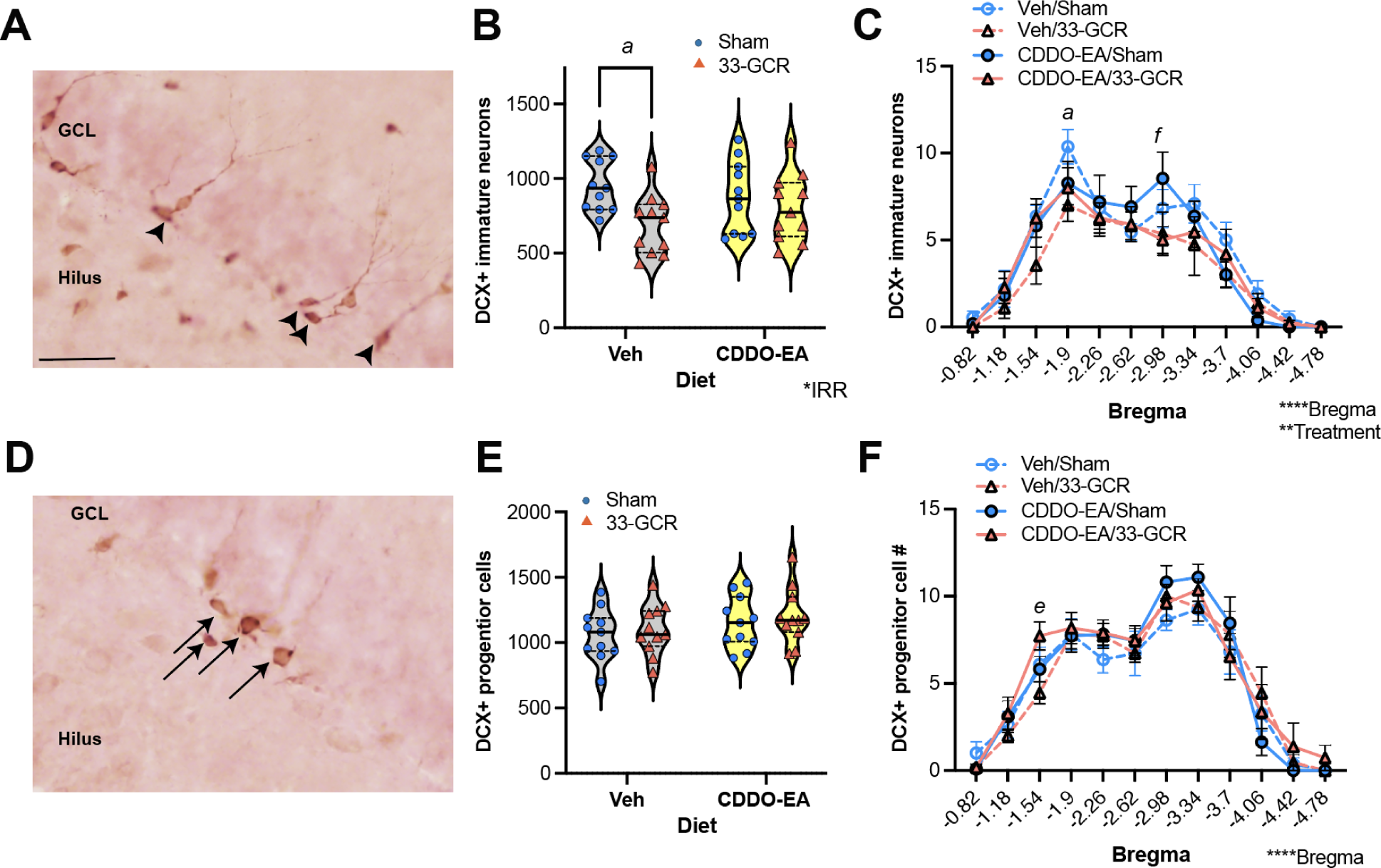
14.5 mon post-IRR, Veh/33-GCR mice have fewer doublecortin-immunoreactive (DCX+) immature neurons in the dentate gyrus (DG) subgranular zone (SGZ) vs Veh/Sham mice, but all groups have a similar number of DCX+ progenitors. (**A-C**) <u>DCX+ immature neurons</u> and (D-F) <u>DCX+ progenitor cells in the DG SGZ.</u> (A) <u>Representative photomicrograph of DCX+ immature neurons</u>. Arrowheads indicate brown-stained DCX+ immature neuron soma in the SGZ at the border of the DG granule cell layer (GCL) and the DG hilus. (B) <u>Number of DCX+ immature neurons represented as a single value per mouse.</u> There are 27.4% fewer DCX+ immature neurons in the DG SGZ of Veh/33-GCR mice vs Veh/Sham mice. (C) <u>Number of DCX+ immature neurons represented as a single value for each distance from bregma.</u> There are fewer DCX+ immature neurons in Veh/33-GCR mice vs Veh/Sham mice at bregma –2.02, and fewer DCX+ immature neurons in CDDO-EA/33-GCR vs CDDO-EA/Sham at bregma –3.22. (D) <u>Representative photomicrograph of DCX+ progenitor cells</u>. Arrows indicate brown-stained DCX+ progenitor cell soma in the SGZ. (E) <u>Number of DCX+ progenitor cells represented as a single value per mouse.</u> All values were similar in the four groups of mice. (F) <u>Number of DCX+ progenitor cells represented as a single value for each distance from bregma.</u> All values were similar in the four groups of mice. (B, E) Two-way ANOVA (Diet X IRR). Main effects and interactions are denoted by *P<0.05. Bonferroni post-hoc analysis significance denotations between groups: Veh/Sham vs Veh/33-GCR, *a* P<0.05, CDDO/Sham vs CDDO/GCR, *f* P<0.05. (C, F) Two-way ANOVA (Bregma x Treatment). Bonferroni post-hoc. +, immunoreactive. Scale bar in (A)=100um, applies to (A, D). IRR, irradiation. Veh, vehicle. Violin plots (B, E) depict median (solid line) and 25% and 75% quartiles (dotted line). Complete data analysis details are provided in **Supp. Table 1**.

## 4. DISCUSSION

To understand the potential risks of 33-GCR to the CNS of female mice, we assessed spaceflight-relevant behavioral and cognitive domains 2 to 14.25 months after administration of CDDO-EA or Veh chow and whole-body exposure to 75cGy (33-GCR) or Sham IRR. Mice were tested on translationally relevant paradigms on an operant touchscreen platform as well as arena-based behavioral tests, and their hippocampal neurogenesis was also examined post-mortem (**Fig. 1**). The data we present are multi-faceted and provide a long-term perspective of how exposure to 33-GCR influences cognition and behavior in female mice, and how administration of the antioxidant CDDO-EA counters 33-GCR-induced changes. Below we discuss the main changes that emerge from this longitudinal study and integrate these new findings into the existing space radiation and cognition literature.

In the touchscreen platform, one of the most notable cognitive changes we report is that Veh/33-GCR mice are slower to acquire a simple stimulus-response association when tested ∼9-mon post-IRR (**Fig. 7C**). These mice can learn the task within 30-min (data not shown) but appear simply slower in reaching the time cut-off often used for this task (finishing within 15-min). Supporting their slower completion, when the entire 30-min of each session is considered, Veh/33-GCR mice have longer sessions (**Fig. 7D**) and longer latencies to retrieve the reward (**Fig. 7F**). Veh/33-GCR mice have, however, normal latency to make a correct touch (**Fig. 7E**) and near-perfect accuracy (**Fig. 7G**). One way to interpret these data is that in this simple stimulus-response association task, Veh/33-GCR mice know what to do and they do it correctly, but they take a longer time to retrieve the reward. This relative slowness could be due to motoric issues, motivation issues, or cognitive issues. The latter is unlikely, as the near-perfect accuracy suggests the relative slowness is not due to cognitive dysfunction. Given that Veh/33-GCR mice make fewer beam breaks and have longer session lengths and latencies to collect the reward, we believe Veh/33-GCR mice have motoric issues or motivation issues or a combination. Attention may also play a role in these 33-GCR effects; male mice exposed to a slightly lower dose of 33-GCR (40-49 cGy vs. our 75 cGy) have worse attentional processes when tested ∼6-mon post-IRR (Desai *et al*. 2023). However, prior work in the simple stimulus-response learning task suggests that one of the metrics (correct touch latency) reflects attentional processes (Swan *et al*. 2014), and radiation does not change this metric in our mice in the present study (**Fig. 7E**). Therefore, we do not favor the interpretation that at this time point 33-GCR mice have disrupted attention. Other work in male mice showed no change in reward sensitivity in male mice collected ∼6-mon post-IRR (Desai *et al*. 2023), which could be considered to be reflective of reward motivation. It is possible that there are motivation deficits in our mice due to a higher dose of 33-GCR used here (75 cGy) relative to this prior work (50 cGy), or that motivation effects are different at distinct time points post-IRR. Future work is needed to dissect if the changes in Veh/33-GCR mice we show here are due to an impact on motor or motivational processes, test difficulty, or a combination. On a related note, we also report administration of CDDO-EA at the time of IRR (9-mon prior) prevented the 33-GCR-induced decrease in reward collection latency (**Fig. 7F**), perhaps normalizing reward motivation in these female mice.

Another point of discussion about the touchscreen platform cognitive data we present is the <u>lack</u> of change seen in female mice in behavioral pattern separation (Small separation, **Fig. 4**), a mission-relevant task reliant on the integrity of the hippocampal dentate gyrus. This is notable since our prior work showed male and female C57BL/6J mice exposed to a single high-energy particle regimen (i.e. ^56^Fe, ∼20 cGy total) performed *<u>better</u>* than Sham IRR mice in a behavioral pattern separation touchscreen task (LDR) 6-8 mon post-IRR (Whoolery *et al*. 2020; Soler *et al*. 2021). In our current work, female C57BL/6J mice were examined 3.5-9 mon after 33-GCR and showed no gross difference in LDR performance (**Fig. 4**). Thus, we conclude that this dose (∼75cGy total in 1 day) of whole-body 33-GCR does not influence behavioral pattern separation assessed via LDR, at least at this time examined post-IRR. In contrast, other work shows that female C57BL/6J mice that received either acute or repeated exposure to 33-GCR (∼50 cGy total over 24 days) had *<u>worse</u>* behavioral pattern separation relative to control female mice, as assessed in the OUL (object in updated location) test (Alaghband *et al*. 2023); the authors state such changes did not “manifest” in the male mice. This prior work and our current 33-GCR work cannot be directly compared, since the distinct behavioral pattern separation tests used (LDR in ours, OUL in theirs) may have different levels of sensitivity to dysfunction. However, clearly more work is warranted on the impact of 33-GCR on behavioral pattern separation in female and male mice. Our present negative LDR finding aside, we also present two interesting CDDO-EA LDR findings. First, in measures of discrimination (Large separation, **Fig. 4B, F, J**), female mice that received CDDO-EA at the time of 33-GCR had *<u>better</u>* dentate gyrus-dependent discrimination vs. CDDO-EA/Sham IRR mice, as shown by greater accuracy and shorter session length. In fact, this leads to the second finding: female mice that received CDDO-EA had *<u>better</u>* behavioral pattern separation (Small separation, **Fig. 4D, H, L**) than Veh mice, regardless of IRR (33-GCR or Sham IRR). It is tempting to conclude that CDDO-EA may improve cognition in the long-term. However, much more work would be needed prior to making such a conclusion, including addressing the possibility that prior behavior testing influenced performance. To this end, one approach would be to test parallel groups of mice in distinct cognitive tasks at the same time point after CDDO-EA administration.

The arena-based analyses we performed also provided several points for discussion, the first being the lack of difference we report between female 33-GCR and Sham IRR mice in OF locomotion (**Fig. 10**). A few studies have established that single particle type exposures lead to fine motor deficits in rats (Blackwell *et al*. 2021), but differences in gross locomotion are scarce in the literature with some studies reporting that post-IRR OF locomotion is increased (Sorokina *et al*. 2020; Kokhan *et al*. 2019; Liu *et al*. 2019; Boutros *et al*. 2021), decreased (Allen *et al*. 2015; Holshouser *et al*. 2020), or unchanged (Impey *et al*. 2023). Time post-IRR and dose may be important in seeing 5-ion (1H, 2 x 1,000 MeV/n; 4He, 400 MeV/n; 16O, 400 MeV/n; 28Si, 600 MeV/n) GCR-induced changes in locomotion (Puukila *et al*. 2023), although there are conflicting results (Boutros *et al*. 2021). The potential impact of time post-IRR on locomotion was one reason we examined this metric at several time points post-IRR. Prior work exposing mice to 33-GCR have not reported differences in locomotion, neither in male mice recorded in activity chambers (75cGy) nor in male or female mice recorded in an OF (acute 40cGy or 50cGy across 4 weeks) (Kiffer *et al*. 2021; Schaeffer *et al*. 2022). In keeping with this recent 33-GCR literature, we also report no radiation-dependent differences in gross locomotion in female mice receiving 75cGy 33-GCR in either the activity chambers (**Fig. 8**) or in an OF (**Fig. 10**). Curiously, when the environment was novel (in the activity chambers and OF arena on Day 1), female CDDO-EA/33-GCR mice spend less time moving than CDDO-EA/Sham mice. However, when the environment was familiar (in the touchscreen chambers for habituation for Acq and OF arena on Day 2), there was no difference in movement between CDDO-EA/Sham and CDDO-EA/33-GCR mice. This implies the effect of CDDO-EA on locomotor activity is more likely to be seen in a novel environment, a fact that may be useful in future study of this potential countermeasure.

There are many other arena-based tests in which we report no difference after 33-GCR and/or CDDO-EA, including sociability, social novelty, anxiety/exploratory drive, habituation learning, object recognition memory, and compulsive-like behavior. Some of these are notable negative findings. For example, we find sociability was intact regardless of radiation or diet; we did not observe preference for social novelty in any group, likely due to the advanced age at the time of 3-CSI (16 mon-of-age). Also, the lack of effect on object recognition memory in our current work is striking given that the most commonly reported behavioral deficit after whole-body charged-particle radiation is seen in novel object recognition (NOR) assessments (Cacao and Cucinotta 2019). However, prior work with female mice also shows intact NOR 3.5 mon after chronic (50cGy) or acute (40 cGy) 33-GCR; notably, male mice in that study had disrupted NOR (Alaghband et al. 2023). Thus, future work should test if 33-GCR-induced deficits in NOR are indeed sex-specific.

In addition to the behavioral and cognitive data we present, we also assessed the number of DCX+ cells in the dentate gyrus subgranular zone, an index of adult neurogenesis. Radiation exposure has long been associated with detrimental effects on proliferative cells, including those in the hippocampal dentate gyrus. Monoenergetic single particle radiation paradigms lower indices of dentate gyrus neurogenesis (Rivera *et al*. 2013; DeCarolis *et al*. 2014; Whoolery *et al*. 2017; Zanni *et al*. 2018; Whoolery *et al*. 2020; Rola *et al*. 2008; Rola *et al*. 2005; Casadesus *et al*. 2005; Rola *et al*. 2004; Simmons *et al*. 2022). However, our data presented here are the first to assess the impact of 33-GCR, and thus the first to assess if multiparticle exposure at a mission-relevant dose can change the number of adult-generated dentate gyrus neurons. We find that female mice 14.25 mon post-33-GCR have fewer immature neurons than Sham IRR mice, with progenitor cell number unchanged. Previously, dentate gyrus neurogenesis was considered crucial for behavioral pattern separation in young adult rodents (Clelland *et al*. 2009; Sahay *et al*. 2011; Nakashiba *et al*. 2012). However, here we show that behavioral pattern separation (measured 6-mon prior) is similar to control values even with decreased number of new hippocampal neurons *postmortem*. Our findings add to the growing evidence that the number of new neurons does not always predict pattern separation ability in aged mice (Whoolery *et al*. 2020; Saxe *et al*. 2007; Cès *et al*. 2018).

There are at least two limitations of the present study. First, the dose rate for delivery of radiation is not reflective of what would happen in space travel. Our experimental model relies on an acute 75cGy radiation dose delivered within 1.5hrs, whereas in spaceflight, there is a constant 0.5mGy/day exposure behind standard spacecraft shielding during solar maximum (Zeitlin *et al*. 2013). A recent study comparing an acute 40cGy exposure to a protracted 50cGy 33-GCR exposure (24 fractions across 4 weeks) points to dose-rate effects in working memory, anxiety, and social interaction, though the unbalanced dose design could also point to dose-dependence (Alaghband *et al*. 2023). Second, the potential countermeasure CDDO-EA here was delivered over a few days whereas any future use in space travel would likely require a longer dosing period which may exert different pharmacological protective effects than described here. We recommend that future work assess GCR risks at varying doses and dose rates, along with chronic administration of potential countermeasures.

In summary, mature female C57BL/6J mice exposed to 33-GCR had worse stimulus-response learning and fewer DCX+ immature neurons relative to control mice over a year post-IRR, yet normal behavioral pattern separation, anxiety-like behavior, exploration, object recognition, and habituation. As for the potential countermeasure CDDO-EA, one radiation effect/CDDO-EA effect emerged: CDDO-EA prevented 33-GCR-induced deficit in stimulus response learning. There were also CDDO-EA effects: CDDO-EA/Sham and CDDO-EA/33-GCR mice had better behavioral pattern separation vs. their respective control groups (Veh/Sham, Veh/33-GCR), and CDDO-EA/33-GCR mice had better cognitive flexibility (reversal number) vs. Veh/33-GCR. Our study implies space radiation is a risk to a female crew’s longitudinal mission-relevant cognitive processes and CDDO-EA is a potential dietary countermeasure for space-radiation CNS risks.

## SUPPLEMENTAL FIGURE LEGENDS

**Supplemental Figure 1.**
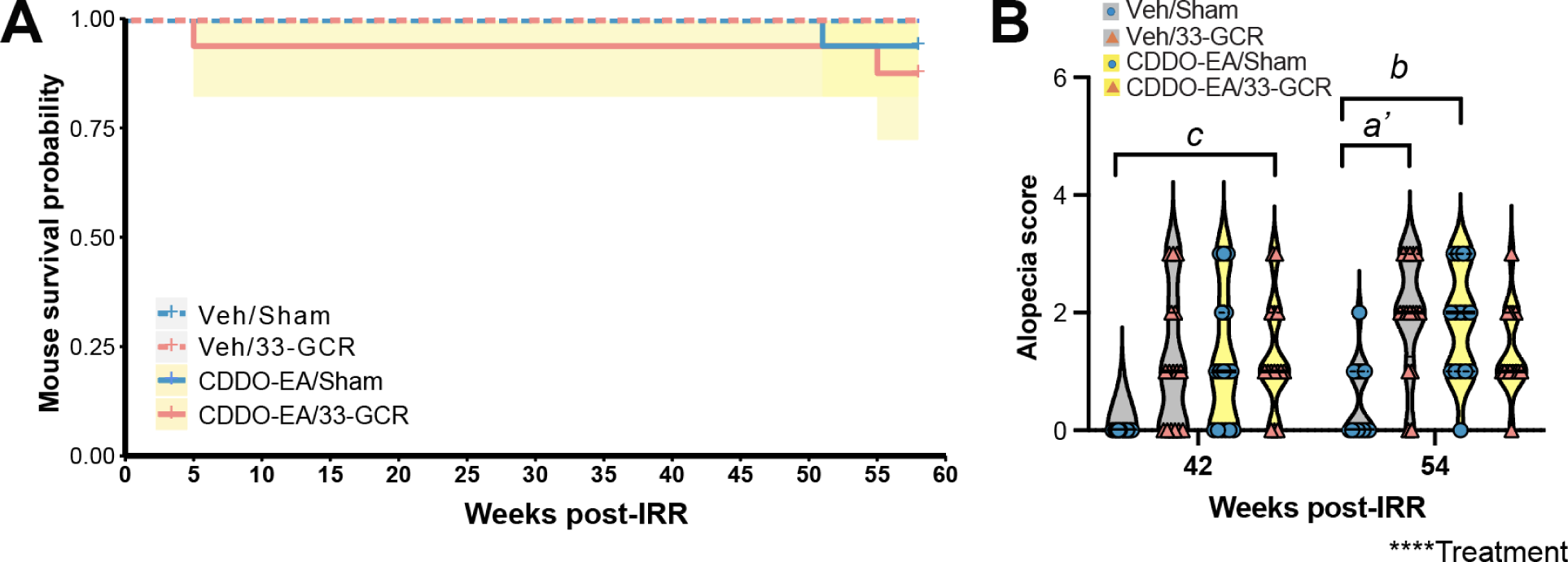
33-GCR and CDDO-EA did not negatively influence survival of mice but did influence coat score. (**A**) <u>Survival curve</u> of mice of the course of the experiment showed no difference in mouse survival across the four groups. **(B)** <u>Alopecia (coat score)</u> of mice 10.5 mon (42 weeks) and 13.5 mon (54 weeks) post-IRR showed at 42 weeks post-IRR, CDDO/33-GCR mice had a higher alopecia score (worse coat state) vs Veh/Sham mice. At 54 weeks post-IRR, Veh/33-GCR and CDDO-EA/Sham mice had higher alopecia scores than Veh/Sham mice (P<0.01 and P<0.05, respectively). Statistical analysis in **A**: Log-Rank, P>0.05 and **B**: Kruskal-Wallis (Treatment), P<0.0001, post-hoc analysis: Veh/Sham vs Veh/33-GCR, a’ P<0.01; Veh/Sham vs CDDO-EA/Sham, *b* P<0.05; Veh/Sham vs CDDO-EA/33-GCR: *c* P<0.05. 14-16 mice per group. Violin plots depict median (solid line) and 25% and 75% quartiles (dotted line). Complete data analysis details are provided in **Supplemental Table 1 (Supp. Table 1)**.

**Supplemental Figure 2.**
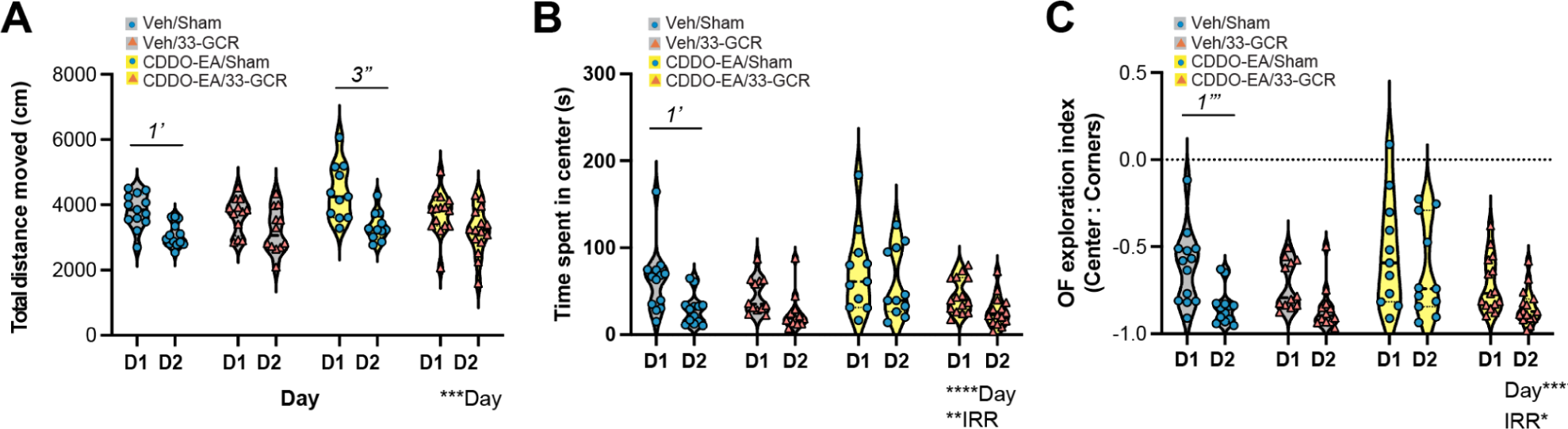
12 mon post-IRR, Veh/Sham and CDDO-EA/Sham mice show habituation to a novel, bright environment, but Veh/33-GCR and CDDO-EA/33-GCR mice do not, and only Veh/Sham mice show less anxiety-like behavior on Day 2 vs Day 1. (**A**) <u>Distance moved on subsequent open field (OF) days.</u> Veh/Sham and CDDO-EA/Sham mice show habituation (moving 75.6% and 68.6% less, respectively, on OF Day 2 vs Day 1) to a novel, bright environment; Veh/33-GCR and CDDO-EA/33-GCR mice do not. **(B)** <u>Center time on subsequent OF days.</u> Veh/Sham mice are the only group that spends less time (53.7% less) in the Center on Day 2 vs Day 1. **(C)** <u>OF exploration index on subsequent OF days.</u> Veh/Sham mice are also the only group that has a lower OF exploration index (27.3% lower) on Day 2 vs Day 1. Main dotted line in **(C)** depicts the zero point of OF exploration index: negative values indicating more exploring time in corners relative to center. Violin plots depict median (solid line) and 25% and 75% quartiles (dotted line). Complete data analysis details are provided in **Supp. Table 1.**

## Abbreviations

33-beam-GCR: 33-beam galactic cosmic radiation
3-CSI: 3-chamber social interaction
AALAC: Association for Assessment and Accreditation of Laboratory Animal Care
Acq: Acquisition
BLAF: BLAF
BNL: Brookhaven National Laboratories
CDDO-EA: 2-cyano-3, 12-dioxooleana-1, 9-dien-28-oic acid-ethylamide
CNS: central nervous system
DCX: doublecortin
DCX+: doublecortin-immunoreactive
EPM: elevated plus maze
Ext: Extinction
GCR: galactic cosmic radiation
h: hour(s)
Hab: habituation
IACUC: Institutional Animal Care and Use Committee
IHC: immunohistochemistry
IRR: irradiation
LDR: location discrimination reversal
LDR Test: location discrimination reversal testing
LDR Train: location discrimination reversal training
Min: minute(s)
mon: month(s)
MT: Must Touch
NOR: novel object recognition
NSRL: NASA Space Radiation Laboratory
OF: open field
PFC: prefrontal cortex
PI: punish incorrect
ROUT: robust outlier testing
s: second(s)
S-: non-rewarded stimulus
S+: rewarded stimulus
SGZ: subgranular zone
Veh: vehicle

## Acknowledgements

We thank all the team members at BNL/NSRL for their help before, during, and after the 19A irradiation campaign, particularly Adam Rusek and Peter Guida. This work was supported by funds to Sanghee Yun (a 2018 PENN McCabe Pilot grant, a 2019 IBRO travel grant, a 2019 NARSAD Young Investigator Grant from the Brain and Behavior Research Foundation, a 2020 PENN Undergraduate Research Foundation grant, 2021 NASA HERO grant 80NSSC21K0814 and Augmentation award, a 2022 Foerderer Fund for Excellence Award, two CHOP Junior Faculty Awards (2021, Bhoj; 2023, Van Batavia), and NIH awards MH076690, MH107945, and MH129970), Fred Kiffer (Translational Research Institute for Space Health [TRISH] through NASA cooperative agreement NNX16AO69A, Penn Provost/CHOP Postdoctoral Fellowship for Academic Diversity, Perelman School of Medicine’s Department of Radiation Oncology Pilot Grant, Amelia Eisch (NIH MH129970; NS007413, DA007290, DA023555, DA016765, MH107945; NASA NNX07AP84G, NNX12AB55G, NNX15AE09G), and Jerry Shay (NNX16AE08G and NNX15AI21G). Sanghee Yun and Amelia Eisch were also supported by NIH DK135871 (PI: SA Zderic), NIH NS088555 (PI: AM Stowe) and NIH MH117628 (PI: Lambert). Amelia Eisch received further support from NIH NS126279 (PI: Ahrens-Nicklas). Catalina Guzman and Ivan Soler were supported by PennPREP (R25 GM071745, PI: KL Jordan-Sciutto), Harley Haas was supported by the 2021 Penn Undergraduate Research Mentoring Program (PURM), the the 2022 Summer Undergraduate Internship Program (SUIP), 2023 Penn College Alumni Society Board of Managers and President’s Undergraduate Research Grant, Jaysen Lara-Jiménez was supported by SUIP. YH was supported by the Undergraduate Translational Research Immersion Program sponsored by PENN Institute for Translational Medicine and Therapeutics. Special thanks to an anonymous donor who has provided support for the Eisch Lab and this project.

## Pre-Submission Author Contributions (for details and explanations see CRediT)

Listed in alphabetical order by last name:

Conceptualization: AJE, SY, JWS, FCK

Data curation: SY, FCK, GLB, HAH, IS, CSG, RP, AJE

Formal analysis: AJE, SY, FCK, HAH, GLB, RP, JLJ, PLK

Funding acquisition: AJE, SY, JWS, FCK

Investigation: SY, FCK, CSG, IS, GLB, HAH, RS, PLK, FHT, KJA, KL

Methodology: SY, FCK, CSG, IS, KL, JWS, AJE

Project administration: SY, FCK, JWS, AJE

Resources: AJE, SY, JWS

Software/Code: RS, YR, RP

Supervision: AJE, SY, FCK, JWS

Validation: HAH, RP, JLJ, PLK, RS

Visualization: SY, FCK

Writing, original draft: FCK, SY, AJE

Writing, review & editing: FCK, SY, JWS, AJE

## Data availability statement

Data are available on reasonable written request.

