## Supplementary table 1 for "The longitudinal behavioral effects of acute exposure to galactic cosmic radiation in female C57BL/6J mice: implications for deep space missions, female crews, and potential antioxidant countermeasures"

**Supplementary Table 1. Statistics for Yun and Kiffer et al.**

| Experiment | Measure | Test Phase | Figure Panel | Group | Sample Size (n) | Mean (predicted or rank) or Median | Gaussian (vs. LogNormal) | Test Statistic | Main Effect or Interaction (bold text: P<0.05) | Dist.-Value (Dfn, Dfd) | Main Effect P-Value (bold text: P<0.05) | % of Total Variation | Effect Size: Partial $\omega^2$ when two-way ANOVA p<0.05. ≥0.01 small; ≥0.06 medium; ≥0.13 large; Partial eta square, when Kruskal-Wallis p<0.05. ≤0.01 small, ≤0.06 medium, and | Can effect size be considered different than zero? | Post-hoc Test Correction | Group Difference P-Value (bold text: P<0.05). Lower case italicized letter (e.g. *a, b, c,* etc) indicates specific group differences. Apostrophe indicates gradations of P values. Post-hoc values of 0.05<P<0.08 are only considered notable (and indicated below and on graph by #) when two additional conditions are met: if effect size is medium or large and if post-hoc 95% CI does not include zero | Effect Size: Cohen's d for parametric multiple comparison and Cliff's delta for non-parametric mulitple comparison (Lenhard, W. & Lenhard, A. (2016), Ho et al., (2019); determined only for P<0.08, | 95% CI | Can post-hoc effect size be considered different than zero? |
|---|---|---|---|---|---|---|---|---|---|---|---|---|---|---|---|---|---|---|---|
| Animal Weights | Weight (Long-term) | | 1B | Veh/Sham | 16-14 | | Yes | Mixed-Effects 3-way Repeated Measures ANOVA | IRR | F (1,56) = 2.53 | P=0.117 | | | | Bonferroni | | | | |
| | | | | Veh/33-GCR | 16 | | Yes | | Diet | F (1,56) = 5.095 | **P=0.028** | | ges = 0.07 (Medium) | | | | | | |
| | | | | CDDO-EA/Sham | 16-15 | | Yes | | Time | F (1,56) = 1.043 | **P<0.0001** | | ges = 0.371 (Large) | | | | all *b* and *b′* are large (>0.8) or medium (>0.73) | | |
| | | | | CDDO-EA/33-GCR | 16-14 | | Yes | | IRR x Diet | | P=0.312 | | | | | Veh/Sham vs CDDO-EA/Sham: *b* **P<0.05**, *b′* **P<0.01** | | | |
| | | | | | | | | | IRR x Time | F (4.35, 243.64) = 2.692 | **P=0.028** | | ges = 0.009 (Small) | | | CDDO-EA/Sham vs CDDO-EA/33-GCR: *f* **P<0.05** | all *f* are medium (0.73< f <0.78) | | |
| | | | | | | | | | Diet x Time | F (4.35, 243.64) = 3.710 | **P<0.001** | | ges = 0.012 (Small) | | | | | | |
| | | | | | | | | | IRR x Diet x Time | F (4.35, 243.64) = 0.637 | P=0.65 | | | | | | | | |
| Animal Weights | 14.25 mon post-IRR | | 1C | Veh/Sham | 14 | 41.35 | Yes | 2-way ANOVA | IRR x Diet | F (1, 55) = 0.7659 | P=0.385 | 1.207 | | | | | | | |
| | | | | Veh/33-GCR | 16 | 42.44 | Yes | | Diet | F (1, 55) = 5.057 | **P=0.028** | 7.971 | partial $\omega^2$ = 0.06 (Medium), 95% CI [0.00, 0.23] | No | Bonferroni | Veh/Sham vs CDDO-EA/Sham: *b*# P=0.0652 | 0.885 (Large) | -1.43, -0.111 | Yes |
| | | | | CDDO-EA/Sham | 15 | 36.18 | Yes | | IRR | F (1, 55) = 2.348 | P=0.131 | 3.701 | | | | | | | |
| | | | | CDDO-EA/33-GCR | 14 | 40.16 | Yes | | | | | | | | | | | | |
| | General Touchscreen Train Habituation 1 | Total Beam breaks | 2A | Veh/Sham | 14 | 196.1 | Yes | 2-way ANOVA | IRR x Diet | F (1, 57) = 1.518 | P=0.223 | 2.471 | | | | NA | | | |
| | | | | Veh/33-GCR | 16 | 192.1 | Yes | | Diet | F (1, 57) = 0.3063 | P=0.582 | 0.498 | | | | | | | |
| | | | | CDDO-EA/Sham | 16 | 203.4 | Yes | | IRR | F (1, 57) = 2.597 | P=0.112 | 4.227 | | | | | | | |
| | | | | CDDO-EA/33-GCR | 15 | 172.9 | Yes | | | | | | | | | | | | |
| General Touchscreen Training | General Touchscreen Train Days to Criteria | Habituation 1 (set as 1 day so Days to Criteria not analyzed) | 2B | Veh/Sham | 14 | 1 | No | NA | | | | | | | | NA | | | |
| | | | | Veh/33-GCR | 16 | 1 | No | | | | | | | | | | | | |
| | | | | CDDO-EA/Sham | 16 | 1 | No | | | | | | | | | | | | |
| | | | | CDDO-EA/33-GCR | 16 | 1 | No | | | | | | | | | | | | |
| | | Habituation 2 (set as 1 day so Days to Criteria not analyzed) | | Veh/Sham | 14 | 1 | No | NA | | | | | | | | NA | | | |
| | | | | Veh/33-GCR | 16 | 1 | No | | | | | | | | | | | | |
| | | | | CDDO-EA/Sham | 16 | 1 | No | | | | | | | | | | | | |
| | | | | CDDO-EA/33-GCR | 16 | 1 | No | | | | | | | | | | | | |
| | | Initial Touch | | Veh/Sham | 14 | 1.143 | No | Kruskal-Wallis | | | | | | | | NA | | | |
| | | | | Veh/33-GCR | 16 | 1.125 | No | | | | P=0.419 | | | | | | | | |
| | | | | CDDO-EA/Sham | 16 | 1.063 | No | | | | | | | | | | | | |
| | | | | CDDO-EA/33-GCR | 16 | 1.333 | No | | | | | | | | | | | | |
| | | Must Touch | | Veh/Sham | 14 | 6.643 | Yes | 2-way ANOVA | IRR x Diet | F (1, 57) = 0.09092 | P=0.764 | 0.129 | | | | Veh/Sham vs CDDO-EA/Sham: *b* **P=0.015** | 0.851 (Large) | -1.43, -0.119 | Yes |
| | | | | Veh/33-GCR | 16 | 6.063 | No | | Diet | F (1, 57) = 13.15 | **P=0.0006** | 18.71 | partial $\omega^2$ = 0.17 (Large), 95% CI [0.03, 0.36] | Yes | Bonferroni | | | | |
| | | | | CDDO-EA/Sham | 16 | 3.813 | Yes | | IRR | F (1, 57) = 0.2539 | P=0.616 | 0.361 | | | | Veh/33-GCR vs CDDO/33-GCR: *e* **P=0.041** | 1.09 (Large) | -1.72, -0.372 | Yes |
| | | | | CDDO-EA/33-GCR | 16 | 3.667 | Yes | | | | | | | | | | | | |
| | | Must Initiate | | Veh/Sham | 14 | 1.429 | No | Kruskal-Wallis | | | | | | | | NA | | | |
| | | | | Veh/33-GCR | 16 | 4.813 | No | | | | P=0.616 | | | | | | | | |
| | | | | CDDO-EA/Sham | 16 | 1.188 | No | | | | | | | | | | | | |
| | | | | CDDO-EA/33-GCR | 16 | 1.2 | No | | | | | | | | | | | | |
| | | Punish Incorrect | | Veh/Sham | 14 | 15.71 | Yes | 2-way ANOVA | IRR x Diet | F (1, 57) = 0.1227 | P=0.727 | 0.191 | | | | | | | |
| | | | | Veh/33-GCR | 16 | 15.69 | No | | Diet | F (1, 57) = 6.838 | **P=0.011** | 10.68 | partial $\omega^2$ = 0.09 (Medium), 95% CI [0.00, 0.26] | Yes | Bonferroni | Veh/33-GCR vs CDDO-EA/33-GCR: P=0.077 | 0.637 (Medium) | -1.14, 0.00977 | No |
| | | | | CDDO-EA/Sham | 16 | 11.63 | Yes | | IRR | F (1, 57) = 0.1333 | P=0.716 | 0.208 | | | | | | | |
| | | | | CDDO-EA/33-GCR | 16 | 10.33 | Yes | | | | | | | | | | | | |
| | Punish Incorrect | Trials to Criterion | 2C | Veh/Sham | 14 | 442.1 | Yes | 2-way ANOVA | IRR x Diet | F (1, 57) = 0.5344 | P=0.467 | 0.778 | | | | | | | |
| | | | | Veh/33-GCR | 16 | 444.9 | No | | Diet | F (1, 57) = 10.79 | **P=0.001** | 15.71 | partial $\omega^2$ = 0.14 (Large), 95% CI [0.02, 0.32] | Yes | Bonferroni | Veh/33-GCR vs CDDO-EA/33-GCR: e **P=0.011** | 0.845 (Large) | -1.27, -0.273 | Yes |
| | | | | CDDO-EA/Sham | 16 | 316.4 | Yes | | IRR | F (1, 57) = 0.4564 | P=0.502 | 0.664 | | | | | | | |
| | | | | CDDO-EA/33-GCR | 15 | 247.2 | Yes | | | | | | | | | | | | |
| | Punish Incorrect | Blank Touches | 2D | Veh/Sham | 14 | 148.8 | Yes | 2-way ANOVA | IRR x Diet | F (1, 57) = 0.735 | P=0.394 | 1.067 | | | | | | | |
| | | | | Veh/33-GCR | 16 | 158.1 | No | | Diet | F (1, 57) = 10.98 | **P=0.001** | 15.95 | partial $\omega^2$ = 0.14 (Large), 95% CI [0.02, 0.33] | Yes | Bonferroni | Veh/33-GCR vs CDDO-EA/33-GCR: *e′* **P=0.008** | 0.885 (Large) | -1.32, -0.244 | Yes |
| | | | | CDDO-EA/Sham | 16 | 101.4 | Yes | | IRR | F (1, 57) = 0.141 | P=0.708 | 0.205 | | | | | | | |
| | | | | CDDO-EA/33-GCR | 15 | 77.67 | Yes | | | | | | | | | | | | |
| | Punish Incorrect | Correct Touch Latency | 2E | Veh/Sham | 14 | 9.829 | Yes | 2-way ANOVA | IRR x Diet | F (1, 57) = 0.731 | P=0.395 | 1.054 | | | | | | | |
| | | | | Veh/33-GCR | 16 | 9.104 | Yes | | Diet | F (1, 57) = 11.71 | **P=0.001** | 16.86 | partial $\omega^2$ = 0.15 (Large), 95% CI [0.02, 0.34] | Yes | Bonferroni | Veh/Sham vs CDDO-EA/Sham: *b′* **P=0.008** | 1.07 (Large) | -1.88, -0.257 | Yes |
| | | | | CDDO-EA/Sham | 16 | 7.601 | Yes | | IRR | F (1, 57) = 0.286 | P=0.594 | 0.413 | | | | | | | |
| | | | | CDDO-EA/33-GCR | 15 | 7.768 | No | | | | | | | | | | | | |
| LDR Train | % subject to complete | | 3B | Veh/Sham | 14 | | | Log-rank (Mantel-Cox) | | $\chi^2$ = 4.064 | P=0.254 | | | | | NA | | | |
| | | | | Veh/33-GCR | 16 | | | | | | | | | | | | | | |
| | | | | CDDO-EA/Sham | 16 | | | | | | | | | | | | | | |
| | | | | CDDO-EA/33-GCR | 15 | | | | | | | | | | | | | | |
| | Days to Completion | | 3C | Veh/Sham | 14 | 18.54 | Yes | 2-way ANOVA | IRR x Diet | F (1, 56) = 4.286 | **P=0.043** | 7.044 | partial $\omega^2$ = 0.05 (Small), 95% CI [0.00, 0.21] | No | | all comparisons: P>0.05 | | | |
| | | | | Veh/33-GCR | 16 | 27.19 | Yes | | Diet | F (1, 56) = 0.1138 | P=0.737 | 0.187 | | | | | | | |
| | | | | CDDO-EA/Sham | 16 | 24 | Yes | | IRR | F (1, 56) = 0.4544 | P=0.503 | 0.746 | | | | | | | |
| | | | | CDDO-EA/33-GCR | 15 | 19.6 | Yes | | | | | | | | | | | | |
| | Total Trials on Last Training Day | | 3D | Veh/Sham | 14 | 46.21 | Yes | 2-way ANOVA | IRR x Diet | F (1, 57) = 1.506 | P=0.224 | 2.02 | | | | | | | |
| | | | | Veh/33-GCR | 16 | 41.75 | Yes | | Diet | F (1, 57) = 11.90 | **P=0.001** | 15.96 | partial $\omega^2$ = 0.15 (Large), 95% CI [0.02, 0.34] | Yes | Tukey | Veh/33-GCR vs CDDO-EA/33-GCR: *e′* **P=0.003** | 1.14 (Large) | 0.478, 1.88 | Yes |
| | | | | CDDO-EA/Sham | 16 | 49.44 | Yes | | IRR | F (1, 57) = 3.425 | P=0.069 | 4.593 | | | | | | | |
| | | | | CDDO-EA/33-GCR | 15 | 48.53 | Yes | | | | | | | | | | | | |
| | Final Session Length | | 3E | Veh/Sham | 14 | 1576 | Yes | 2-way ANOVA | IRR x Diet | F (1, 57) = 0.056 | P=0.813 | 0.0842 | | | | | | | |
| | | | | Veh/33-GCR | 16 | 1674 | Yes | | Diet | F (1, 57) = 7.293 | **P=0.009** | 10.93 | partial $\omega^2$ = 0.09 (Medium), 95% CI [0.00, 0.27] | Yes | Tukey | all comparisons: P>0.05 | | | |
| | | | | CDDO-EA/Sham | 16 | 1427 | Yes | | IRR | F (1, 57) = 1.959 | P=0.167 | 2.936 | | | | | | | |
| | | | | CDDO-EA/33-GCR | 15 | 1497 | No | | | | | | | | | | | | |
| | Last Day % Correct | | 3F | Veh/Sham | 14 | 53.32 | No | 2-way ANOVA | IRR x Diet | F (1, 57) = 0.540 | P=0.465 | 0.906 | | | | NA | | | |
| | | | | Veh/33-GCR | 16 | 54.47 | Yes | | Diet | F (1, 57) = 1.940 | P=0.169 | 3.254 | | | | | | | |
| | | | | CDDO-EA/Sham | 16 | 59.59 | Yes | | IRR | F (1, 57) = 0.120 | P=0.730 | 0.201 | | | | | | | |
| | | | | CDDO-EA/33-GCR | 15 | 56.41 | Yes | | | | | | | | | | | | |
| | | Block 1 | | Veh/Sham | 14 | 67.19 | Yes | | | | | | | | | | | | |
| | | | | Veh/33-GCR | 16 | 57.91 | Yes | | | | | | | | | | | | |

**Supplementary Table 1. Statistics for Yun and Kiffer et al.**

| Experiment | Measure | Test Phase | Figure Panel | Group | Sample Size (n) | Mean (predicted or rank) or Median | Gaussian (vs. LogNormal) | Test Statistic | Main Effect or Interaction (bold text: P<0.05) | Dist.-Value (Dfn, Dfd) | Main Effect P-Value (bold text: P<0.05) | % of Total Variation | Effect Size: Partial ω^2 when two-way ANOVA p<0.05. ≥0.01 small; ≥0.06 medium; ≥0.13 large; Partial eta square, when Kruskal-Wallis p<0.05. ≤0.01 small, ≤0.06 medium, and | Can effect size be considered different than zero? | Post-hoc Test Correction | Group Difference P-Value (bold text: P<0.05). Lower case italicized letter (e.g. *a, b, c,* etc) indicates specific group differences. Apostrophe indicates gradations of P values. Post-hoc values of 0.05<P<0.08 are only considered notable (and indicated below and on graph by #) when two additional conditions are met: if effect size is medium or large and if post-hoc 95% CI does not include zero | Effect Size: Cohen's d for parametric multiple comparison and Cliff's delta for non-parametric mulitple comparison (Lenhard, W. & Lenhard, A. (2016), Ho et al., (2019); determined only for P<0.08, | 95% CI | Can post-hoc effect size be considered different than zero? |
|---|---|---|---|---|---|---|---|---|---|---|---|---|---|---|---|---|---|---|---|
| | | Block 1 | | CDDO-EA/Sham | 16 | 67.5 | Yes | | | | | | | | | | | | |
| | | | | CDDO-EA/33-GCR | 15 | 65.59 | Yes | | | | | | | | | | | | |
| | | Block 2 | | Veh/Sham | 14 | 66.08 | Yes | | | | | | | | | | | | |
| | | | | Veh/33-GCR | 16 | 60.37 | Yes | | | | | | | | | | | | |
| | | | | CDDO-EA/Sham | 16 | 68.62 | Yes | | | | | | | | | | | | |
| | | | | CDDO-EA/33-GCR | 15 | 65.91 | Yes | | | | | | | | | | | | |
| | | Block 3 | | Veh/Sham | 14 | 63.8 | Yes | | Block x Treatment | F (12, 228) = 1.638 | P=0.082 | 5.459 | | | | | | | |
| | % Correct to 1st Reversal Large Separation | | 4B | Veh/33-GCR | 16 | 59.5 | Yes | Repeated Measures 2-way ANOVA | Block | F (3.835, 218.6) = 0.781 | P=0.533 | 0.868 | | | Tukey | Veh/33-GCR vs CDDO-EA/33-GCR in Block 5: *e* **P=0.048** | 0.979 (Large) | 0.285, 1.51 | Yes |
| | | | | CDDO-EA/Sham | 16 | 73.32 | Yes | | Treatment | F (3, 57) = 3.079 | **P=0.034** | 4.245 | partial ω^2 = 0.17 (Large), 95% CI [0.08, 0.25] | Yes | | | | | |
| | | | | CDDO-EA/33-GCR | 15 | 63.12 | No | | | | | | | | | | | | |
| | | Block 4 | | Veh/Sham | 14 | 52.9 | Yes | | | | | | | | | | | | |
| | | | | Veh/33-GCR | 16 | 63.04 | Yes | | | | | | | | | | | | |
| | | | | CDDO-EA/Sham | 16 | 62.3 | Yes | | | | | | | | | | | | |
| | | | | CDDO-EA/33-GCR | 15 | 67.55 | No | | | | | | | | | | | | |
| | | Block 5 | | Veh/Sham | 14 | 62.72 | Yes | | | | | | | | | | | | |
| | | | | Veh/33-GCR | 16 | 59.26 | No | | | | | | | | | | | | |
| | | | | CDDO-EA/Sham | 16 | 63.64 | Yes | | | | | | | | | | | | |
| | | | | CDDO-EA/33-GCR | 15 | 74.77 | Yes | | | | | | | | | | | | |
| | | Block 1 | | Veh/Sham | 14 | 55.04 | Yes | | | | | | | | | | | | |
| | | | | Veh/33-GCR | 16 | 53.72 | Yes | | | | | | | | | | | | |
| | | | | CDDO-EA/Sham | 16 | 55.91 | Yes | | | | | | | | | | | | |
| | | | | CDDO-EA/33-GCR | 15 | 55.97 | Yes | | | | | | | | | | | | |
| | | Block 2 | | Veh/Sham | 14 | 44.2 | Yes | | | | | | | | | | | | |
| | | | | Veh/33-GCR | 16 | 51.03 | Yes | | Block x Treatment | F (12, 228) = 1.837 | **P=0.043** | 6.411 | partial ω^2 = 0.11 (Medium), 95% CI [0.01, 0.15] | Yes | | | | | |
| | | | | CDDO-EA/Sham | 16 | 49.84 | Yes | | Block | F (3.872, 220.7) = 1.587 | P=0.180 | 1.846 | | | | | | | |
| | | | | CDDO-EA/33-GCR | 15 | 67.32 | Yes | | Treatment | F (3, 57) = 3.493 | **P=0.021** | % Var = 3.975 | partial ω^2 = 0.17 (Large), 95% CI [0.08, 0.25] | Yes | | | | | |
| | % Correct to 1st Reversal Small Separation | Block 3 | 4D | Veh/Sham | 14 | 54.28 | Yes | Repeated Measures 2-way ANOVA | Subject | F (57, 228) = 1.304 | P=0.089 | % Var = 21.62 | | | Tukey | Veh/Sham vs CDDO-EA/33-GCR in Block 2: *c′* **P=0.005**<br>Veh/Sham vs CDDO-EA/Sham in Block 5: *b* **P=0.021**<br>Veh/Sham vs CDDO-EA/33-GCR in Block 5: *c*' **P=0.008** | 1.36 (Large)<br>1.11 (Large)<br>1.32 (Large) | 0.59, 2.1<br>0.326, 1.83<br>0.474, 2.06 | Yes<br>Yes<br>Yes |
| | | | | Veh/33-GCR | 16 | 55.71 | Yes | | | | | | | | | | | | |
| | | | | CDDO-EA/Sham | 16 | 57.58 | Yes | | | | | | | | | | | | |
| | | | | CDDO-EA/33-GCR | 15 | 56.52 | Yes | | | | | | | | | | | | |
| | | Block 4 | | Veh/Sham | 14 | 61.91 | Yes | | | | | | | | | | | | |
| | | | | Veh/33-GCR | 16 | 50.24 | Yes | | | | | | | | | | | | |
| | | | | CDDO-EA/Sham | 16 | 60.53 | Yes | | | | | | | | | | | | |
| | | | | CDDO-EA/33-GCR | 15 | 58.78 | Yes | | | | | | | | | | | | |
| | | Block 5 | | Veh/Sham | 14 | 40.76 | Yes | | | | | | | | | | | | |
| | | | | Veh/33-GCR | 16 | 44.28 | Yes | | | | | | | | | | | | |
| | | | | CDDO-EA/Sham | 16 | 60.53 | Yes | | | | | | | | | | | | |
| | | | | CDDO-EA/33-GCR | 15 | 57.7 | Yes | | | | | | | | | | | | |
| | % Correct to 1st Reversal Large Separation Block 1 | | 4E | Veh/Sham | 14 | 55.04 | Yes | 2-way ANOVA | IRR x Diet | F (1, 57) = 0.024 | P=0.875 | 0.0434 | | | | NA | | | |
| | | | | Veh/33-GCR | 16 | 53.72 | Yes | | Diet | F (1, 57) = 0.126 | P=0.723 | 0.2211 | | | | | | | |
| | | | | CDDO-EA/Sham | 16 | 55.91 | Yes | | IRR | F (1, 57) = 0.020 | P=0.886 | 0.0358 | | | | | | | |
| | | | | CDDO-EA/33-GCR | 15 | 55.97 | Yes | | | | | | | | | | | | |
| | % Correct to 1st Reversal Large Separation Block 5 | | 4F | Veh/Sham | 14 | 62.72 | Yes | 2-way ANOVA | IRR x Diet | F (1, 57) = 3.372 | P=0.071 | 5.14 | | | | | | | |
| | | | | Veh/33-GCR | 16 | 59.26 | No | | Diet | F (1, 57) = 4.283 | **P=0.043** | 6.529 | partial ω^2 = 0.05 (Small), 95% CI [0.00, 0.20] | No | Bonferroni | Veh/33-GCR vs CDDO-EA/33-GCR: *e* **P=0.043** | 0.979 (Large) | 0.285, 1.51 | Yes |
| | | | | CDDO-EA/Sham | 16 | 63.64 | Yes | | IRR | F (1, 57) = 0.934 | P=0.337 | 1.425 | | | | | | | |
| | | | | CDDO-EA/33-GCR | 15 | 74.77 | Yes | | | | | | | | | | | | |
| LDR Test | % Correct to 1st Reversal Small Separation Block 1 | | 4G | Veh/Sham | 14 | 55.04 | Yes | 2-way ANOVA | IRR x Diet | F (1, 57) = 0.024 | P=0.875 | 0.043 | | | | NA | | | |
| | | | | Veh/33-GCR | 16 | 53.72 | Yes | | Diet | F (1, 57) = 0.126 | P=0.723 | 0.221 | | | | | | | |
| | | | | CDDO-EA/Sham | 16 | 55.91 | Yes | | IRR | F (1, 57) = 0.020 | P=0.886 | 0.035 | | | | | | | |
| | | | | CDDO-EA/33-GCR | 15 | 55.97 | Yes | | | | | | | | | | | | |
| | % Correct to 1st Reversal Small Separation Block 5 | | 4H | Veh/Sham | 14 | 40.76 | Yes | 2-way ANOVA | IRR x Diet | F (1, 57) = 0.551 | P=0.460 | 0.76 | | | Bonferroni | | | | |
| | | | | Veh/33-GCR | 16 | 44.28 | Yes | | Diet | F (1, 57) = 15.08 | **P=0.0003** | 20.78 | partial ω^2 = 0.19 (Large), 95% CI [0.04, 0.38] | Yes | | Veh/Sham vs CDDO-EA/Sham: *b′* **P=0.004** | 1.11 (Large) | 0.326, 1.83 | Yes |
| | | | | CDDO-EA/Sham | 16 | 60.53 | Yes | | IRR | F (1, 57) = 0.006 | P=0.936 | 0.009 | | | | Veh/33-GCR vs CDDO-EA/33-GCR: *e#* P=0.057 | 0.866 (Large) | 0.0538, 1.64 | Yes |
| | | | | CDDO-EA/33-GCR | 15 | 57.7 | Yes | | | | | | | | | | | | |
| | Time to 1st Reversal Large Separation Block 1 | | 4I | Veh/Sham | 14 | 698.5 | No | 2-way ANOVA | IRR x Diet | F (1, 57) = 0.378 | P=0.540 | 0.588 | | | | NA | | | |
| | | | | Veh/33-GCR | 16 | 1029 | No | | Diet | F (1, 57) = 2.066 | P=0.156 | 3.21 | | | | | | | |
| | | | | CDDO-EA/Sham | 16 | 600 | Yes | | IRR | F (1, 57) = 4.589 | **P=0.036** | 7.132 | partial ω^2 = 0.06 (Medium), 95% CI [0.00, 0.021] | No | | | | | |
| | | | | CDDO-EA/33-GCR | 15 | 783.1 | Yes | | | | | | | | | | | | |
| | Time to 1st Reversal Large Separation Block 5 | | 4J | Veh/Sham | 14 | 678.4 | No | 2-way ANOVA | IRR x Diet | F (1, 57) = 3.538 | P=0.065 | 5.366 | | | | | | | |
| | | | | Veh/33-GCR | 16 | 918.5 | Yes | | Diet | F (1, 57) = 5.201 | **P=0.026** | 7.887 | partial ω^2 = 0.06 (Medium), 95% CI [0.00, 0.23] | No | Bonferroni | Veh/33-GCR vs CDDO-EA/33-GCR: *e′* **P=0.008** | 1.08 (Large) | -1.81, -0.242 | Yes |
| | | | | CDDO-EA/Sham | 16 | 630.6 | No | | IRR | F (1, 57) = 0.01578 | P=0.900 | 0.023 | | | | | | | |
| | | | | CDDO-EA/33-GCR | 15 | 420.5 | No | | | | | | | | | | | | |
| | Time to 1st Reversal Small Separation Block 1 | | 4K | Veh/Sham | 14 | 1141 | No | 2-way ANOVA | IRR x Diet | F (1, 57) = 0.01203 | P=0.913 | 0.021 | | | | | | | |
| | | | | Veh/33-GCR | 16 | 1155 | Yes | | Diet | F (1, 57) = 0.1185 | P=0.731 | 0.207 | | | | | | | |
| | | | | CDDO-EA/Sham | 16 | 1106 | No | | IRR | F (1, 57) = 0.0004 | P=0.983 | 0.0007 | | | | | | | |
| | | | | CDDO-EA/33-GCR | 15 | 1086 | No | | | | | | | | | | | | |
| | Time to 1st Reversal Small Separation Block 5 | | 4L | Veh/Sham | 14 | 1419 | No | 2-way ANOVA | IRR x Diet | F (1, 57) = 0.043 | P=0.836 | 0.062 | | | | | | | |
| | | | | Veh/33-GCR | 16 | 1355 | No | | Diet | F (1, 57) = 12.00 | **P=0.001** | 17.31 | partial ω^2 = 0.15 (Large), 95% CI [0.02, 0.34] | Yes | | Veh/Sham vs CDDO-EA/Sham: *b#* P=0.052 Veh/33-GCR vs CDDO-EA/33-GCR: *e* **P=0.022** | 0.873 (Large) | -1.64, -0.105 | Yes |
| | | | | CDDO-EA/Sham | 16 | 961.3 | Yes | | IRR | F (1, 57) = 0.442 | P=0.508 | 0.638 | | | | | 0.906 (Large) | -1.86, -0.0809 | Yes |
| | | | | CDDO-EA/33-GCR | 15 | 838.6 | No | | | | | | | | | | | | |
| | Trials to 1st Reversal Large Separation | | 4M | Veh/Sham | 14 | 18 | No | 2-way ANOVA | IRR x Diet | F (1, 57) = 0.038 | P=0.845 | 0.063 | | | | NA | | | |
| | | | | Veh/33-GCR | 16 | 23.56 | Yes | | Diet | F (1, 57) = 0.250 | P=0.618 | 0.416 | | | | | | | |
| | | | | CDDO-EA/Sham | 16 | 20.06 | Yes | | IRR | F (1, 57) = 2.823 | P=0.098 | 4.704 | | | | | | | |

Supplementary Table 1. Statistics for Yun and Kiffer et al.

| Experiment | Measure | Test Phase | Figure Panel | Group | Sample Size (n) | Mean (predicted or rank) or Median | Gaussian (vs. LogNormal) | Test Statistic | Main Effect or Interaction (bold text: P<0.05) | Dist.-Value (Dfn, Dfd) | Main Effect P-Value (bold text: P<0.05) | % of Total Variation | Effect Size: Partial ω^2 when two-way ANOVA p<0.05. ≥0.01 small; ≥0.06 medium; ≥0.13 large; Partial eta square, when Kruskal-Wallis p<0.05. ≤0.01 small, ≤0.06 medium, and [illegible] | Can effect size be considered different than zero? | Post-hoc Test Correction | Group Difference P-Value (bold text: P<0.05). Lower case italicized letter (e.g. *a*, *b*, *c*, etc) indicates specific group differences. Apostrophe indicates gradations of P values. Post-hoc values of 0.05<P<0.08 are only considered notable (and indicated below and on graph by #) when two additional conditions are met: if effect size is medium or large and if post-hoc 95% CI does not include zero | Effect Size: Cohen's d for parametric multiple comparison and Cliff's delta for non-parametric mulitple comparison (Lenhard, W. & Lenhard, A. (2016), Ho et al., (2019); determined only for P<0.08, | 95% CI | Can post-hoc effect size be considered different than zero? |
|---|---|---|---|---|---|---|---|---|---|---|---|---|---|---|---|---|---|---|---|
| | Block 1 | | | CDDO-EA/33-GCR | 15 | 24.47 | Yes | | | | | | | | | | | | |
| | Trials to 1st Reversal Large Separation Block 5 | | 4N | Veh/Sham | 14 | 18 | No | 2-way ANOVA | IRR x Diet | F (1, 57) = 3.407 | P=0.070 | 5.458 | | | | NA | | | |
| | | | | Veh/33-GCR | 16 | 19.44 | Yes | | Diet | F (1, 57) = 0.417 | P=0.520 | 0.668 | | | | | | | |
| | | | | CDDO-EA/Sham | 16 | 20.88 | Yes | | IRR | F (1, 57) = 1.552 | P=0.217 | 2.487 | | | | | | | |
| | | | | CDDO-EA/33-GCR | 15 | 13.47 | No | | | | | | | | | | | | |
| | Trials to 1st Reversal Small Separation Block 1 | | 4O | Veh/Sham | 14 | 30.14 | Yes | 2-way ANOVA | IRR x Diet | F (1, 57) = 0.227 | P=0.635 | 0.393 | | | | NA | | | |
| | | | | Veh/33-GCR | 16 | 28.75 | No | | Diet | F (1, 57) = 0.446 | P=0.506 | 0.773 | | | | | | | |
| | | | | CDDO-EA/Sham | 16 | 31 | Yes | | IRR | F (1, 57) = 0.027 | P=0.869 | 0.047 | | | | | | | |
| | | | | CDDO-EA/33-GCR | 15 | 33.87 | Yes | | | | | | | | | | | | |
| | Trials to 1st Reversal Small Separation Block 5 | | 4P | Veh/Sham | 14 | 40.93 | No | 2-way ANOVA | IRR x Diet | F (1, 57) = 1.421 | P=0.238 | 5.458 | | | | NA | | | |
| | | | | Veh/33-GCR | 16 | 29.44 | Yes | | Diet | F (1, 57) = 3.121 | P=0.078 | 0.668 | | | | | | | |
| | | | | CDDO-EA/Sham | 16 | 28.81 | Yes | | IRR | F (1, 57) = 2.684 | P=0.106 | 2.487 | | | | | | | |
| | | | | CDDO-EA/33-GCR | 15 | 27 | No | | | | | | | | | | | | |
| | Block 5 Reversal Number Large Separation | | 5A | Veh/Sham | 14 | 1.857 | Yes | 2-way ANOVA | IRR x Diet | F (1, 57) = 0.861 | P=0.357 | 1.307 | | | | | | | |
| | | | | Veh/33-GCR | 16 | 1.438 | No | | Diet | F (1, 57) = 7.725 | **P=0.007** | 11.71 | partial ω^2 = 0.10 (Medium), 95% CI [0.00, 0.28] | Yes | Bonferroni | Veh/33-GCR vs CDDO-EA/33-GCR: *e* **P=0.021** | 1.15 (Large) | 0.28, 2.11 | Yes |
| | | | | CDDO-EA/Sham | 16 | 2.438 | Yes | | IRR | F (1, 57) = 0.168 | P=0.683 | 0.254 | | | | | | | |
| | | | | CDDO-EA/33-GCR | 15 | 2.6 | Yes | | | | | | | | | | | | |
| | Block 5 Total Reversals Small Separation | | 5B | Veh/Sham | 14 | 0.8571 | No | 2-way ANOVA | IRR x Diet | F (1, 57) = 0.644 | P=0.425 | 0.958 | | | | | | | |
| | | | | Veh/33-GCR | 16 | 0.5 | No | | Diet | F (1, 57) = 8.877 | **P=0.004** | 13.19 | partial ω^2 = 0.11 (Medium), 95% CI [0.01, 0.30] | Yes | Bonferroni | Veh/33-GCR vs CDDO-EA/33-GCR: *e* **P=0.018** | 0.998 (Large) | 0.177, 1.79 | Yes |
| | | | | CDDO-EA/Sham | 16 | 1.375 | Yes | | IRR | F (1, 57) = 0.487 | P=0.488 | 0.488 | | | | | | | |
| | | | | CDDO-EA/33-GCR | 15 | 1.4 | No | | | | | | | | | | | | |
| Elevated Plus Maze | Time Spent in Open Arms | | 6A | Veh/Sham | 14 | 64.48 | Yes | 2-way ANOVA | IRR x Diet | F (1, 56) = 0.948 | P=0.334 | 1.634 | | | | NA | | | |
| | | | | Veh/33-GCR | 16 | 48.89 | Yes | | Diet | F (1, 56) = 0.039 | P=0.842 | 0.068 | | | | | | | |
| | | | | CDDO-EA/Sham | 16 | 58.45 | Yes | | IRR | F (1, 56) = 1.057 | P=0.308 | 1.819 | | | | | | | |
| | | | | CDDO-EA/33-GCR | 14 | 58.03 | Yes | | | | | | | | | | | | |
| | Visit to Open Arms | | 6B | Veh/Sham | 14 | 11.5 | Yes | 2-way ANOVA | IRR x Diet | F (1, 56) = 0.09462 | P=0.759 | 0.1622 | | | | NA | | | |
| | | | | Veh/33-GCR | 16 | 11.56 | No | | Diet | F (1, 56) = 2.029 | P=0.159 | 3.478 | | | | | | | |
| | | | | CDDO-EA/Sham | 16 | 9.188 | Yes | | IRR | F (1, 56) = 0.1256 | P=0.724 | 0.2154 | | | | | | | |
| | | | | CDDO-EA/33-GCR | 14 | 10.07 | Yes | | | | | | | | | | | | |
| | Exploration Ratio | | 6C | Veh/Sham | 14 | -0.4665 | Yes | 2-way ANOVA | IRR x Diet | F (1, 56) = 1.159 | P=0.286 | 1.992 | | | | NA | | | |
| | | | | Veh/33-GCR | 16 | -0.5988 | Yes | | Diet | F (1, 56) = 0.0617 | P=0.804 | 0.106 | | | | | | | |
| | | | | CDDO-EA/Sham | 16 | -0.5202 | No | | IRR | F (1, 56) = 0.9308 | P=0.338 | 1.6 | | | | | | | |
| | | | | CDDO-EA/33-GCR | 14 | -0.5129 | Yes | | | | | | | | | | | | |
| | Total Distance Moved | | 6D | Veh/Sham | 14 | 596.2 | Yes | 2-way ANOVA | IRR x Diet | F (1, 56) = 0.06432 | P=0.800 | 0.1116 | | | | NA | | | |
| | | | | Veh/33-GCR | 16 | 573.3 | Yes | | Diet | F (1, 56) = 1.377 | P=0.245 | 2.388 | | | | | | | |
| | | | | CDDO-EA/Sham | 16 | 554.9 | Yes | | IRR | F (1, 56) = 0.2909 | P=0.591 | 0.5046 | | | | | | | |
| | | | | CDDO-EA/33-GCR | 14 | 546.6 | Yes | | | | | | | | | | | | |
| Stimulus-Response Acquisition | Total Beam Breaks | | 7A | Veh/Sham | 12 | 282.9 | Yes | 2-way ANOVA | IRR x Diet | F (1, 49) = 0.002 | P=0.958 | 0.004 | | | | | | | |
| | | | | Veh/33-GCR | 13 | 196.8 | Yes | | Diet | F (1, 49) = 1.447 | P=0.234 | 2.386 | | | Bonferroni | Veh/Sham vs Veh/33-GCR: *a#* P=0.063 | 1.21 (Large) | -2.22, -0.216 | Yes |
| | | | | CDDO-EA/Sham | 14 | 313.7 | Yes | | IRR | F (1, 49) = 10.01 | **P=0.002** | 16.51 | partial ω^2 = 0.11 (Medium), 95% CI [0.01, 0.30] | **Yes** | | CDDO-EA/Sham vs CDDO-EA/33-GCR: *f#* P=0.055 | 0.721 (Medium) | -1.52, 0.0495 | No |
| | | | | CDDO-EA/33-GCR | 14 | 230.4 | Yes | | | | | | | | | | | | |
| | Days to Completion | | 7C | Veh/Sham | 12 | 12.83 | Yes | 2-way ANOVA | IRR x Diet | F (1, 49) = 7.457 | **P=0.0088** | 6.649 | partial ω^2 = 0.11 (Medium), 95% CI [0.00, 0.30] | Yes | Tukey | Veh/Sham vs Veh/33-GCR: *a'''* **P<0.0001** | 1.67 (Large) | 0.632, 2.8 | Yes |
| | | | | Veh/33-GCR | 13 | 24.38 | Yes | | Diet | F (1, 49) = 35.79 | **P<0.0001** | 31.92 | partial ω^2 = 0.40 (Large), 95% CI [0.18, 0.59] | Yes | | Veh/33-GCR vs. CDDO-EA/Sham: *d'''* **P<0.0001** | 3.09 (Large) | -4.73, -1.77 | Yes |
| | | | | CDDO-EA/Sham | 14 | 7.571 | No | | IRR | F (1, 49) = 19.43 | **P<0.0001** | 17.33 | partial ω^2 = 0.26 (Large), 95% CI [0.07, 0.47] | Yes | | Veh/33-GCR vs. CDDO-EA /33-GCR: *e'''* **P<0.0001** | 2.28 (Large) | -3.63, -1.12 | Yes |
| | | | | CDDO-EA/33-GCR | 14 | 10.29 | No | | | | | | | | | | | | |
| | Session Length | Day 1 | 7D | Veh/Sham | 12 | 1060 | Yes | Repeated Measures 2-way ANOVA | Day x Treatment | F (6, 98) = 1.629 | P=0.147 | 2.695 | | | Tukey | | | | |
| | | | | Veh/33-GCR | 13 | 1454 | No | | Day | F (1.819, 89.15) = 56.32 | **P<0.0001** | 31.06 | partial ω^2 = 0.34 (Large), 95% CI [0.19, 0.48] | Yes | | | | | |
| | | | | CDDO-EA/Sham | 14 | 969.4 | Yes | | Treatment | F (3, 49) = 4.892 | **P=0.004** | 9.056 | partial ω^2 = 0.12 (Medium), 95% CI [0.01, 0.24] | Yes | | Veh/33-GCR vs CDDO-EA/Sham on Day1: *d'* **P=0.007** | 1.37 (Large) | -2.32, -0.45 | Yes |
| | | | | CDDO-EA/33-GCR | 14 | 1223 | Yes | | Subject | F (49, 98) = 2.238 | **P=0.0004** | 30.24 | partial ω^2 = 0.19 (Large), 95% CI [0.00, 1.00] | No | | | | | |
| | | Day 4 | | Veh/Sham | 12 | 715.7 | Yes | | | | | | | | | Veh/Sham vs Veh/33-GCR on Day 4: *a* **P=0.022** | 1.25 (Large) | 0.456, 1.98 | Yes |
| | | | | Veh/33-GCR | 13 | 1109 | Yes | | | | | | | | | | | | |
| | | | | CDDO-EA/Sham | 14 | 717.8 | No | | | | | | | | | Veh/33-GCR vs CDDO-EA/Sham on Day 4: *d* **P=0.046** | 1.08 (Large) | -1.97, 0.0495 | No |
| | | | | CDDO-EA/33-GCR | 14 | 797.6 | No | | | | | | | | | | | | |
| | | Final Day | | Veh/Sham | 12 | 548.4 | Yes | | | | | | | | | All comparisons Last Day P>0.05 | | | |
| | | | | Veh/33-GCR | 13 | 674.5 | Yes | | | | | | | | | | | | |
| | | | | CDDO-EA/Sham | 14 | 604 | Yes | | | | | | | | | | | | |
| | | | | CDDO-EA/33-GCR | 14 | 610.6 | Yes | | | | | | | | | | | | |
| | Correct Touch Latency | Day 1 | 7E | Veh/Sham | 12 | 7.883 | No | Mixed-Effects Repeated Measures 2-way ANOVA | | | | | | | Tukey | All comparisons Day 1: P>0.05 | | | |
| | | | | Veh/33-GCR | 13 | 14.55 | No | | | | | | | | | | | | |
| | | | | CDDO-EA/Sham | 14 | 9.399 | No | | | | | | | | | | | | |
| | | | | CDDO-EA/33-GCR | 13 | 9.751 | Yes | | | | | | | | | | | | |
| | | Day 4 | | Veh/Sham | 12 | 6.54 | No | | Day x Treatment | F (6, 97) = 0.863 | P=0.524 | NA | | | | Veh/33-GCR vs CDDO-EA/33-GCR Day 4: *e* **P=0.048** | 1.11 (Large) | -1.91, -0.335 | Yes |
| | | | | Veh/33-GCR | 13 | 10.51 | Yes | | Day | F (1.868, 90.62) = 19.64 | **P<0.0001** | NA | | | | | | | |
| | | | | CDDO-EA/Sham | 14 | 5.653 | No | | Treatment | F (3, 49) = 2.844 | **P=0.047** | NA | | | | | | | |
| | | | | CDDO-EA/33-GCR | 14 | 5.573 | No | | | | | | | | | | | | |
| | | Last Day | | Veh/Sham | 12 | 3.898 | Yes | | | | | | | | | Veh/33-GCR vs CDDO-EA/Sham Last Day: *d* ***P=0.047*** | 1.11 (Large) | -1.88, -0.299 | Yes |
| | | | | Veh/33-GCR | 13 | 5.195 | Yes | | | | | | | | | | | | |
| | | | | CDDO-EA/Sham | 14 | 3.637 | Yes | | | | | | | | | | | | |
| | | | | CDDO-EA/33-GCR | 14 | 4.334 | Yes | | | | | | | | | | | | |
| | | Day 1 | | Veh/Sham | 12 | 3.488 | No | | | | | | | | | Veh/Sham vs Veh/33-GCR Day 1: *a* **P=0.020** | 1.27 (Large) | 0.284, 2.02 | Yes |
| | | | | Veh/33-GCR | 13 | 7.574 | Yes | | | | | | | | | | | | |
| | | | | CDDO-EA/Sham | 14 | 2.968 | No | | | | | | | | | | 1.55 (Large) | -2.24, -0.743 | Yes |
| | | | | CDDO-EA/33-GCR | 13 | 4.888 | No | | | | | | | | | Veh/33-GCR vs CDDO-EA/Sham Day 1: *d'* **P=0.006** | | | |

**Supplementary Table 1. Statistics for Yun and Kiffer et al.**

| Experiment | Measure | Test Phase | Figure Panel | Group | Sample Size (n) | Mean (predicted or rank) or Median | Gaussian (vs. LogNormal) | Test Statistic | Main Effect or Interaction (bold text: P<0.05) | Dist.-Value (Dfn, Dfd) | Main Effect P-Value (bold text: P<0.05) | % of Total Variation | **Effect Size:** **Partial ω^2 when two-way ANOVA p<0.05.** ≥0.01 small; ≥0.06 medium; ≥0.13 large; **Partial eta square, when Kruskal-Wallis p<0.05.** ≤0.01 small, ≤0.06 medium, and [illegible] | Can effect size be considered different than zero? | Post-hoc Test Correction | **Group Difference P-Value (bold text:** P<0.05). Lower case italicized letter (e.g. *a, b, c,* etc) indicates specific group differences. Apostrophe indicates gradations of P values. Post-hoc values of 0.05<P<0.08 are only considered notable (and indicated below and on graph by #) when two additional conditions are met: if effect size is medium or large and if post-hoc 95% CI does not include zero | **Effect Size: Cohen's d** for parametric multiple comparison and **Cliff's delta** for non-parametric mulitple comparison (Lenhard, W. & Lenhard, A. (2016), Ho et al., (2019); determined only for P<0.08, | 95% CI | Can post-hoc effect size be considered different than zero? |
|---|---|---|---|---|---|---|---|---|---|---|---|---|---|---|---|---|---|---|---|
| | Reward Collection Latency | Day 4 | 7F | Veh/Sham | 12 | 1.747 | Yes | | Day x Treatment | F (6, 97) = 4.430 | **P=0.0005** | NA | | | Tukey | | | | |
| | | | | Veh/33-GCR | 13 | 2.794 | No | Mixed-Effects Repeated Measures 2-way ANOVA | Day | F (1.089, 52.76) = 44.38 | **P<0.0001** | NA | | | | Veh/Sham vs Veh/33-GCR Day 4: *a* **P=0.020** | 1.27 (Large) | 0.493, 1.9 | Yes |
| | | | | CDDO-EA/Sham | 14 | 1.555 | Yes | | Treatment | F (3, 49) = 8.322 | **P=0.0001** | NA | | | | Veh/33-GCR vs CDDO-EA/Sham Day 4: *d'* **P=0.004** | 1.69 (Large) | -2.34, -0.981 | Yes |
| | | | | CDDO-EA/33-GCR | 14 | 1.67 | No | | | | | | | | | Veh/33-GCR vs CDDO-EA/33-GCR Day 4: *e* **P=0.010** | 1.41 (Large) | -2.1, -0.577 | Yes |
| | | Last Day | | Veh/Sham | 12 | 1.78 | Yes | | | | | | | | | | | | |
| | | | | Veh/33-GCR | 13 | 1.826 | Yes | | | | | | | | | All comparisons Last Day: P>0.05 | | | |
| | | | | CDDO-EA/Sham | 14 | 1.653 | Yes | | | | | | | | | | | | |
| | | | | CDDO-EA/33-GCR | 14 | 1.664 | Yes | | | | | | | | | | | | |
| | Correct touches | Day 1 | 7G | Veh/Sham | 12 | 28.42 | Yes | | | | | | | | | | | | |
| | | | | Veh/33-GCR | 13 | 26.69 | Yes | | | | | | | | | | | | |
| | | | | CDDO-EA/Sham | 14 | 30 | Yes | | | | | | | | | | | | |
| | | | | CDDO-EA/33-GCR | 14 | 26.57 | Yes | | | | | | | | | | | | |
| | | Day 4 | | Veh/Sham | 12 | 30 | Yes | | Day x Treatment | F (6, 98) = 1.359 | P=0.2389 | 4.352 | | | Tukey | | | | |
| | | | | Veh/33-GCR | 13 | 28.92 | Yes | Mixed-Effects Repeated Measures 2-way ANOVA | Day | F (1.455, 71.31) =5.956 | **P=0.0087** | 6.359 | | | | | | | |
| | | | | CDDO-EA/Sham | 14 | 29 | Yes | | Treatment | F (3, 49) = 0.8894 | P=0.4532 | 1.911 | | | | | | | |
| | | | | CDDO-EA/33-GCR | 14 | 29.64 | Yes | | | | | | | | | | | | |
| | | Last Day | | Veh/Sham | 12 | 30 | Yes | | | | | | | | | | | | |
| | | | | Veh/33-GCR | 13 | 30 | Yes | | | | | | | | | | | | |
| | | | | CDDO-EA/Sham | 14 | 30 | Yes | | | | | | | | | | | | |
| | | | | CDDO-EA/33-GCR | 14 | 30 | Yes | | | | | | | | | | | | |
| Extinction Learning | Days to Completion | Extinction | 7I | Veh/Sham | 12 | 13.42 | No | | IRR x Diet | F (1, 48) = 2.003 | P=0.1634 | 3.772 | | | | | | | |
| | | | | Veh/33-GCR | 12 | 8.75 | Yes | 2-way ANOVA | Diet | F (1, 48) = 0.001 | P=0.9734 | 0.0021 | | | | NA | | | |
| | | | | CDDO-EA/Sham | 14 | 11.36 | Yes | | IRR | F (1, 48) = 3.488 | P=0.0679 | 6.567 | | | | | | | |
| | | | | CDDO-EA/33-GCR | 14 | 10.71 | No | | | | | | | | | | | | |
| | Session Length | Day 1 | 7J | Veh/Sham | 12 | 507.6 | No | | | | | | | | | Veh/Sham Day 1 vs Day 8: *1'''* **P<0.0001** | 3.55 (Large) | 2.24, 5.02 | Yes |
| | | | | Veh/33-GCR | 12 | 525.6 | Yes | | | | | | | | | Veh/Sham Day 1 vs Last Day: *1'''* **P<0.0001** | 5.18 (Large) | 3.79, 6.5 | Yes |
| | | | | CDDO-EA/Sham | 14 | 513 | Yes | | | | | | | | | | | | |
| | | | | CDDO-EA/33-GCR | 14 | 506.7 | Yes | | Day x Treatment | F (6, 96) = 0.8179 | P=0.5587 | 0.8711 | | | | Veh/GCR Day 1 vs Day 8: *2'''* **P<0.0001** | 3.05 (Large) | 1.64, 4.4 | Yes |
| | | Day 8 | | Veh/Sham | 12 | 574.9 | Yes | | Day | F (1.880, 90.23) = 196.9 | **P<0.0001** | 69.92 | partial ω^2 = 0.72(Large), 95% CI [0.60, 0.80] | Yes | | Veh/GCR Day 1 vs Last Day: *2'''* **P<0.0001** | 4.75 (Large) | 3.73, 5.69 | Yes |
| | | | | Veh/33-GCR | 12 | 583.7 | Yes | Repeated Measures 2-way ANOVA | Treatment | F (3, 48) = 1.877 | P=0.1460 | 1.175 | | | Tukey | | | | |
| | | | | CDDO-EA/Sham | 14 | 579.5 | Yes | | Subject | F (48, 96) = 1.175 | P=0.2496 | 10.01 | | | | CDDO/Sham Day 1 vs Day 8: *3'''* **P<0.0001** | 2.46 (Large) | 1.54, 3.38 | Yes |
| | | | | CDDO-EA/33-GCR | 14 | 583.9 | Yes | | | | | | | | | CDDO/Sham Day 1 vs Last Day: *3'''* **P<0.0001** | 2.91 (Large) | 1.91, 3.95 | Yes |
| | | Final Day | | Veh/Sham | 12 | 581.2 | Yes | | | | | | | | | | | | |
| | | | | Veh/33-GCR | 12 | 587.1 | Yes | | | | | | | | | CDDO/GCR Day 1 vs Day 8: *4'''* **P<0.0001** | 3.47 (Large) | 2.41, 4.84 | Yes |
| | | | | CDDO-EA/Sham | 14 | 589.2 | Yes | | | | | | | | | CDDO/GCR Day 1 vs Last Day: *4'''* **P<0.0001** | 2.46 (Large) | 1.31, 3.66 | Yes |
| | | | | CDDO-EA/33-GCR | 14 | 576.8 | Yes | | | | | | | | | | | | |
| | Total Responses | Day 1 | 7K | Veh/Sham | 12 | 15.58 | No | | Day x Treatment | F (6, 96) = 1.555 | P=0.168 | 1.798 | | | | Veh/Sham Day 1 vs Day 8: *1''* **P=0.0006** | 2.52 (Large) | -5.63, -0.645 | Yes |
| | | | | Veh/33-GCR | 12 | 15.58 | Yes | | Day | F (1.938, 93.03) = 177.4 | **P<0.0001** | 68.37 | partial ω^2 = 0.70 (Large), 95% CI [0.58, 0.79] | Yes | | Veh/Sham Day 1 vs Last Day: *1''* **P=0.0004** | 1.88 (Large) | -4.16, -0.458 | Yes |
| | | | | CDDO-EA/Sham | 14 | 17.21 | Yes | | Treatment | F (3, 48) = 0.5010 | P=0.6834 | 0.3024 | | | | | | | |
| | | | | CDDO-EA/33-GCR | 14 | 17 | Yes | | Subject | F (48, 96) = 1.044 | P=0.4206 | 9.656 | | | | Veh/GCR Day 1 vs Day 8: *2'''* **P<0.0001** | 3.26 (Large) | -4.32, -1.91 | Yes |
| | | Day 8 | | Veh/Sham | 12 | 5.667 | Yes | | | | | | | | | Veh/GCR Day 1 vs Last Day: *2'''* **P<0.0001** | 4.75 (Large) | -6.27, -3.38 | Yes |
| | | | | Veh/33-GCR | 12 | 5.333 | Yes | Repeated Measures 2-way ANOVA | | | | | | | Tukey | | | | |
| | | | | CDDO-EA/Sham | 14 | 6.071 | Yes | | | | | | | | | CDDO/Sham Day 1 vs Day 8: *3'''* **P<0.0001** | 2.92 (Large) | -3.76, -1.88 | Yes |
| | | | | CDDO-EA/33-GCR | 14 | 5.071 | Yes | | | | | | | | | CDDO/Sham Day 1 vs Last Day: *3'''* **P<0.0001** | 3.8 (Large) | -4.85, -2.55 | Yes |
| | | Final Day | | Veh/Sham | 12 | 6.917 | Yes | | | | | | | | | | | | |
| | | | | Veh/33-GCR | 12 | 4.833 | No | | | | | | | | | CDDO/GCR Day 1 vs Day 8: *4'''* **P<0.0001** | 4.13 (Large) | -5.63, -2.68 | Yes |
| | | | | CDDO-EA/Sham | 14 | 3.571 | Yes | | | | | | | | | CDDO/GCR Day 1 vs Last Day: *4'''* **P<0.0001** | 2.7 (Large) | -4.01, -1.63 | Yes |
| | | | | CDDO-EA/33-GCR | 14 | 6.214 | Yes | | | | | | | | | | | | |
| Lcomotor Activity Chambers | Distance moved | | 8A | Veh/Sham | 13 | 830.6 | Yes | | IRR x Diet | F (1, 50) = 0.1063 | P=0.7458 | 0.193 | | | | | | | |
| | | | | Veh/33-GCR | 12 | 781.8 | Yes | | Diet | F (1, 50) = 0.05484 | P=0.8158 | 0.09957 | | | | | | | |
| | | | | CDDO-EA/Sham | 14 | 788.3 | No | 2-way ANOVA | IRR | F (1, 50) = 4.737 | **P=0.0343** | 8.6 | partial ω^2 = 0.06 (Medium), 95% CI [0.00, 0.24] | No | Bonferroni | All comparisons P>0.05 | | | |
| | | | | CDDO-EA/33-GCR | 15 | 685 | Yes | | | | | | | | | | | | |
| | Mean velocity | | 8B | Veh/Sham | 13 | 26.42 | Yes | | IRR x Diet | F (1, 50) = 0.1154 | P=0.7355 | 0.2291 | | | | | | | |
| | | | | Veh/33-GCR | 12 | 26.43 | Yes | 2-way ANOVA | Diet | F (1, 50) = 0.09048 | P=0.7648 | 0.1797 | | | | NA | | | |
| | | | | CDDO-EA/Sham | 14 | 26.37 | No | | IRR | F (1, 50) = 0.1234 | P=0.7269 | 0.245 | | | | | | | |
| | | | | CDDO-EA/33-GCR | 15 | 27.15 | No | | | | | | | | | | | | |
| | Ambulatory time | | 8C | Veh/Sham | 13 | 37.43 | No | | IRR x Diet | F (1, 50) = 0.2921 | P=0.591 | 0.4944 | | | | | | | |
| | | | | Veh/33-GCR | 12 | 34.89 | Yes | | Diet | F (1, 50) = 0.7303 | P=0.396 | 1.236 | | | | | | | |
| | | | | CDDO-EA/Sham | 14 | 34.66 | No | 2-way ANOVA | IRR | F (1, 50) = 7.612 | **P=0.008** | 12.88 | partial ω2 = 0.11 (Medium), 95% CI [0.00, 0.30] | **Yes** | Bonferroni | CDDO-EA/Sham vs CDDO-EA/33-GCR: *f* **P=0.037** | 0.839 (Large) | -1.55, -0.0436 | Yes |
| | | | | CDDO-EA/33-GCR | 15 | 28.85 | Yes | | | | | | | | | | | | |
| | Ambulatory events | | 8D | Veh/Sham | 13 | 46.88 | Yes | | IRR x Diet | F (1, 50) = 0.1975 | P=0.6586 | 0.3404 | | | | | | | |
| | | | | Veh/33-GCR | 12 | 44.28 | Yes | | Diet | F (1, 50) = 0.8323 | P=0.3660 | 1.434 | | | | | | | |
| | | | | CDDO-EA/Sham | 14 | 43.86 | Yes | 2-way ANOVA | IRR | F (1, 50) = 6.606 | **P=0.0132** | 11.38 | partial ω^2 = 0.09 (Medium), 95% CI [0.00, 0.028] | No | Bonferroni | CDDO-EA/Sham vs CDDO-EA/33-GCR: *f#* P=0.062 | 0.774 (Medium) | -1.46, -0.00907 | Yes |
| | | | | CDDO-EA/33-GCR | 15 | 36.49 | Yes | | | | | | | | | | | | |
| | Rearing time | | 8E | Veh/Sham | 13 | 12.5 | Yes | | IRR x Diet | F (1, 50) = 3.636 | P=0.0623 | 6.31 | | | | | | | |
| | | | | Veh/33-GCR | 12 | 12.82 | Yes | 2-way ANOVA | Diet | F (1, 50) = 0.7987 | P=0.3758 | 1.386 | | | | NA | | | |
| | | | | CDDO-EA/Sham | 14 | 13.59 | Yes | | IRR | F (1, 50) = 2.594 | P=0.1136 | 4.501 | | | | | | | |
| | | | | CDDO-EA/33-GCR | 15 | 9.819 | Yes | | | | | | | | | | | | |
| | Rearing # | | 8F | Veh/Sham | 13 | 21.85 | Yes | | IRR x Diet | F (1, 50) = 0.5228 | P=0.4730 | 0.9439 | | | | | | | |
| | | | | Veh/33-GCR | 12 | 21.47 | No | 2-way ANOVA | Diet | F (1, 50) = 2.318 | P=0.1342 | 4.184 | | | | NA | | | |
| | | | | CDDO-EA/Sham | 14 | 20.16 | Yes | | IRR | F (1, 50) = 2.207 | P=0.1437 | 3.984 | | | | | | | |
| | | | | CDDO-EA/33-GCR | 15 | 15.08 | Yes | | | | | | | | | | | | |
| | | | | Veh/Sham (Left vs Right) | 13 | 245.8 | Yes | | Chamber | F (1, 55) = 13.60 | **P=0.0005** | 18.55 | partial ω^2 = 0.17 (Large), 95% CI [0.03, 0.37] | **Yes** | | | | | |
| | | | | Veh/Sham (Left vs Right) | 13 | 191.1 | Yes | | Diet | F (1, 55) = 0.238 | P=0.627 | 0.025 | | | | | | | |
| | | | | Veh/33-GCR (Left vs Right) | 16 | 242.8 | Yes | | IRR | F (1, 55) = 1.556 | P=0.217 | 0.162 | | | | All comparisons P>0.05 | | | |
| | | | | Veh/33-GCR (Left vs Right) | 16 | 196.1 | Yes | | Chamber x Diet | F (1, 55) = 0.155 | P=0.695 | 0.211 | | | | | | | |

Supplementary Table 1. Statistics for Yun and Kiffer et al.

| Experiment | Measure | Test Phase | Figure Panel | Group | Sample Size (n) | Mean (predicted or rank) or Median | Gaussian (vs. LogNormal) | Test Statistic | Main Effect or Interaction (bold text: P<0.05) | Dist.-Value (Dfn, Dfd) | Main Effect P-Value (bold text: P<0.05) | % of Total Variation | Effect Size: Partial ω^2 when two-way ANOVA p<0.05. ≥0.01 small; ≥0.06 medium; ≥0.13 large; Partial eta square, when Kruskal-Wallis p<0.05. ≤0.01 small, ≤0.06 medium, and | Can effect size be considered different than zero? | Post-hoc Test Correction | Group Difference P-Value (bold text: P<0.05). Lower case italicized letter (e.g. a, b, c, etc) indicates specific group differences. Apostrophe indicates gradations of P values. Post-hoc values of 0.05<P<0.08 are only considered notable (and indicated below and on graph by #) when two additional conditions are met: if effect size is medium or large and if post-hoc 95% CI does not include zero | Effect Size: Cohen's d for parametric multiple comparison and Cliff's delta for non-parametric mulitple comparison (Lenhard, W. & Lenhard, A. (2016), Ho et al., (2019); determined only for P<0.08, | 95% CI | Can post-hoc effect size be considered different than zero? |
|---|---|---|---|---|---|---|---|---|---|---|---|---|---|---|---|---|---|---|---|
| 3-Chamber Social Interaction | Habituation | | 9A | CDDO-EA/Sham (Left vs Right) | 15 | 254.8 | Yes | Repeated Measure 3-way ANOVA | Chamber x IRR | F (1, 55) = 0.032 | P=0.859 | 0.043 | | | Bonferroni | | | | |
| | | | | CDDO-EA/Sham (Left vs Right) | 15 | 190.6 | Yes | | Diet x IRR | F (1, 55) = 2.205 | P=0.143 | 0.229 | | | | | | | |
| | | | | CDDO-EA/33-GCR (Left vs Right) | 15 | 241.8 | Yes | | Chamber x Diet x IRR | F (1, 55) = 0.007 | P=0.933 | 0.01 | | | | | | | |
| | | | | CDDO-EA/33-GCR (Left vs Right) | 15 | 180.4 | Yes | | | | | | | | | | | | |
| | Sociability | | 9B | Veh/Sham (Nov. Mouse 1) | 13 | 321.9 | Yes | Repeated Measure 3-way ANOVA | Chamber | F (1, 55) = 86.05 | **P<0.0001** | 58.16 | partial ω^2 = 0.58 (Large), 95% CI [0.39, 0.72] | **Yes** | Bonferroni | Veh/Sham Stranger 1 vs empty: *1'''* **P<0.0001** | *2.43 (Large)* | -3.22, -1.66 | Yes |
| | | | | Veh/Sham (Nov. Object) | 13 | 157.2 | Yes | | Diet | F (1, 55) = 0.237 | P=0.628 | 0.0140 | | | | | | | |
| | | | | Veh/33-GCR (Nov. Mouse 1) | 16 | 292.6 | Yes | | IRR | F (1, 55) = 0.381 | P=0.539 | 0.0230 | | | | Veh/33-GCR Stranger 1 vs empty: *2''* **P=0.005** | *1.37 (Large)* | -2.07, -0.699 | Yes |
| | | | | Veh/33-GCR (Nov. Object) | 16 | 193.9 | Yes | | Chamber x Diet | F (1, 55) = 0.497 | P=0.483 | 0.3360 | | | | | | | |
| | | | | CDDO-EA/Sham (Nov. Mouse 1) | 15 | 314.6 | Yes | | Chamber x IRR | F (1, 55) = 0.580 | P=0.449 | 0.3920 | | | | CDDO-EA/Sham Stranger 1 vs empty: *3'''* **P<0.0001** | *3.54 (Large)* | -4.7, -2.46 | Yes |
| | | | | CDDO-EA/Sham (Nov. Object) | 15 | 170.8 | Yes | | Diet x IRR | F (1, 55) = 0.035 | P=0.851 | 0.0020 | | | | | | | |
| | | | | CDDO-EA/33-GCR (Nov. Mouse 1) | 15 | 326.1 | Yes | | Chamber x Diet x IRR | F (1, 55) = 1.917 | P=0.171 | 1.2960 | | | | CDDO-EA/33-GCR Stranger 1 vs empty: *4'''* **P<0.0001** | *2.69 (Large)* | -3.5, -1.91 | Yes |
| | | | | CDDO-EA/33-GCR (Nov. Object) | 15 | 163.2 | Yes | | | | | | | | | | | | |
| | Preference for Social Novelty | | 9C | Veh/Sham (Nov. Mouse 1) | 12 | 191.6 | Yes | Repeated Measure 3-way ANOVA | Stranger Sniff Zone | F (1, 52) = 0.091 | P=0.763 | 0.1399 | | | Bonferroni | All comparisons P>0.05 | | | |
| | | | | Veh/Sham (Nov. Mouse 2) | 12 | 182.1 | Yes | | Diet | F (1, 52) = 4.157 | **P=0.046** | 1.401 | partial ω^2 = 0.01 (Small), 95% CI [0.00, 1.00] | **Yes** | | | | | |
| | | | | Veh/33-GCR (Nov. Mouse 1) | 14 | 197.6 | Yes | | IRR | F (1, 52) = 0.275 | P=0.602 | 0.0927 | | | | | | | |
| | | | | Veh/33-GCR (Nov. Mouse 2) | 14 | 209.5 | Yes | | Stranger Sniff Zone x Diet | F (1, 52) = 0.124 | P=0.726 | 0.1902 | | | | | | | |
| | | | | CDDO-EA/Sham (Nov. Mouse 1) | 15 | 219.1 | Yes | | Stranger Sniff Zone x IRR | F (1, 52) = 0.023 | P=0.879 | 0.0356 | | | | | | | |
| | | | | CDDO-EA/Sham (Nov. Mouse 2) | 15 | 221.6 | Yes | | Diet x IRR | F (1, 52) = 0.956 | P=0.332 | 0.3225 | | | | | | | |
| | | | | CDDO-EA/33-GCR (Nov. Mouse 1) | 15 | 232 | Yes | | Stranger Sniff Zone x Diet x IRR | F (1, 52) = 0.366 | P=0.547 | 0.5623 | | | | | | | |
| | | | | CDDO-EA/33-GCR (Nov. Mouse 2) | 15 | 198.6 | No | | | | | | | | | | | | |
| Open field test | Movement distance (Day 1) | | 10A | Veh/Sham | 13 | 3827 | Yes | 2-way ANOVA | IRR x Diet | F (1, 47) = 1.919 | P=0.1725 | 3.449 | | | Bonferroni | CDDO/Sham vs CDDO/33-GCR: *f* **P=0.030** | 0.874 (Large) | -1.64, -0.0333 | **Yes** |
| | | | | Veh/33-GCR | 12 | 3684 | Yes | | Diet | F (1, 47) = 2.866 | P=0.0971 | 5.151 | | | | | | | |
| | | | | CDDO-EA/Sham | 11 | 4394 | Yes | | IRR | F (1, 47) = 4.673 | **P=0.0358** | 8.399 | partial ω^2 = 0.07 (Medium), 95% CI [0.00, 0.25] | No | | | | | |
| | | | | CDDO-EA/33-GCR | 15 | 3741 | Yes | | | | | | | | | | | | |
| | Center time (Day 1) | | 10B | Veh/Sham | 13 | 65.72 | No | 2-way ANOVA | IRR x Diet | F (1, 47) = 0.3124 | P=0.5788 | 0.5854 | | | Bonferroni | CDDO/Sham vs CDDO/33-GCR: P=0.077 | 0.799 (Medium) | -1.53, 0.12 | No |
| | | | | Veh/33-GCR | 12 | 39.24 | Yes | | Diet | F (1, 47) = 0.3041 | P=0.5839 | 0.5698 | | | | | | | |
| | | | | CDDO-EA/Sham | 11 | 72.65 | Yes | | IRR | F (1, 47) = 5.952 | **P=0.0185** | 11.15 | partial ω^2 = 0.09 (Medium), 95% CI [0.00, 0.28] | No | | | | | |
| | | | | CDDO-EA/33-GCR | 15 | 44.94 | Yes | | | | | | | | | | | | |
| | Exploration Ratio (Day 1) | | 10C | Veh/Sham | 13 | -0.5973 | Yes | 2-way ANOVA | IRR x Diet | F (1, 47) = 0.3885 | P=0.5361 | 0.727 | | | Bonferroni | CDDO/Sham vs CDDO/33-GCR: P=0.070 | 0.765 (Medium) | -1.63, 0.118 | No |
| | | | | Veh/33-GCR | 12 | -0.7588 | No | | Diet | F (1, 47) = 0.5109 | P=0.4783 | 0.9559 | | | | | | | |
| | | | | CDDO-EA/Sham | 11 | -0.5345 | Yes | | IRR | F (1, 47) = 5.831 | **P=0.0197** | 10.91 | partial ω^2 = 0.09 (Medium), 95% CI [0.00, 0.27] | No | | | | | |
| | | | | CDDO-EA/33-GCR | 15 | -0.7161 | Yes | | | | | | | | | | | | |
| | Movement distance (Day 2) | | 10D | Veh/Sham | 13 | 3077 | Yes | 2-way ANOVA | IRR x Diet | F (1, 47) = 0.7256 | P=0.3986 | 1.503 | | | | NA | | | |
| | | | | Veh/33-GCR | 12 | 3212 | Yes | | Diet | F (1, 47) = 0.6110 | P=0.4383 | 1.266 | | | | | | | |
| | | | | CDDO-EA/Sham | 11 | 3345 | Yes | | IRR | F (1, 47) = 0.0008680 | P=0.9766 | 0.001 | | | | | | | |
| | | | | CDDO-EA/33-GCR | 15 | 3200 | Yes | | | | | | | | | | | | |
| | Center time (Day 2) | | 10E | Veh/Sham | 13 | 28.88 | Yes | 2-way ANOVA | IRR x Diet | F (1, 47) = 4.130 | **P=0.0478** | 6.778 | partial ω^2 = 0.06 (Medium), 95% CI [0.00, 0.23] | **Yes** | Tukey | Veh/Sham vs CDDO-EA/Sham: *b* **P=0.026** | 1 (Large) | 0.178, 1.84 | **Yes** |
| | | | | Veh/33-GCR | 12 | 25.66 | Yes | | Diet | F (1, 47) = 4.932 | **P=0.0312** | 8.096 | partial ω^2 = 0.07 (Medium), 95% CI [0.00, 0.26] | No | | Veh/33-GCR vs CDDO-EA/Sham: *d* **P=0.013** | 1.04 (Large) | 0.117, 1.82 | **Yes** |
| | | | | CDDO-EA/Sham | 11 | 58.92 | No | | IRR | F (1, 47) = 6.191 | **P=0.0164** | 10.16 | partial ω^2 = 0.09 (Medium), 95% CI [0.00, 0.29] | | | CDDO-EA/Sham vs CDDO-EA/33-GCR: *f* **P=0.012** | 1.09 (Large) | -1.96, -0.258 | **Yes** |
| | | | | CDDO-EA/33-GCR | 15 | 26.99 | No | | | | | | | | | | | | |
| | OF Exploration Index (Day 2) | | 10F | Veh/Sham | 13 | -0.8457 | No | 2-way ANOVA | IRR x Diet | F (1, 47) = 4.958 | **P=0.0308** | 7.973 | partial ω^2 = 0.07 (Medium), 95% CI [0.00, 0.26] | No | Tukey | Veh/Sham vs CDDO-EA/Sham: *b* **P=0.012** | 0.209 (Small) | 0.0609, 0.392 | **Yes** |
| | | | | Veh/33-GCR | 12 | -0.8545 | Yes | | Diet | F (1, 47) = 5.832 | **P=0.0197** | 9.378 | partial ω^2 = 0.09 (Medium), 95% CI [0.00, 0.28] | No | | Veh/33-GCR vs CDDO-EA/Sham: *d* **P=0.010** | 0.217 (Small) | 0.0627, 0.394 | **Yes** |
| | | | | CDDO-EA/Sham | 11 | -0.6371 | No | | IRR | F (1, 47) = 5.872 | **P=0.0193** | 9.442 | partial ω^2 = 0.09 (Medium), 95% CI [0.00, 0.29] | No | | CDDO-EA/Sham vs CDDO-EA/33-GCR: *f'* **P=0.009** | 0.209 (Small) | 0.0636, 0.39 | **Yes** |
| | | | | CDDO-EA/33-GCR | 15 | -0.8461 | Yes | | | | | | | | | | | | |
| Novel Object Recognition | Object exploration | | 11A | Veh/Sham (Familiar) | 14 | 21.03 | Yes | 3-way ANOVA | Object | F (1, 56) = 203.2 | **P<0.0001** | 46.11 | partial ω^2 = 0.51 (large), 95% CI [0.31, 0.67] | **Yes** | Bonferroni | | | | |
| | | | | Veh/Sham (Novel) | 14 | 69.96 | Yes | | Diet | F (1, 56) = 7.642 | **P=0.0077** | 4.2460 | partial ω^2 = 0.08 (Large), 95% CI [0.00, 0.25] | No | | Veh/Sham, Familiar vs Novel: *1'''* **P<0.0001** | 1.57 (Large) | 0.678, 2.56 | **Yes** |
| | | | | Veh/33-GCR (Familiar) | 16 | 18.39 | Yes | | IRR | F (1, 56) = 4.317 | **P=0.0423** | 2.3990 | partial ω^2 = 0.05 (Samll), 95% CI [.000, 0.20] | No | | Veh/33-GCR, Familiar vs Novel: *2''* **P<0.0001** | 2.33 (Large) | 1.17, 3.47 | **Yes** |
| | | | | Veh/33-GCR (Novel) | 16 | 70.03 | Yes | | Object x Diet | F (1, 56) = 4.330 | **P=0.0420** | 0.9826 | partial ω^2 = 0.02 (Small), 95% CI [0.00, 0.14] | No | | | | | |
| | | | | CDDO-EA/Sham (Familiar) | 15 | 38.62 | Yes | | Object x IRR | F (1, 56) = 0.3584 | P=0.5518 | 0.0813 | | | | CDDO-EA/Sham, Familiar vs Novel: *3'''* **P<0.0001** | 2.21 (Large) | 1.32, 3.15 | **Yes** |
| | | | | CDDO-EA/Sham (Novel) | 15 | 112.4 | Yes | | Diet x IRR | F (1, 56) = 3.529 | P=0.0655 | 1.9610 | | | | | | | |
| | | | | CDDO-EA/33-GCR (Familiar) | 15 | 19.35 | Yes | | Object x Diet x IRR | F (1, 56) = 0.8578 | P=0.3583 | 0.1947 | | | | CDDO-EA/33-GCR, Familiar vs Novel: *4'''* **P<0.0001** | 1.96 (Large) | 1.06, 2.67 | **Yes** |
| | | | | CDDO-EA/33-GCR (Novel) | 15 | 80.52 | Yes | | | | | | | | | | | | |
| | Object discrimination ratio | | 11B | Veh/Sham | 14 | 0.5539 | Yes | 2-way ANOVA | IRR x Diet | F (1, 56) = 0.6946 | P=0.4081 | 1.182 | | | NA | | | | |
| | | | | Veh/33-GCR | 16 | 0.5783 | Yes | | Diet | F (1, 56) = 0.02332 | P=0.8792 | 0.0397 | | | | | | | |
| | | | | CDDO-EA/Sham | 15 | 0.5128 | Yes | | IRR | F (1, 56) = 2.012 | P=0.1616 | 3.424 | | | | | | | |
| | | | | CDDO-EA/33-GCR | 15 | 0.6066 | Yes | | | | | | | | | | | | |
| Marble Burying | Total number of marbles buried | | 12A | Veh/Sham | 14 | 8.5 | Yes | 2-way ANOVA | IRR x Diet | F (1, 56) = 0.060 | P=0.805 | 0.1052 | | | NA | | | | |
| | | | | Veh/33-GCR | 16 | 10.38 | Yes | | Diet | F (1, 56) = 0.350 | P=0.556 | 0.6045 | | | | | | | |
| | | | | CDDO-EA/Sham | 15 | 9.533 | Yes | | IRR | F (1, 56) = 1.627 | P=0.207 | 2.806 | | | | | | | |

**Supplementary Table 1. Statistics for Yun and Kiffer et al.**

| Experiment | Measure | Test Phase | Figure Panel | Group | Sample Size (n) | Mean (predicted or rank) or Median | Gaussian (vs. LogNormal) | Test Statistic | Main Effect or Interaction (bold text: P<0.05) | Dist.-Value (Dfn, Dfd) | Main Effect P-Value (bold text: P<0.05) | % of Total Variation | Effect Size: **Partial ω^2 when two-way ANOVA p<0.05.** ≥0.01 small; ≥0.06 medium; ≥0.13 large; **Partial eta square, when Kruskal-Wallis p<0.05.** ≤0.01 small, ≤0.06 medium, and [illegible] | Can effect size be considered different than zero? | Post-hoc Test Correction | **Group Difference P-Value (bold text:** P<0.05). Lower case italicized letter (e.g. *a*, *b*, *c*, etc) indicates specific group differences. Apostrophe indicates gradations of P values. Post-hoc values of 0.05<P<0.08 are only considered notable (and indicated below and on graph by #) when two additional conditions are met: if effect size is medium or large and if post-hoc 95% CI does not include zero | **Effect Size: Cohen's d** for parametric multiple comparison and **Cliff's delta** for non-parametric mulitple comparison (Lenhard, W. & Lenhard, A. (2016), Ho et al., (2019); determined only for P<0.08, | 95% CI | Can post-hoc effect size be considered different than zero? |
|---|---|---|---|---|---|---|---|---|---|---|---|---|---|---|---|---|---|---|---|
| [illegible] | | | | CDDO-EA/33-GCR | 15 | 10.8 | Yes | | | | | | | | | | | | |
| | | | | Veh/Sham | 14 | 1646 | Yes | | IRR x Diet | F (1, 56) = 0.03481 | P=0.852 | 0.05348 | | | | | | | |
| | | | | Veh/33-GCR | 16 | 1249 | Yes | | Diet | F (1, 56) = 2.691 | P=0.106 | 4.134 | | | | | | | |
| **Nestlet Shredding** | **Intact nestlet weight** | | **12B** | CDDO-EA/Sham | 15 | 1963 | No | 2-way ANOVA | IRR | F (1, 56) = 6.094 | **P=0.016** | 9.363 | partial ω^2 = 0.08 (Medium), 95% CI [0.00, 0.25] | No | Bonferroni | All comparisons P>0.05 | | | |
| | | | | CDDO-EA/33-GCR | 15 | 1502 | Yes | | | | | | | | | | | | |
| | | | | Veh/Sham | 11 | 954 | Yes | | IRR x Diet | F (1, 40) = 2.366 | P=0.1319 | 4.811 | | | | | | | |
| | | | | Veh/33-GCR | 11 | 692.2 | Yes | | Diet | F (1, 40) = 0.0603 | P=0.8073 | 0.1226 | | | | | | | Yes |
| | **Total DCX+ Immature Neurons** | | **13B** | CDDO-EA/Sham | 11 | 872.2 | No | 2-way ANOVA | IRR | F (1, 40) = 6.749 | **P=0.0131** | 13.73 | partial ω^2 = 0.12 (Medium), 95% CI [0.00, 0.33] | No | Bonferroni | Veh/Sham vs Veh/33-GCR, *a* **P=0.0339** | 1.42 (Large) | -2.35, -0.407 | |
| | | | | CDDO-EA/33-GCR | 11 | 805.1 | No | | | | | | | | | | | | |
| | | | | Veh/Sham | 11 | | Yes | | Bregma x Treatment | F (33, 480) = 0.7650 | P=0.8255 | 2.463 | | | | | | | |
| **Hippocampal Neurogenesis** | **DCX+ Immature Neurons** | | **13C** | Veh/33-GCR | 11 | | Yes | 2-way ANOVA | Bregma | F (11, 480) = 46.14 | **P<0.0001** | 49.52 | partial ω^2 = 0.48 (Large), 95% CI [0.41, 0.54] | **Yes** | Bonferroni | Bregma -2.02: Veh/Sham vs Veh/33-GCR, *a* **P=0.0372** | 1.21 (Large) | -1.81, -0.488 | Yes |
| | | | | CDDO-EA/Sham | 11 | | No | | Treatment | F (3, 480) = 4.036 | **P=0.0075** | 1.181 | partial ω^2 = 0.06 (Medium), 95% CI [0.02, 0.11] | **Yes** | | Bregma -3.22: CDDO/Sham vs CDDO/33-GCR, *f* **P=0.0247** | 0.885 (Large) | -1.43, -0.111 | Yes |
| | | | | CDDO-EA/33-GCR | 11 | | No | | | | | | | | | | | | |
| | | | | Veh/Sham | 11 | 1062 | Yes | | IRR x Diet | F (1, 40) = 0.01187 | P=0.9138 | 0.02748 | | | | | | | |
| | **Total DCX+ Progenitor Cells** | | **13E** | Veh/33-GCR | 11 | 1098 | Yes | 2-way ANOVA | Diet | F (1, 40) = 2.670 | P=0.1101 | 6.183 | | | NA | | | | |
| | | | | CDDO-EA/Sham | 11 | 1154 | No | | IRR | F (1, 40) = 0.5014 | P=0.4830 | 1.161 | | | | | | | |
| | | | | CDDO-EA/33-GCR | 11 | 1203 | No | | | | | | | | | | | | |
| | | | | Veh/Sham | 11 | | Yes | | Bregma x Treatment | F (33, 480) = 0.7723 | P=0.8165 | 1.961 | | | | | | | |
| | **DCX+ Progenitor Cells** | | **13F** | Veh/33-GCR | 11 | | Yes | 2-way ANOVA | Bregma | F (11, 480) = 71.85 | **P<0.0001** | 60.81 | partial ω^2 = 0.60 (Medium), 95% CI [0.53, 0.64] | **Yes** | Post-hoc on Bregma is not reported given that this is not the focus of our study. | | | | |
| | | | | CDDO-EA/Sham | 11 | | No | | Treatment | F (3, 480) = 1.297 | P=0.2747 | 0.2994 | | | | | | | |
| | | | | Veh/Sham | 14 | | | | | | | | | | | | | | |
| **Survival** | **Mouse Attrition** | | **Supp. 1A** | Veh/33-GCR | 16 | | | Log-rank (Mantel-Cox) | | $\chi^2$ = 3.6 | P=0.3 | | | | | | | | |
| | | | | CDDO-EA/Sham | 16 | | | | | | | | | | | | | | |
| | | | | CDDO-EA/33-GCR | 16 | | | | | | | | | | | | | | |
| | | | | Day1: Veh/Sham | 14 | 21 | No | | | | | | | | | | | | |
| | | | | Day1: Veh/33-GCR | 16 | 58.84 | No | | | | | | | | | | | | |
| | | | | Day1: CDDO-EA/Sham | 16 | 58.84 | No | | | | | | | | | 42 Wk: Veh/Sham vs CDDO-EA/33-GCR: *c* **P=0.010** | 0.8 (Large) | 0.467, 0.933 | Yes |
| **Alopecia Scores** | **Mouse Alopecia** | | **Supp. 1B** | Day1: CDDO-EA/33-GCR | 15 | 65.17 | No | Kruskal-Wallis | Treatment | H = 40.55 | **P<0.0001** | | Eta^2 = 0.23 (Large) | **Yes** | Dunn's Corr. | 54 Wk: Veh/Sham vs Veh/33-GCR: *a'* **P=0.007** | 0.728 (Large) | 0.357, 0.911 | Yes |
| | | | | Day2: Veh/Sham | 14 | 39.89 | No | | | | | | | | | 54 Wk: Veh/Sham vs CDDO-EA/Sham: *b* **P=0.0234** | 0.714 (Large) | 0.324, 0.895 | Yes |
| | | | | Day2: Veh/33-GCR | 16 | 84.56 | No | | | | | | | | | | | | |
| | | | | Day2: CDDO-EA/Sham | 16 | 81.3 | No | | | | | | | | | | | | |
| | | | | DAy2: CDDO-EA/33-GCR | 15 | 69.61 | No | | | | | | | | | | | | |
| | | | | Day1: Veh/Sham | 13 | 3077 | Yes | | Day | F (1, 47) = 45.71 | **P<0.0001** | 23.81 | | | | | | | |
| | | | | Day2: Veh/Sham | 13 | 750.4 | Yes | | Diet | F (1, 47) = 2.465 | P=0.123 | 2.336 | | | | | | | |
| | | | | Day1: Veh/33-GCR | 12 | 3212 | Yes | | IRR | F (1, 47) = 2.065 | P=0.157 | 1.957 | | | | | | | |
| | | | | Day2: Veh/33-GCR | 12 | 472.2 | Yes | | Day x Diet | F (1, 47) = 0.777 | P=0.382 | 0.405 | | | | | 1.65 (Large) | -2.66, -0.591 | Yes |
| | **Locomotion** | | **Supp. 2A** | Day1: CDDO-EA/Sham | 11 | 3345 | Yes | 3-way ANOVA | Day x IRR | F (1, 47) = 3.578 | P=0.064 | 1.864 | omega_p^2 = 0.25 (Large), 95% CI [0.06, 0.46] | **Yes** | Bonferroni | Veh/Sham Day 1 vs Day 2: *1''* **P=0.007** | | | |
| | | | | Day2: CDDO-EA/Sham | 11 | 1049 | Yes | | Diet x IRR | F (1, 47) = 1.985 | P=0.165 | 1.882 | | | | CDDO-EA/Sham Day 1 vs Day 2: *3'''* **P=0.0003** | 1.55 (Large) | -2.35, -0.671 | Yes |
| | | | | Day1: CDDO-EA/33-GCR | 15 | 3200 | Yes | | | | | | | | | | | | |
| | | | | Day2: CDDO-EA/33-GCR | 15 | 540.4 | Yes | | Day x Diet x IRR | F (1, 47) = 0.306 | P=0.582 | 0.1597 | | | | | | | |
| | | | | Day1: Veh/Sham | 13 | 62.39 | No | | Day | F (1, 47) = 27.88 | **P<0.0001** | 10.49 | omega_p^2 =0.12 (Medium), 95% CI [0.00, 0.32] | **Yes** | | | | | |
| | | | | Day2: Veh/Sham | 13 | 28.88 | Yes | | Diet | F (1, 47) = 2.092 | P=0.1547 | 2.535 | | | | | | | |
| | | | | Day1: Veh/33-GCR | 12 | 45.01 | Yes | | IRR | F (1, 47) = 7.797 | **P=0.0075** | 9.446 | omega_p^2 = 0.11 (Medium), 95% CI [0.00, 0.30] | **Yes** | | Veh/Sham Day 1 vs Day 2: 1' **P =0.001** | 1.15 (Large) | -1.83, -0.35 | Yes |
| **Open field** | **Center time** | | **Supp. 2B** | Day2: Veh/33-GCR | 12 | 25.66 | No | 3-way ANOVA | Day x Diet | F (1, 47) = 1.749 | P=0.1924 | 0.6583 | | | Bonferroni | | | | |
| | | | | Day1: CDDO-EA/Sham | 11 | 72.65 | Yes | | Day x IRR | F (1, 47) = 0.3850 | P=0.5379 | 0.1449 | | | | All comparisons of Sham vs IRR on the same day, P>0.05 | | | |
| | | | | Day2: CDDO-EA/Sham | 11 | 58.92 | Yes | | Diet x IRR | F (1, 47) = 1.846 | P=0.1808 | 2.236 | | | | | | | |
| | | | | Day1: CDDO-EA/33-GCR | 15 | 44.94 | Yes | | Day x Diet x IRR | F (1, 47) = 1.317 | P=0.2570 | 0.4956 | | | | | | | |
| | | | | Day2: CDDO-EA/33-GCR | 15 | 26.99 | No | | | | | | | | | | | | |
| | | | | Day1: Veh/Sham | 13 | -0.6145 | Yes | | Day | F (1, 47) = 44.90 | **P<0.0001** | 12.28 | omega_p^2 = 0.14 (Large), 95% CI [0.01, 0.35] | **Yes** | | | | | |
| | | | | Day2: Veh/Sham | 13 | -0.8457 | Yes | | Diet | F (1, 47) = 2.485 | P=0.1216 | 3.155 | | | | | | | |
| | | | | Day1: Veh/33-GCR | 12 | -0.7216 | Yes | | IRR | F (1, 47) = 6.968 | **P=0.0112** | 8.846 | omega_p^2 = 0.10 (Medium), 95% CI [0.00, 0.30] | **Yes** | | | | | |
| | **Exploration ratio** | | **Supp. 2C** | Day2: Veh/33-GCR | 12 | -0.8545 | No | 3-way ANOVA | Day x Diet | F (1, 47) = 2.182 | P=0.1463 | 0.5968 | | | Bonferroni | Veh/Sham Day 1 vs Day 2: *1'''* **P <0.0001** All comparisons of Sham vs IRR on the same day, P>0.05 | 1.38 (Large) | -2.17, -0.571 | Yes |
| | | | | Day1: CDDO-EA/Sham | 11 | -0.5345 | Yes | | Day x IRR | F (1, 47) = 0.6352 | P=0.4295 | 0.1737 | | | | | | | |
| | | | | Day2: CDDO-EA/Sham | 11 | -0.6371 | Yes | | Diet x IRR | F (1, 47) = 2.049 | P=0.1589 | 2.601 | | | | | | | |
| | | | | Day1: CDDO-EA/33-GCR | 15 | -0.7161 | Yes | | Day x Diet x IRR | F (1, 47) = 1.988 | P=0.1651 | 0.5437 | | | | | | | |
| | | | | Day2: CDDO-EA/33-GCR | 15 | -0.8461 | No | | | | | | | | | | | | |
